# Tripartite holobiont system in a vent snail broadens the concept of chemosymbiosis

**DOI:** 10.1101/2020.09.13.295170

**Authors:** Yi Yang, Jin Sun, Chong Chen, Yadong Zhou, Yi Lan, Cindy Lee Van Dover, Chunsheng Wang, Jian-Wen Qiu, Pei-Yuan Qian

## Abstract

Many animals inhabiting deep-sea vents are energetically dependent on chemosynthetic endosymbionts, but how such symbiont community interacts with host, and whether other nutritional sources are available to such animals remain unclear. To reveal the genomic basis of symbiosis in the vent snail *Alviniconcha marisindica*, we sequenced high-quality genomes of the host and gill campylobacterial endosymbionts, as well as metagenome of the gut microbiome. The gill endosymbiont has a streamlined genome for efficient chemoautotrophy, but also shows metabolic heterogeneity among populations. Inter- and intra-host variabilities among endosymbiont populations indicate the host poses low selection on gill endosymbionts. Virulence factors and genomic plasticity of the endosymbiont provide advantages for cooperating with host immunity to maintain mutualism and thriving in changing environments. In addition to endosymbiosis, the gut and its microbiome expand the holobiont’s utilisation of energy sources. Host-microbiota mutualism contributes to a highly flexible holobiont that can excel in various extreme environments.

## Introduction

Since the discovery of deep-sea hydrothermal vents in 1977, many intricate symbioses have been reported between vent-endemic animals and chemoautotrophic microbes. While some crustaceans such as the shrimp *Rimicaris exoculata* (Durand et al., 2015; Petersen et al., 2010) and the squat lobster *Shinkaia crosnieri* (Watsuji et al., 2015) rely on ectosymbionts living on their gills or their chaetae for nutrients, annelids in the family Siboglinidae (Dubilier et al., 2008) and molluscs in several families such as Vesicomyidae and Mytilidae (Dubilier et al., 2008) host endosymbiotic microbes in their bacteriocytes. Due to their intimate relationships with endosymbionts, it is generally accepted that in endosymbiosis the host relies entirely on symbionts for nutrition (Childress and Girguis, 2011; Dubilier et al., 2008).

*Alviniconcha* is a genus of chemosymbiotic provannid vent snails, with five species distributed in the Pacific Ocean and one in the Indian Ocean (Johnson et al., 2015). Among genera in the superfamily Abyssochrysoidea and those currently assigned to family Provannidae, only *Alviniconcha* and its sister genus *Ifremeria* that live in hydrothermal vents harbour endosymbionts in the gill epithelia (Beinart et al., 2019). Five species of *Alviniconcha* have been reported – four from the Pacific Ocean and one from the Indian Ocean (Johnson et al., 2015). The *Alviniconcha* species from the South Pacific are known to harbour both chemoautotrophic Gammaproteobacteria and Campylobacteria in its gills (Beinart et al., 2019), with the endosymbiont type and relative abundance varying with differences in vent fluid geochemistry (Beinart et al., 2012; Sanders et al., 2013). In contrast, *A. marisindica* from vent fields on the Central Indian Ridge (CIR) hosts a single ribotype of Campylobacterota endosymbionts (Miyazaki et al., 2020). Campylobacterota are abundant in vent habitats and also live as ectosymbionts on polychaete worms, molluscs, and crustaceans (Assié et al., 2016; Campbell et al., 2006; Goffredi, 2010; Watsuji et al., 2015), but do not commonly assume the role of intracellular symbionts. Campylobacterota are capable of oxidising sulfur, formate, and hydrogen to produce energy (Beinart et al., 2019; Miyazaki et al., 2020; Takai et al., 2005), and mostly rely on the reductive tricarboxylic acid cycle (rTCA) for carbon fixation with the exception of a bathymodiolin mussel epibiont possessing a complete Calvin–Benson–Bassham (CBB) cycle (Assié et al., 2020). Depending on the abundance of hydrogen and hydrogen sulfide, the Campylobacterota endosymbiont of *A. marisindica* is capable of shifting between these reduced compounds as its main energy source (Miyazaki et al., 2020).

Unlike siboglinid tubeworms and clam *Solemya reidi* that have lost their gut (Dubilier et al., 2008), *Alviniconcha* species retained theirs, albeit a much reduced one (Warèn and Bouchet, 1993). Dissection revealed soft biogenic substances and mineral grains inside the snail gut, indicating it is functionally active (Suzuki et al., 2005). As *Alviniconcha* hosts endosymbionts in the gill and has a previously overlooked functional gut, it serves as a good model system to tease out the complex host-microbiota interactions that are key to our understanding of the adaptations of these animals to the extreme environments in the deep ocean (McFall-Ngai et al., 2013). Here, we report comprehensive analyses of the holobiont of *Alviniconcha marisindica* from a newly discovered northern Indian Ocean population (Zhou et al., 2019). Through analysing the symbiont genome and transcriptome, we aim to unravel the chemoautotrophic metabolism of the symbionts and their machinery for interaction with host, whether such symbiont populations contain streamlined and heterogeneous genomes that may enable them to utilise diverse substrates effectively, and how genomic plasticity of such populations provide advantages for thriving in their deep-sea habitat and interacting with host to establish symbiosis. Through analysing the host genome and transcriptome, we aim to understand how the host cooperates with symbionts to maintain mutualism and how the host’s innate immunity has been remodelled to support the symbiosis. We also test the hypothesis that the gut and its microbiome are likely to provide nutrients that supplement the nutrition provided by the endosymbionts. Through dissecting the complex relations among the host, gill endosymbiont and gut microbiome, our study refine the holobiont concept in chemosymbiotic ecosystems which have enabled many animals to thrive in the extreme hydrothermal vent environments.

## Results

### Hologenome assembly and characterisation

The genome of the snail *Alviniconcha marisindica,* sequenced using a hybrid approach, is 829.61 Mb in length (N50 = 727.6 kb, genome completeness 96.5%) (Supplementary Table S1 and S2) with 21,456 predicted gene models (79.5% comparatively annotated) (Supplementary Figure S1). Comparative analyses among available lophotrochozoan genomes (n = 26; Figure 1A) reveal a lack of obvious gene family expansion in the *A. marisindica* genome. The *A. marisindica* genome encodes 737 unique gene families (6.8%; Figure 1B) when compared with the genomes of a freshwater snail, a vent-endemic chemosymbiont hosting snail in a distant clade, and a scallop. Analyses of these unique genes reveal an enrichment of genes related to oxidoreductases, hydrolases, endocytosis, transporters, and signal transduction (Supplementary Figure S2 A and B). Among the annotated genes, those involved in immune response, substrate transportation, macromolecular digestion, and absorption are highly expressed in the intestinal tissue (Supplementary Figure S2C), indicating active functioning of the *A. marisindica* gut.

**Figure 1.**
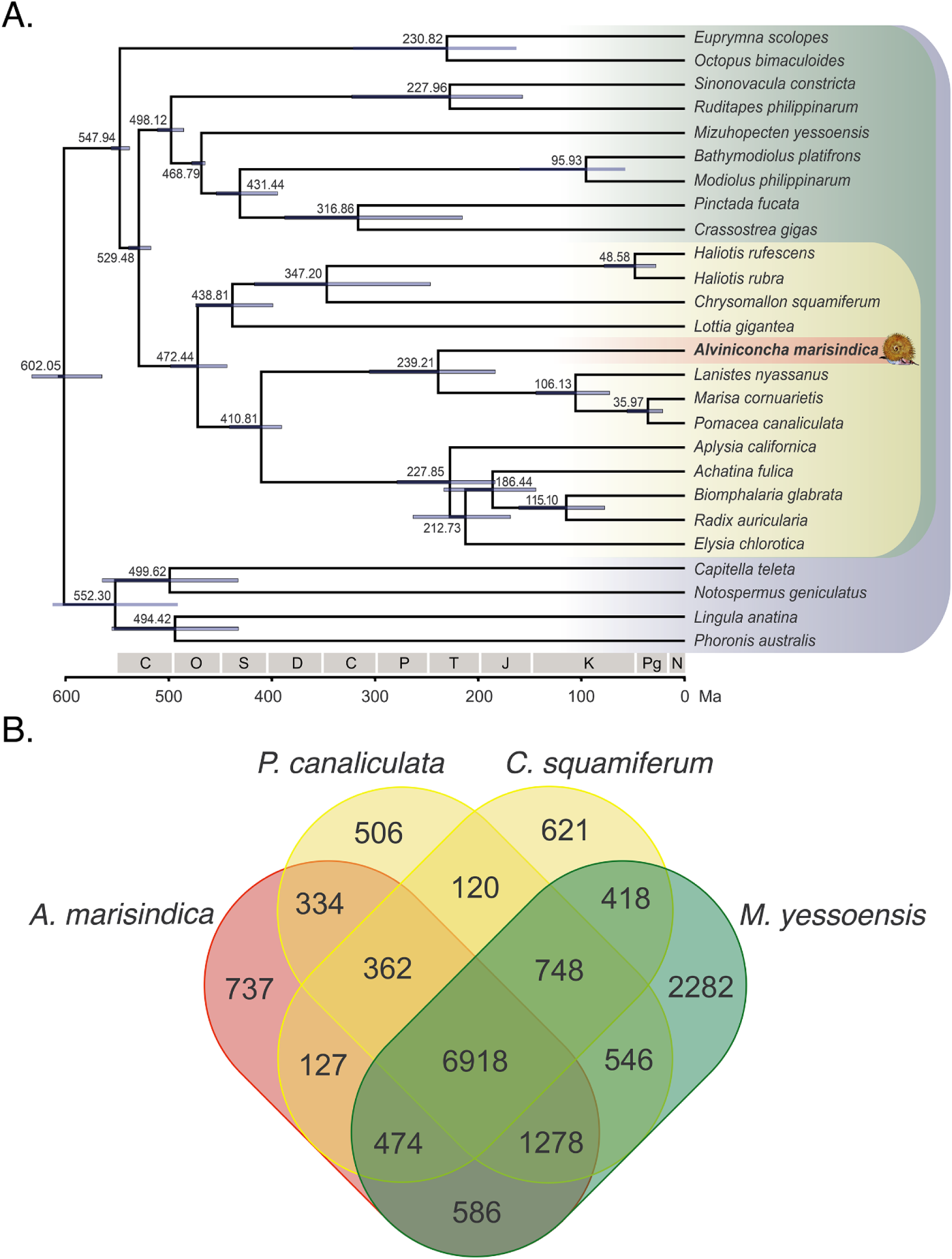
Genomic comparisons and gene family analyses across Lophotrochozoa. (**A**) Genome-based phylogeny of selected taxa showing the position of *Alviniconcha marisindica* among lophotrochozoans and divergence times among molluscan lineages. Error bars indicate 95% confidence levels. (**B**) Venn diagram depicting unique and shared gene families among four lophotrochozoan genomes.

Sequencing a bacterial 16S rRNA gene clone library from the gill tissue revealed over 99% sequence similarity among the clones, confirming the presence of a single endosymbiont phylotype in the bacteriocytes. The campylobacterotal endosymbiont genome is 1.47 Mb in length, located in two scaffolds (98.16% completeness, 0.82% contamination) with 1,429 predicted genes, among which 92.65% were successfully annotated (Figure 2A, Supplementary Figure S3A). The campylobacterotal endosymbiont genome, here named *Sulfurovum alviniconcha* CR, possesses fewer coding sequences than other available whole genomes within the phylum (Figure 3), but has the highest coding density (97.0%, Table 1) and minimal loss-of-function mutations (Figure 4A), and similar average lengths in coding regions (Supplementary Figure S4A). There are almost no flagellar or chemotaxis genes in this endosymbiont genome. When compared with its four Campylobacterota close relatives (Table 1), *Sulfurovum alviniconcha* CR lacks many cell envelope biogenesis and non-essential metabolic genes. For example, genes involved in capsular polysaccharides biogenesis (*cps*), and genes involved in partial Citrate cycle (*ace* and *DLAT*) which is one of the optional from pyruvate to aceyl-coA are missing. *Sulfurovum alviniconcha* CR and its pathogenic relatives lack many DNA-repair genes (Supplementary Figure S5) that will lead to frequent gene loss, mutation, and recombination (Kang and Blaser, 2006; Monack et al., 2004). Nevertheless, *Sulfurovum alviniconcha* CR genome contains 180 unique orthologues when compared with its four Campylobacterota close relatives (Table 1, Figure 4B), including those involved in cell wall/membrane/envelope biogenesis that modify the bacterial surface for immune evasion (e.g. *eptA*), enzymes related to oxidoreductases and translocases that promote energy production and conversion in the endosymbiont (e.g. *putA*), and extracellular proteases secretion enhancing bacterial virulence factors that associated with symbiotic interactions (e.g. *aprE* and *pulD*) (Supplementary Figure S4 B and C). In addition, *Sulfurovum alviniconcha* CR shares many virulence genes with its non-pathogenic (*Sulfurovum* species) and pathogenic (*Helicobacter* and *Campylobacter* species) Campylobacterota relatives, such as bacterial virulence factors haemolysin and MviN/MurJ, intracellular invasion CiaB, and N-linked glycosylation (NLG) with vital roles in infectivity.

**Figure 2.**
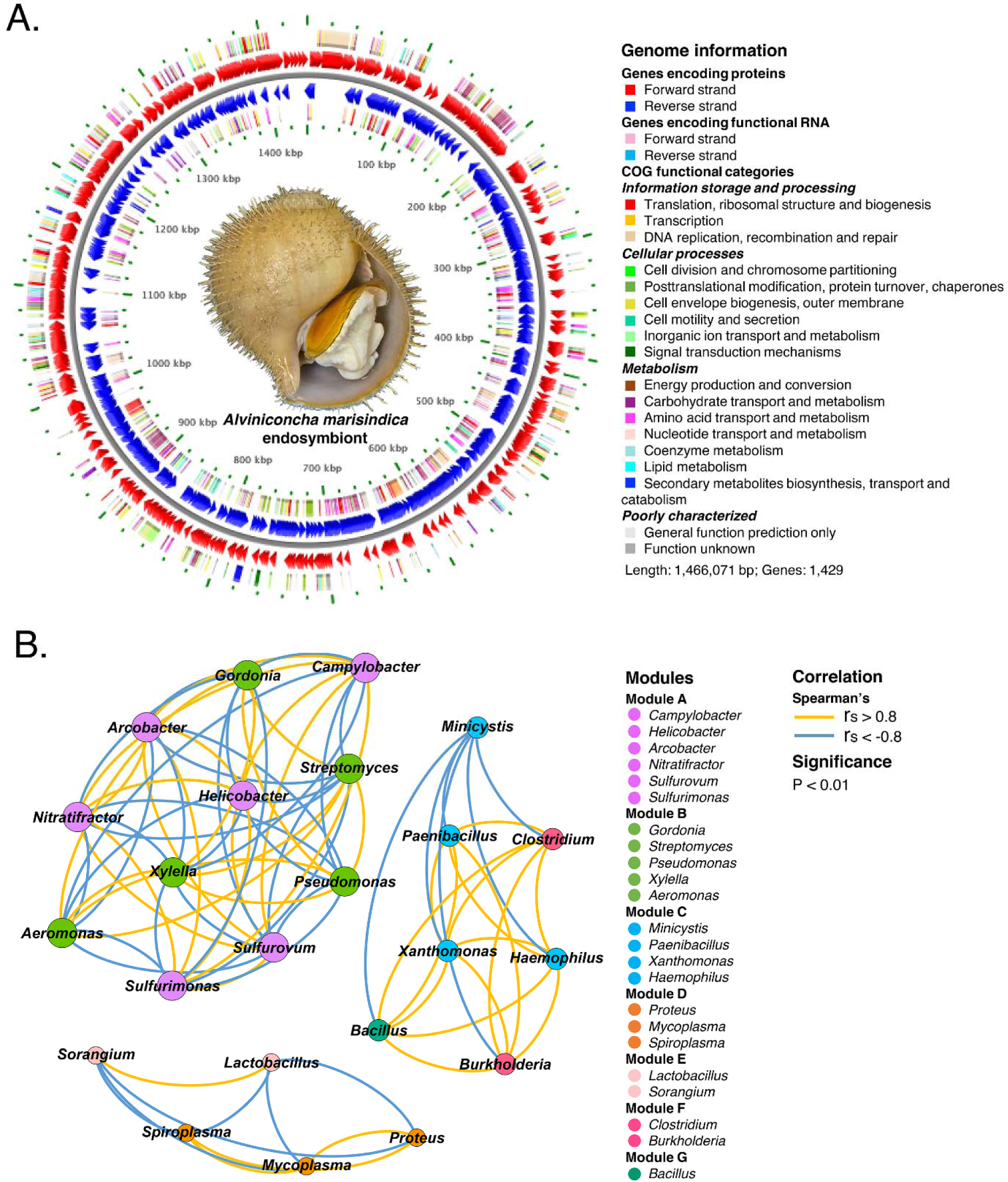
Gill endosymbionts and intestinal microbiome of *Alviniconcha marisindica* from the Wocan vent field. (**A**) Circle diagram showing an overview of genome information of the binned endosymbiont based on COG annotation. (**B**) Correlation-based network of intestinal bacteria genera (relative abundance ≥ 0.5%) from three *A. marisindica* individuals. The network analysis displays the intra-associations within each sub-community and inter-associations between sub-communities. Node size is proportional to the number of connections (i.e. degree of connectivity). Connection between nodes represents strong (Spearman correlation efficiency >0.8 (yellow) or <-0.8 (blue)) and significant (p-value <0.01) correlation. The same colour of nodes shows their highly modularised (clustered) property within the network.

**Figure 3.**
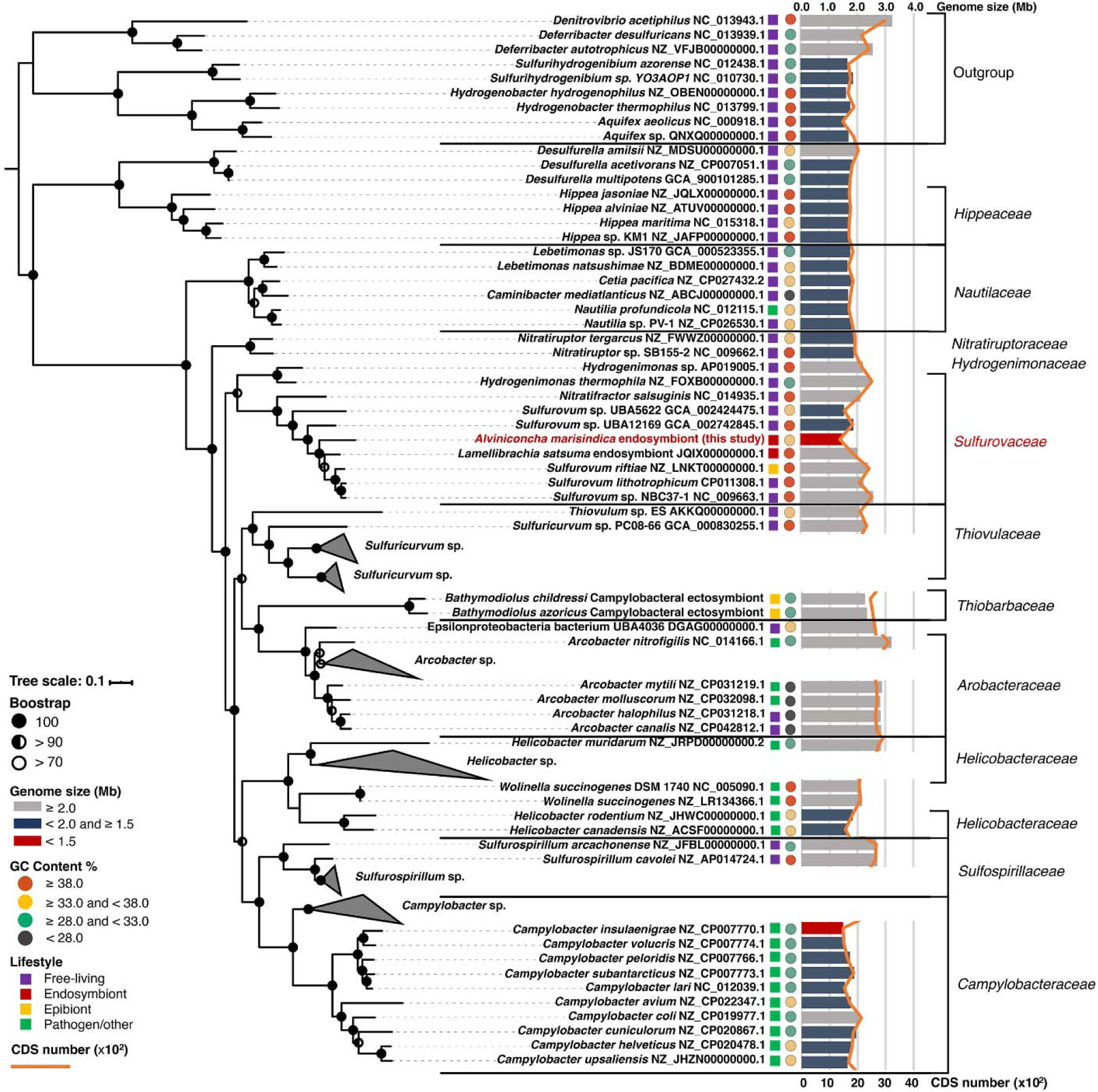
Genome-based phylogeny of campylobacterotal representatives. The position of the *Alviniconcha marisindica* endosymbiont among Campylobacterota belongs to the family Sulfurovaceae and marked in red. Nine deltaproteobacterial species are used to root the tree. Different lifestyles of the selected taxa are indicated by squares of different colours (purple: free-living, red: endosymbiont, yellow: epibiont, and green: pathogen/other). The right histogram indicates the size of each genome. The colour of a column represents the size range (grey: >2.0 Mb, dark blue: <2.0 and ≥ 5 Mb, red: <1.5 Mb). The line chart in orange indicates the number 1. of coding sequences (CDS) of each genome. Circles of different colours are used to indicate different ranges of GC content in % (red: ≥ .0, yellow: ≥ .0 and <38.0, green: ≥ 0 and <33.0, black: <28.0). The genome size of the *A. marisindica* endosymbiont is the smallest among whole genomes within the phylum, and its GC content is slightly lower than those of the four closest relatives in the same clade.

**Figure 4.**
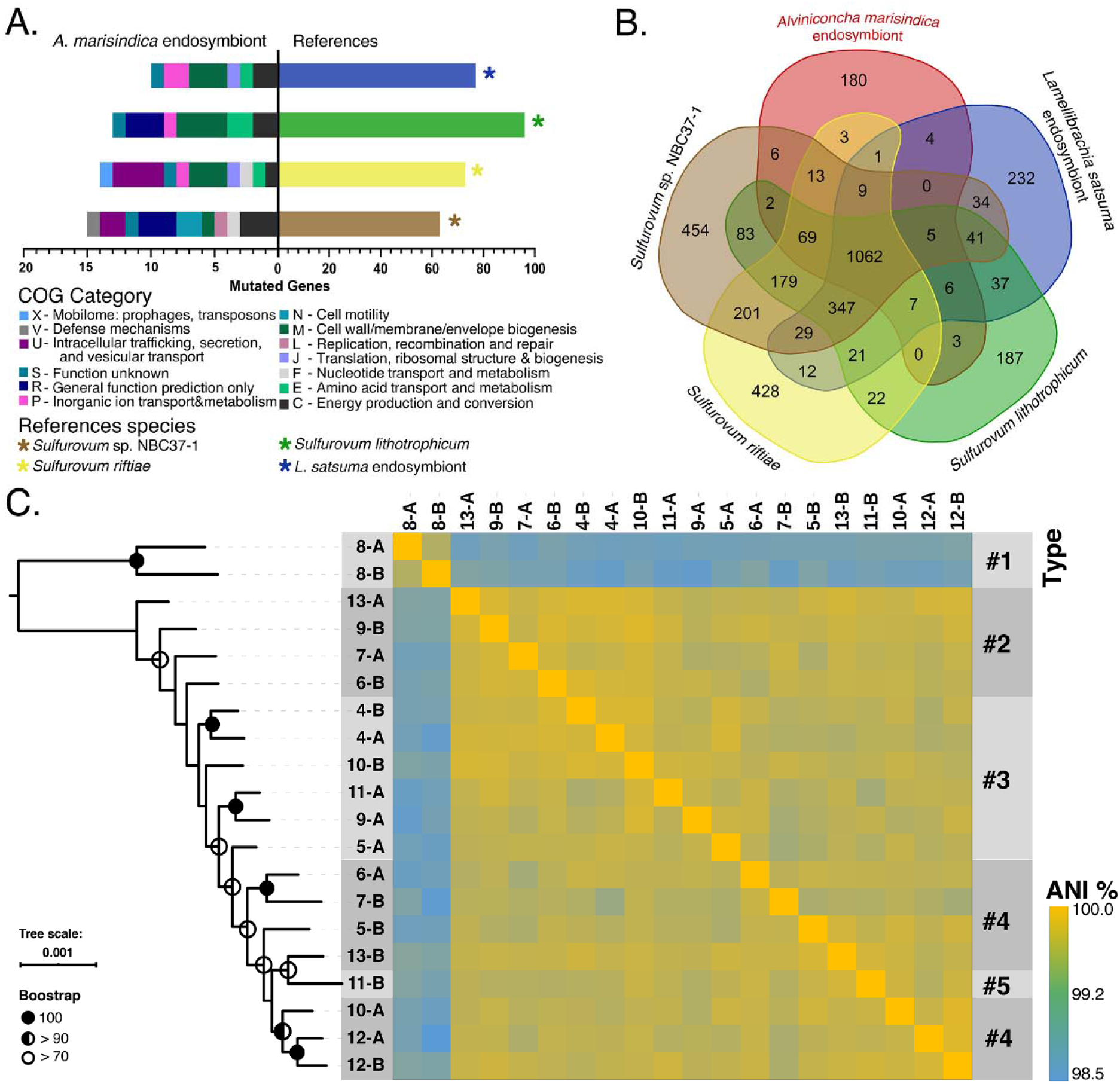
Genomic comparisons and gene family analyses of the Wocan *Alviniconcha marisindica* endosymbiont and four closely related members of Campylobacterota. (**A**) The loss-of-function genes of the *A. marisindica* endosymbiont shown in different COG categories obtained via pairwise comparison with genomes of the other four Campylobacterota members. The histogram on the left presents the result of comparing the *A. marisindica* endosymbiont with the other four campylobacterotal bacteria. The *A. marisindica* endosymbiont has significantly fewer mutated genes than do the references. (**B**) Venn diagram depicting unique and shared gene families among the five campylobacterotal genomes. (**C**) SNP-based phylogeny on the whole-genome level of 20 endosymbiotic isolates from 10 *A. marisindica* individuals showing the inter- and intra-individual relationships of *A. marisindica* endosymbionts. The genomic ANIs among these 20 isolates obtained via pairwise comparisons are shown in the heat map. The number in the name of each isolate represents the host individual, and the capital A and B represent the anterior and posterior parts of the gills, respectively.

**Table 1.** Comparison of general genomic features of *Sulfurovum alviniconcha* CR and references.

| Name | *Lamellibrachia satsuma* | *Sulfurovum lithotrophicum* | *Sulfurovum riftiae* | *Sulfurovum* sp. NBC37-1 | *Sulfurovum alviniconcha* CR |
| --- | --- | --- | --- | --- | --- |
| INSDC | JQIX000000000.1 | CP011308.1 | LNKT000000000.1 | AP009179.1 | — |
| Size (Mb) | 2.00 | 2.22 | 2.37 | 2.56 | 1.47 |
| GC (%) | 39.7 | 44.3 | 45.6 | 43.9 | 37.1 |
| Protein | 1,852 | 2,148 | 2,317 | 2,481 | 1,386 |
| tRNA | 37 | 44 | 45 | 44 | 40 |
| Gene | 2,019 | 2,227 | 2,432 | 2,583 | 1,429 |
| Percentage coding (%) | 91.7 | 96.5 | 95.3 | 96.1 | 97.0 |
| Pseudogene | 126 | 23 | 62 | 46 | — |
| Habitat | Trophosome | Sediments | Tube | Sulfide mound | Gills |
| Depth (m) | ~112 | 1,033 | 2,500 | 1,000 | 2,919 |

Genome data — *Sulfurovum alviniconcha* CR: this study; the endosymbiont of *Lamellibrachia* *satsuma*: Patra AK et al., 2016; *Sulfurovum lithotrophicum*: Inagaki F et al., 2004; *Sulfurovum* *riftiae*: Giovannelli et al., 2016; *Sulfurovum* sp. NBC37-1: Nakagawa et al., 2007.

In contrast to harbouring only one dominant phylotype of endosymbiont in the gill, the *A. marisindica* gut contains diverse microbiota. Analyses of guts from three snail individuals reveal 169 microbial genera from 38 phyla, with a different composition and relative abundance compared to those in gastropods that do not rely energetically on endosymbionts (Aronson et al., 2016; Li et al., 2019). For example, the dominant genus in gut microbes of *A. marisindica* is *Sulfurovum* (Supplementary Figure S3B) – a genus of chemoautotrophic Campylobacterota. *Sulfurovum* is a minor community in the gut of deep-sea bone-eating snail *Rubyspira osteovora* (Aronson et al., 2016) and it is rare in the gut of fresh-water polyphagous snail *Pomacea canaliculata* (Li et al., 2019). The multi-taxa associations of gut microbiome in *A. marisindica* exhibit a significant non-random co-occurrence pattern (Figure 2B), indicating the effects of the intestinal microenvironment in shaping microbial community composition. Especially, lactic acid bacteria, vital for maintaining the gut ecological balance (Koleva et al., 2014), account for at least ∼2.7% of gut microbes in *A. marisindica*, also shows that the gut microbiome is not contaminants even if they are in low density (2.61–5.57%) in *A. marisindica* (Supplementary Figure S6).

### Diversity and metabolism of the gill endosymbiont

Sequencing 23 isolates from 13 host snails (Table 2) reveals a *Sulfurovum alviniconcha* CR core-campylobacterotal genome with 1,001 shared gene families (51.8–73.0% of the predicted orthologues per isolate genome). Thiosulfate oxidation, oxygen reduction, and key reverse TCA cycle genes are present in the core-genome. Each isolate represents a subpopulation, the pan-genome of 23 subpopulations contains 2,783 orthologues, of which 1,475 are isolate-specific, and exhibit high metabolic flexibility, especially along the chemoautotrophic pathways of sulfur metabolism, hydrogen oxidation, and carbon fixation (Supplementary Figure S7). For example, the pan-genome contains hydrogen oxidation genes (*hydABCDE* gene cluster), nitrate reduction genes (*napA* and *napD*), and genes involved in hydrogen sulfide utilisation (*metZ*, catalyses the formation of L-homocysteine from O-succinyl-L-homoserine and hydrogen sulfide). Principal component and phylogenomic analyses on 941 shared single-copy orthologues of 23 isolates show that *Sulfurovum alviniconcha* CR are not clustered by their host individuals (Supplementary Figure S8). Among 23 isolates, selecting 20 isolates from the anterior and posterior gills of 10 host snails (Table 2), 20 *Sulfurovum alviniconcha* CR genomes are obtained with a total of 190 genomic average nucleotide identity (ANI) values ranging from 98.5% to 99.7% (Figure 4C). Nevertheless, these isolates belong to the same phylotype but have 28,448 single-nucleotide polymorphisms (SNPs) among them, indicating a high genetic diversity. Based on the results of genomic ANI and phylogenomic analysis of SNPs, the 20 endosymbiont isolates are classified into five types (Figure 4C) in a panmictic state among the 10 snails, showing that each snail hosts genetically diverse endosymbionts and with different types.

**Table 2.**
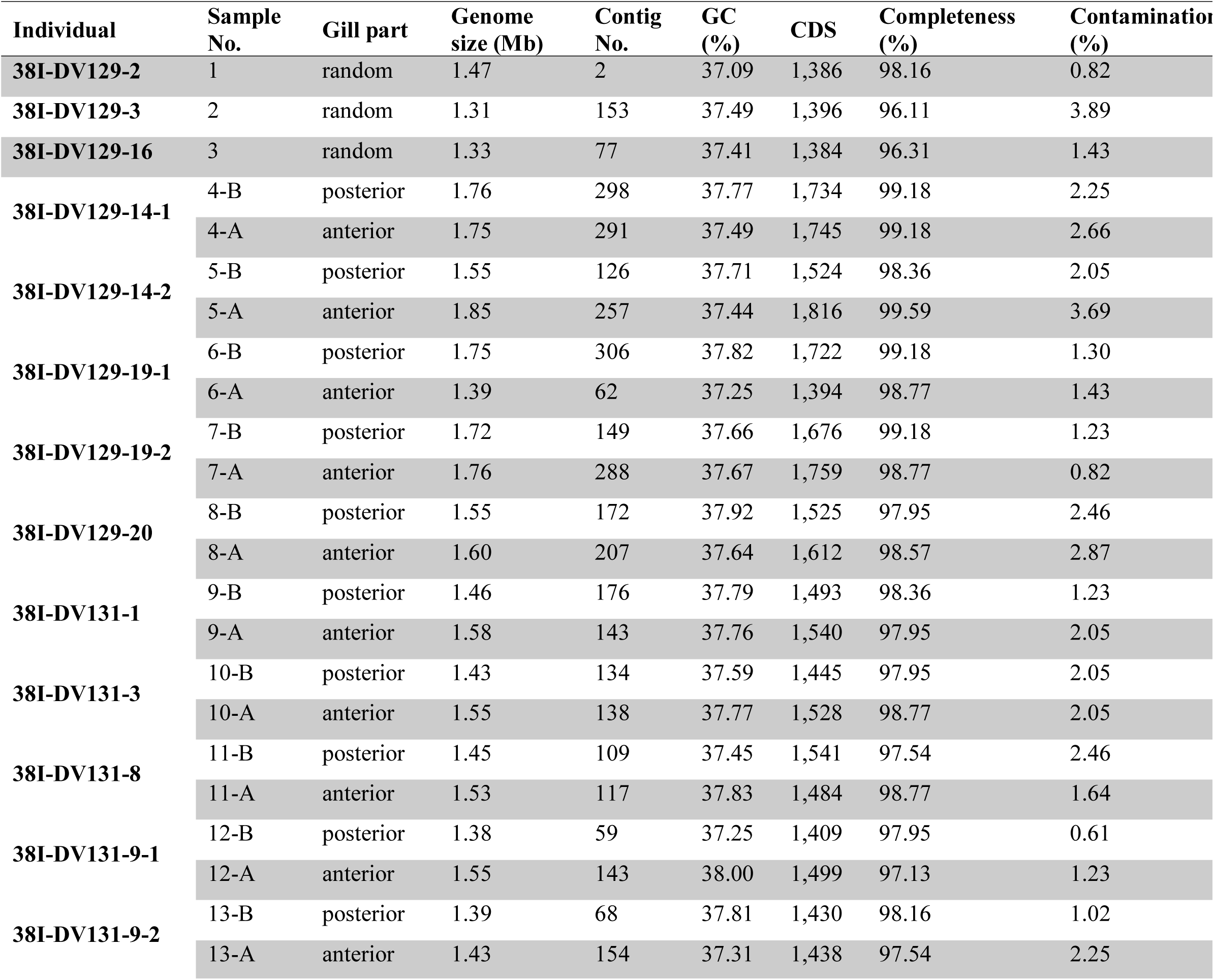
General genomic features of the binned endosymbionts of *Alviniconcha marisindica* extracted from 23 metagenome datasets of gill filaments.

Analysis of the core metabolic genes of *Sulfurovum alviniconcha* CR genome reveals its chemolithoautotroph capability, especially in anaerobic oxidation of thiosulfate (*sox* genes), but it lacks the sulfide oxidation pathway as indicated by the lack of genes in the *dsrAB* complex. The sox multi-enzyme system allows generation of energy from thiosulfate oxidation, and *soxX*-*soxY*-*soxZ*-*soxA*-*soxB* genes are highly expressed (among the top 150) in *Sulfurovum alviniconcha* CR (Supplementary Figure S9A). The absence of a sulfate/thiosulfate transporter in *Sulfurovum alviniconcha* CR genome indicates that it can only use thiosulfate from endogenous organic sulfur compounds (Figure 5). A previous study shows the gill tissue of *A. marisindica* from the Kairei hydrothermal site actively consumed environmental sulfide (Miyazaki et al., 2020), which is consistent with *sqr* (25th in transcriptome, Supplementary Figure S9A) and *cysK* genes in the *Sulfurovum alviniconcha* CR genome involving in the conversion of sulfide to polysulfides. In addition, *Sulfurovum alviniconcha* CR lacks the sulfur globule protein genes (*sgp*) for intracellular sulfur storage, indicating this endosymbiont might dependent on intracellular polysulfides for sulfur storage.

**Figure 5.**
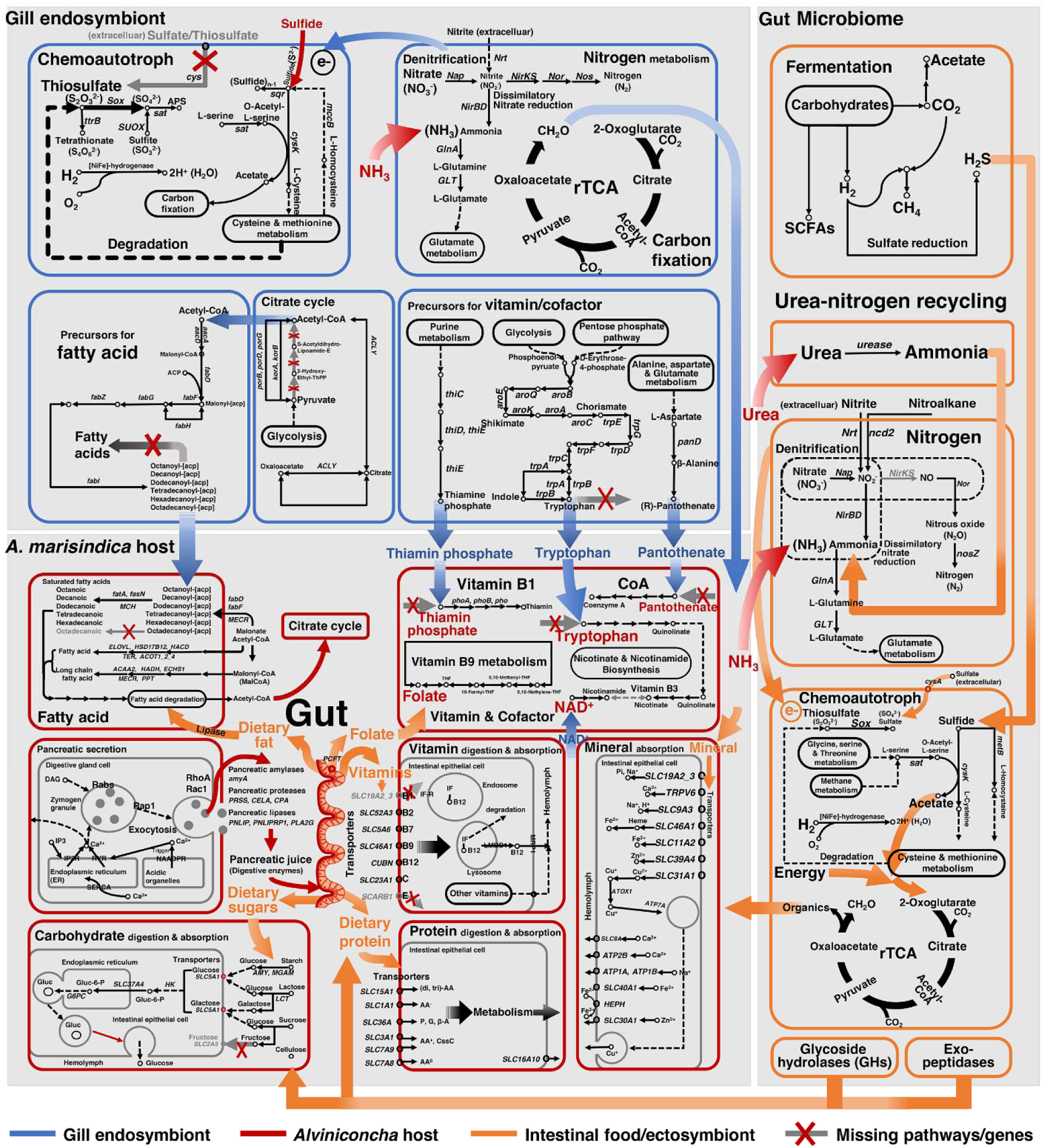
Overview of metabolic pathways of the *Alviniconcha marisindica* holobiont from the Wocan vent field. Metabolic pathways of different organisms including the gill endosymbiont, intestinal microbiome, and the *A. marisindica* host are presented in different colours (blue: gill endosymbiont, red: *A. marisindica* host, orange: intestinal microbiome, and grey: missing genes/pathways). Similarly, metabolites from different sources are also shown in different colours (blue: from gill endosymbiont, red: from *A. marisindica* itself, and orange: from intestinal food or microbiome). The compensation mechanism is revealed by the host’s collaborating with its symbionts to synthesise nutrients or their mutually using important metabolic intermediates. The combination of endogenous and exogenous energy sources is shown here to explain the adaptive mechanism of the entire *A. marisindica* holobiont.

### Host-microbe syntrophic interactions

The tripartite *A. marisindica* holobiont is supported by their tight metabolic complementarity (Figure 5). Both *Sulfurovum alviniconcha* CR and the gut microbiome of the Wocan *A. marisindica* possess typical metabolic pathways for synthesising carbohydrates, amino acids, and vitamins/cofactors and transporters for supplying these to the host. *Sulfurovum alviniconcha* CR uses the rTCA cycle to fix carbon and synthesises 20 amino acids and 4 vitamins/cofactors (Supplementary Figure S10A). The gut microbiome as a whole possess biosynthetic pathways for 10 amino acids and 4 vitamins (Supplementary Figure S10A), among them all of the 10 amino acids and 2 of the vitamins are shared with *Sulfurovum alviniconcha* CR, but the vitamins thiamine and nicotinate (and its derivative nicotinamide) are unique to the gut microbiome. Four amino acids and eight vitamins/cofactors cannot be synthesised *de novo* by either the symbiont system alone (Supplementary Figure S10B) and their production requires the complementary metabolic pathways of the host and symbionts to collaborate. For example, only *Sulfurovum alviniconcha* CR is capable of synthesising tryptophan yet it lacks genes for tryptophan metabolism, whereas the host genome contains the full tryptophan metabolic pathway from tryptophan to quinolinate. The host is further able to use quinolinate as a principal precursor to synthesise nicotinate and nicotinamide (vitamin B3) (Figure 5). Neither the host nor *Sulfurovum alviniconcha* CR alone can synthesise thiamine (vitamin B1), and the host lacks the thiamine transporter (*THTR*) for absorbing thiamine extracellularly. However, *Sulfurovum alviniconcha* CR can produce the thiamine phosphate precursor and pass it to the host. Thiamine is then synthesised as indicated by the highly expressed *PHO* that catalysing the conversion of thiamine phosphate to thiamine in the gill (Figure 5 and Supplementary Figure S9B). Similarly, the host cannot synthesise pantothenate but can obtain it from *Sulfurovum alviniconcha* CR in order to synthesise coenzyme A (Figure 5). Fatty acids (FAs) are essential nutrients required by most animals (Pranal et al., 1996). Holo-[carboxylase] serves as a biotin carrier protein and is essential in the biosynthesis of fatty acids in *A. marisindica*. Since only *Sulfurovum alviniconcha* CR can synthesise biotin (Supplementary Figure S10A), *A. marisindica* likely uses biotin derived from its endosymbionts *Sulfurovum alviniconcha* CR. Although *Sulfurovum alviniconcha* CR only possess biosynthetic pathways for saturated FA precursors (Figure 5), they may provide these precursors to the host, which can continue the FA biosynthesis by using the genes *MCH* and *fasN*, both of which are highly expressed in the gills (Figure 5 and Supplementary Figure S9B).

Neither *Sulfurovum alviniconcha* CR nor gut microbiome alone are able to generate all nutrients needed by the host (Supplementary Figure S10B). For example, the host expresses highly active pathways of pancreatic secretion and bile secretion in addition to metabolic pathways of folate and octadecanoic acid (Figure 5 and Supplementary Figure S9B), among other nutrients that cannot be synthesised by the host or endosymbiont. Numerous genes responsible for key hydrolases that are responsible for breaking down macromolecules, and specialised transport proteins are highly expressed and enriched in the intestine (Supplementary Figure S2C, S11A and S12A). In addition, the gut microbial enzymes include hydrolases (30.6–34.7%) (Dataset S1), transferases (26.7–28.2%), and oxidoreductases (9.6–17.4%). Large amounts of multi-exohydrolase complexes in gut microbiome may promote the host’s intestinal nutrient digestion (Supplementary Figure S9C, Table S3). For example, lactic acid bacteria (LAB) in the gut are found to possess a major facilitator, sugar transporters and enzymes for utilising large carbohydrate molecules. The gut microbiome even encodes additional enzymes such as oligoendopeptidase F (*pepF1*) and alginate lyase (*algL*) that can enhance digestion. Importantly, Campylobacterota in the gut are chemoautotrophic and found to encode the Sox system and [NiFe]-hydrogenases, and fix carbon with a complete rTCA cycle (Figure 5).

### Strategies of symbiosis maintenance

*Sulfurovum alviniconcha* CR lacks genes to assemble surface layer proteins (SLPs) or capsular polysaccharides (CPs). Nevertheless, *Sulfurovum alviniconcha* CR encodes and actively expresses transmembrane signalling receptors, lipid A and its modification (Supplementary Figure S11B). *Sulfurovum alviniconcha* CR does not encode putative virulence-related proteins (*pag*) for Cationic antimicrobial peptides (CAMPs) resistance, but its genome harbours the *eptA* gene (Supplementary Figure S11B) involving in bacterial surface charge modification. In addition, genes encoding various proteases (e.g. subtilisin-like serine proteases) and the type II secretion system (T2SS) are highly expressed in *Sulfurovum alviniconcha* CR (Supplementary Figure S11B), along with Sec and Tat secretory pathways. On the other hand, genes involved in the assembly of bacterial cloaks (CP, SLP), lipopolysaccharide (LPS), and other surface-associated antigens responsible for bacterial adhesion to the intestinal epithelium and activating the complement system (Sára and Sleytr, 2000; Futoma-Koloch, 2016) are found in the gut microbiome. Surface-layer glycoprotein variation in the gut microbiome is evident from the differential expression of S-layer genes, a type of antigenic variation responding to the lytic activity of the host immune system (Supplementary Figure S11B). In the gut microbiome of *A. marisindica*, genes encoding for sialate O-acetylesterase (SIAE) are highly expressed (Supplementary Figure S9C), which indicate their active sialic acid degradation in the gut.

The host immune system responds differently to *Sulfurovum alviniconcha* CR and the gut microbiome. The gills harbour a much higher abundance of bacteria than the gut (Supplementary Figure S6), but with weaker host immune responses (Figure 6A). Pattern recognition receptors (PRRs) are essential in the host’s innate immune system. They can be divided into membrane-bound PRRs and cytoplasmic PRRs. Genes encoding membrane-bound C-type lectin receptors (CLRs) and cytoplasmic RIG-I-like receptors (RLRs) are more active in the intestine than in the gills of host invertebrates (Figure 6A). Toll-like receptors (TLRs) recognise structurally conserved molecules derived from microbes and activate immune responses. Genes encoding an endosomal TLR13 are highly expressed in the gill tissue, similar to the finding in the symbiont-hosting gills of the vent mussel *Bathymodiolus platifrons* (Sun et al., 2017). In the gut, however, membrane-bound TLR2 and TLR6 are more active (Figure 6A). Once the host recognises the symbionts, the gut and gills take different approaches to deal with the invading symbionts. In the gut tissue of *A. marisindica*, the component cascade is activated as indicated by the highly expressed complement component 1 complex (*C1*) and complement C3 (*C3*) (Figure 6A and 6B). In the gill tissue, genes encoding the signal repressor NF-κB1 (*NFKB1*), negative regulators (NF-κB inhibitor zeta (*NFKBIZ*)), and TNFAIP3-interacting protein 1 (*TNIP1*) of NF-κB response, all of which involved in attenuation of NF-κB, are highly expressed (Figure 6A). Genes encoding four members of the GiMAP gene family are also highly expressed in the gill tissue (Figure 6A). In addition, genes involved in TNF signalling, the MyD88-independent TLR signalling pathway, and leukocyte differentiation, which related to antimicrobial activity, are enriched in the gills (Supplementary Figure S12B).

**Figure 6.**
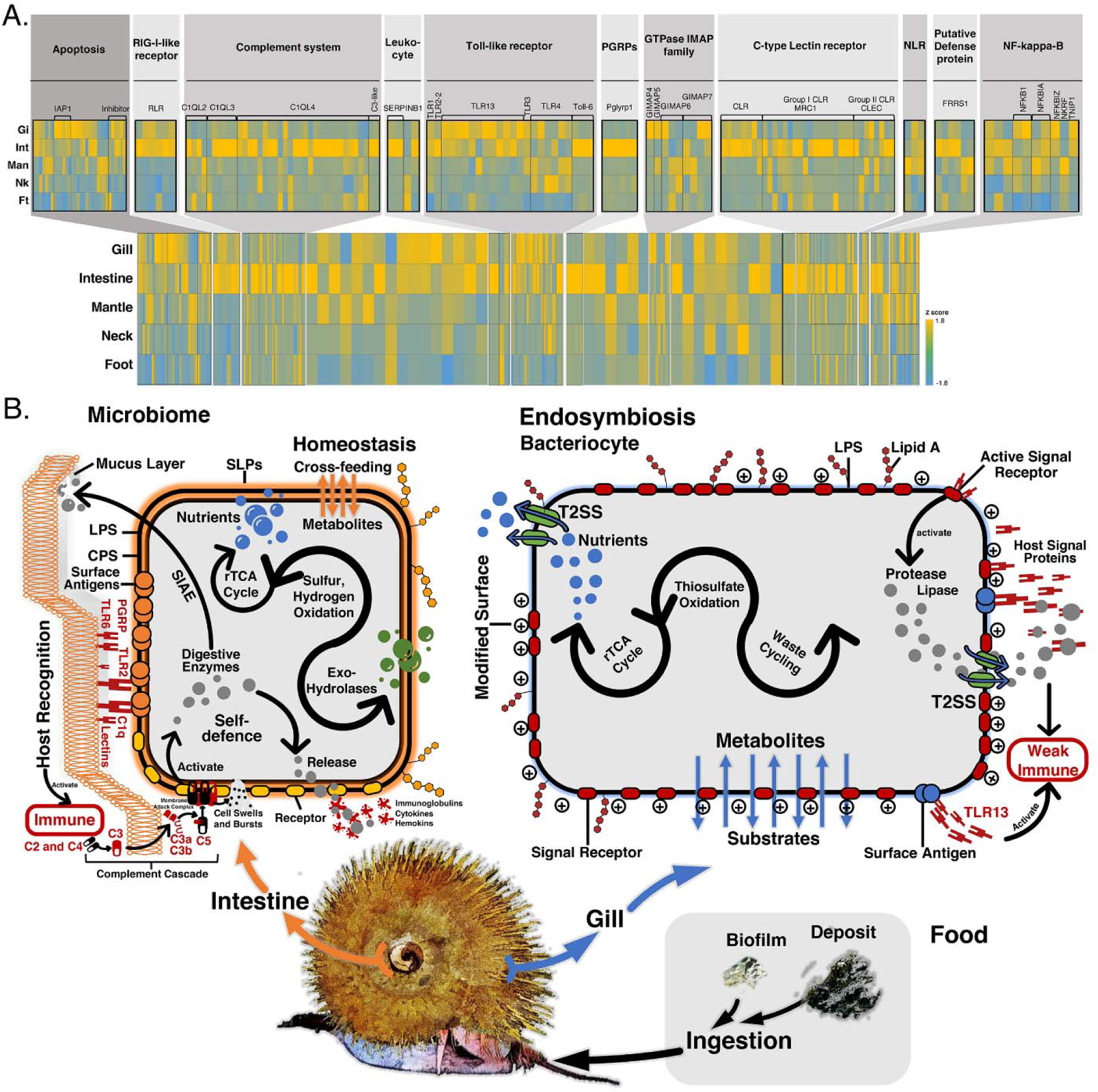
Symbiosis constraints of the *Alviniconcha marisindica* holobiont. (**A**) Heat map of the transcriptional activity of genes involved in host innate immunity in the foot, neck, mantle, intestine, and gill tissues showing distinct immune-expression profiles regulated by the two types of symbionts in the *A. marisindica* snail. Each grid in the heat map represents an identified gene. The colour represents the gene expression level (based on normalised TPM values of the selected tissues). The annotated gene names and their functional classifications are listed on the top side. (**B**) Symbiosis model of the *A. marisindica* holobiont with two different symbiotic constraints and interactions with the external environment. All pattern recognition receptors (PRRs) and pathogen-associated molecular patterns (PAMPs) shown here are identified from the genome and transcriptome data. SLPs, surface layer proteins; LPS, lipopolysaccharide; CPS, capsular polysaccharides; SIAE, sialate O-acetylesterase; PGRPs, peptidoglycan recognition proteins; TLRs, toll-like receptors; C1q, complement component 1q; T2SS, type II secretion system.

## Discussion

*Sulfurovum alviniconcha* CR has a relatively compact (1.47-Mbp) and streamlined genome. As maintaining the symbionts involves costs (Douglas and Smith, 1983; Meyer and Weis, 2012), the host may prefer a cellularly economised symbiont genome for energetic efficiency (Nicks and Rahn-Lee, 2017). A small endosymbiont genome may also enhance growth efficiency and intracellular competitiveness (Moran, 2002). *Sulfurovum alviniconcha* CR lacks most flagellar or chemotaxis genes like its Campylobacterota close relatives from deep-sea sediments (Inagaki et al., 2004) and mounds (Nakagawa et al., 2007), whereas the Campylobacterota endosymbionts of *Alviniconcha boucheti* from Kilo Moana vent field at the Eastern Lau Spreading Centre has complete flagellar genes but no genes for the chemotactic signaling system (Beinart et al., 2019). The Campylobacterota endosymbionts of *Alviniconcha boucheti* are thought to be motile at free-living stage and such motility machinery could be used for finding a host. In this case, the non-motile *Sulfurovum alviniconcha* CR has a different machinery for adhesion to or interaction with *Alviniconcha marisindica*. Several symbioses have shown that motility and chemotaxis are not indispensable for recruiting symbionts from the environment, for example, some non-motile sulfate-reducing bacteria and methane-producing archaea in marine sediments use adhesins to colonise their host (Orphan et al., 2001; Raina et al., 2019). In addition, low-fidelity repair in the *Sulfurovum alviniconcha* CR genome increase its mutagenic potential, and such genomic plasticity has been found in human/animal pathogenic Campylobacterota (Kang and Blaser, 2006; Monack et al., 2004) and deep-sea vent Campylobacterota (Nakagawa et al., 2007), leading to micro-diversity increasement that confers a competitive advantages enabling bacteria persist in infections (Kang and Blaser, 2006; Monack et al., 2004) or thriving in ever-changing environments such as deep-sea vents (Nakagawa et al., 2005; Nakagawa et al., 2007). The *Sulfurovum alviniconcha* CR genome has the core of virulence for important animal pathogens, indicating its infectivity. Even if the *Sulfurovum alviniconcha* CR genome lacks many genes, it shows the ability to face with a changing environment, infect the animal host and survive intracellularly.

The host selectivity of endosymbionts in *Alviniconcha* snails is low when compared to other chemosymbiotic animals such as tubeworms (Beinart et al., 2012; Beinart et al., 2019; Yang et al., 2020) which may harbour a high diversity (even multiple classes) of endosymbionts with different types of metabolism within a single host (Beinart et al., 2012). This probably reflects the combined effect of environment selectivity on the available phylotypes and differences in vent fluid chemistry (Wang et al., 2017). A rarely discussed anatomical characteristic of the gill endosymbionts in the *Alviniconcha* species is that these endosymbionts residing inside bacteriocytes are present in a state between truly intracellular and extracellular (Endow and Ohta, 1989). Electron microscopy revealed that the vacuoles in bacteriocytes housing the endosymbionts are exposed to the ambient seawater through duct-like openings (Endow and Ohta, 1989). Considering the aggregated distribution of symbionts near the more exposed, outer surface of the bacteriocytes, the *Alviniconcha* gill symbionts are in a ‘semi-endosymbiotic’ condition (Windoffer and Giere, 1997), which likely provides *Alviniconcha* snails with the ability to exchange or reacquire gill symbionts according to the local habitat and environment selectivity through the endocytosis of free-living bacteria. Such ‘semi-endosymbiotic’ condition provides gill endosymbiont populations with heterogeneous genomes regarding metabolic genes along the chemoautotrophic pathways that may enable the utilisation of diverse substrates. *Sulfurovum alviniconcha* CR is capable of anaerobically oxidising thiosulfate and hydrogen. The environmental sulfide is conversed to polysulfides in *Sulfurovum alviniconcha* CR and then bacterial organic polysulfides such as sulfur-containing amino acids are degraded to produce intracellular thiosulfate for oxidation to produce cellular energy (Figure 5). This method of sulfide utilisation and storage is different from those seen in many deep-sea holobionts such as in siboglinid tubeworms, where the host haemoglobin binds to and transports the sulfides to endosymbionts for direct oxidation or storage in bacterial sulfur globule proteins (Yang et al., 2020), and in *Bathymodiolus* mussels, where the host oxidise sulfides and provide a reservoir of thiosulfate for the endosymbionts’ oxidation (Ponnudurai et al., 2020). We supposed that the storage of environmental sulfides in *Sulfurovum alviniconcha* CR’s polysulfides and the utilisation of thiosulfate degraded from these intracellular sulfur compounds, is more efficient than those symbionts which use thiosulfate provided by extracellular host tissues.

The synergistic biosynthesis of nutrients in *A. marisindica* gives the holobiont a capability of nutrient production that is controlled by mutual supply of intermediates between the host and the endosymbionts. Although the semi-endosymbiotic mode of housing the gill endosymbiont provides *Alviniconcha* with the ability to utilise a rather wide array of bacteria as gill endosymbiont (Beinart et al., 2019), it comes at a cost in that some symbiont phylotypes may lack genes for certain syntrophic interactions. Although the digestive tract is substantially reduced in the adult snail (Warèn and Bouchet, 1993), *A. marisindica* has a functioning gut which contains faecal-like black substances suggesting that this snail ingests food by either grazing or filter-feeding like *A. hessleri* from the Mariana Back Arc Basin (Warèn and Bouchet, 1993) and *A. marisindica* from the Central Indian Ridge (Suzuki et al., 2005). By supplying genes and functions that are missing in *Sulfurovum alviniconcha* CR and the host, the gut microbiome help ensures the nutritional viability of the holobiont as a whole. For example, *pepF1* and *algL* genes in the gut microbiome that enhance digestion are lacking in the host snail and thus provide direct evidence that the gut microbiota has the potential to fulfil the nutritional demands of the holobiont. Overall, the results show that the gut microbiome has the potential to provide nutrition benefits to the *Alviniconcha* snail, an aspect of symbiosis that has been neglected in previous studies of many deep-sea endosymbiotic holobionts. As the *Alviniconcha* species has a semi-endosymbiotic model of gill symbiosis, the endosymbionts have flexible symbiotic associations with the host, which may at times impose nutritional limits on the holobiont system. In such situations, the gut likely contributes to keep the holobiont functionally versatile, ensuring its thriving in vent fields featuring different geochemical environments and available energy sources.

*Sulfurovum alviniconcha* CR lacks two common bacterial physical “cloaks” – SLPs and CPs that protect intracellular bacteria from host defences but also being recognised by the host as immunodominant antigens (Zamze et al., 2002; Sára and Sleytr, 2000). The absence of CPs likely helps *Sulfurovum alviniconcha* CR enter host cells (Deghmane et al., 2002) and reduces the risk of polysaccharide recognition by the host immune system (Zamze et al., 2002). In addition, lipid A modification enzymes and surface signal receptors help bacterial pathogens to avoid detection by TLRs (Thakur et al., 2019), this may also apply to *Sulfurovum alviniconcha* CR. CAMPs are key components of the host’s innate immune response (Le et al., 2017; Noore et al., 2013). The presence of the *eptA* gene (Supplementary Figure S11B) involved in surface charge modification implies that *Sulfurovum alviniconcha* CR increases its surface positive charge to repel CAMPs. T2SS enables the transport of various cytoplasmic proteins into extracellular milieu, including bacterial toxins and degradative enzymes such as proteases and lipases. Previous study of tubeworm endosymbionts shows that the endosymbiont may use serine proteases to modulate the host’s immune response by diminishing the function of host signal proteins (Yang et al., 2020). *Sulfurovum alviniconcha* CR may use a similar strategy. Surface antigenic molecules of *Sulfurovum alviniconcha* CR are distinct from the gut microbiome, indicating its host-specific immune-evasion mechanisms. In gut tissues, the mucus layer is the interface between the gut flora and the host, and sialic acids are prominent carbohydrates of the intestinal mucus layer (Schroeder, 2019). Thus sialic acid breakdown of the gut microbiome indicates the way of intestinal bacterial encroachment and survival. Such differences in interaction with the host can lead to the establishment of different animal-microbe associations (Koropatnick et al., 2004).

Accordingly, the *A. marisindica* host has distinct recognition profiles for *Sulfurovum alviniconcha* CR and the gut microbiome (Figure 6A). After being recognised by the host, the invading symbionts will be controlled by the host’s different corresponding strategies. The component cascade is activated in the gut which attack the microbe’s cell membrane and eliminate microbes to control bacterial infections (Janeway et al., 2001) (Figure 6B). In the gills, potential attenuation of NF-κB are important as they grant the invaded cells additional protection (Burns et al., 2017; Best et al., 2019), and four members of the GiMAP gene family play critical roles in constraining and compartmentalising pathogens within cells (Weiss et al., 2013; Hunn et al., 2011). In addition, genes responsible for the majority of antimicrobial activity are enriched in the gills (Supplementary Figure S12B). In this case, the gill tissue shows a strong potential to constrain the intracellular symbionts and resist environmental invasion, which also indicate the ability of *Sulfurovum alviniconcha* CR to evade these host antimicrobial activities at the free-living stage. The weak host immune responses in the gills (Figure 6A) indicate *Sulfurovum alviniconcha* CR are more adept at evading recognition by the host immune system or inhibiting activation of the host immune system. Overall, the results show that *Sulfurovum alviniconcha* CR may have evolved an immunomodulation mechanism that they modulate the cell’s outermost layer and release proteins enabling them to effectively evade recognition by the host immune system. In addition, the semi-intracellular position of *Sulfurovum alviniconcha* CR may allow it to avoid areas of high lysosomal activity in host cells that are part of the host self-defence mechanism (Endow and Ohta, 1989). The balance between the host’s immune activity and bacterial counter-defence contributes to the complexity of the persistent symbioses.

We show, through hologenomic and holotranscriptomic analyses, that the *Alviniconcha marisindica* holobiont is more complex than previously recognised, being a tripartite system with the host snail and gill endosymbiont additionally supported by functional gut microbiome. The relative importance of each partner in *A. marisindica* may fluctuate depending on the immediate availability of resources impacting the interplay downstream, as has been shown for other invertebrate symbioses (Morris et al., 2019; Belda et al., 1993). We unravel complex interactions among symbiotic parties in the *A. marisindica* holobiont, which deepen our understanding of the adaptation of many dominant chemosymbiotic holobionts that rely on gill endosymbionts for nutrition and also retain a functional gut, such as *Alviniconcha*’s sister genus *Ifremeria* (Windoffer and Giere, 1997), peltospirid snails (Chen et al., 2018), and *Bathymodiolus* mussels (Page et al., 1991).

## Materials and Methods

### Sample collection and nucleic acid preparation

*Alviniconcha marisindica* individuals were collected from a water depth of 2,919 m at the Wocan vent site on the Carlsberg Ridge (CR) of the northwestern Indian Ocean (60.53°E, 6.36°N) (Supplementary Figure S13). Sampling was conducted using the human occupied vehicle (HOV) *Jiaolong* onboard the research vehicle *Xiangyanghong 9* on March 19, 2017. Snails were placed into an insulated bio-box with a closed lid using a manipulator to minimise changes in water temperature. *Jiaolong* took approximately 2.5 hours to return to deck. Once the snails were onboard the research vessel, all specimens were immediately flash-frozen in liquid nitrogen and then transferred to a -80°C freezer for storage. The morphological observation and molecular taxonomy of snail samples were shown in Supplementary Note 1.

The frozen snails were thawed in RNAlater® (Invitrogen, USA) on ice, dissected with different tissues fixed separately in RNAlater®, and then prepared for nucleic acid extraction. A single specimen of *Alviniconcha marisindica* from Wocan was used for the holobiont genome assembly. The foot and neck muscles were used for host genomic DNA extraction, and the endosymbiont-harbouring gills were used for endosymbiont DNA extraction. A total of three snail individuals, including the one used for identifying the host genome, were dissected into 7–10 tissue types each with RNA extraction performed on the different tissues. The gills of 10 other individuals were divided into anterior and posterior parts, and the DNA of these 20 parts were extracted separately for metagenome sequencing. The intestines of the three individuals were dissected for total DNA and RNA extraction for the gut microbiome, and the gills of these individuals were also dissected for total RNA extraction (Supplementary Note 1). Genomic DNA (gDNA) was extracted using the E.Z.N.A.® Mollusc DNA Kit (Omega Bio-tek, Georgia, USA) and then purified using Genomic DNA Clean & Concentrator^TM^-10 Kit (Zymo Research, CA, USA) according to the manufacturer’s protocol. Total DNA of the gills and that of the intestines were extracted using the same protocol. Total RNA was extracted using Trizol (Invitrogen, USA) from different tissues following the manufacturer’s protocol and prepared for RNA-Seq. Nucleic acid quality was evaluated using agarose gel electrophoresis and a BioDrop µLITE (BioDrop, Holliston, MA, US), and nucleic acid concentrations were quantified using a Qubit fluorometer v3.0 (Thermo Fisher Scientific, Singapore).

### Library construction and sequencing

Genomic DNA was aliquoted and submitted to three sequencing platforms: Illumina, PacBio Sequel, and Oxford Nanopore Technologies (ONT). A library with a 350 bp insert size was constructed from gDNA following the standard protocol provided by Illumina (San Diego, CA, USA). After paired-end sequencing of the library at Novogene (Beijing, China), approximately 50 Gb of Illumina NovaSeq reads with a read length of 150 bp were generated. Illumina sequencing of total DNA from the gills and that of total DNA from the intestines were conducted similarly, with approximately 50 Gb of reads generated from each gill sample for endosymbiont genome assembly, approximately 6–8 Gb of reads generated from each of 20 gill filaments for symbiont genetic diversity analysis, and approximately 12 Gb of reads generated from each of three intestine specimens for metagenome analysis (see overview of sequencing data in Supplementary Note 2).

For preparation of the single-molecule real-time (SMRT) DNA template for PacBio sequencing, the gDNA was sheared into large fragments (10 kb on average) using a Covaris® g-TUBE® device and then concentrated using AMPure® PB beads. DNA repair and purification were carried out according to the manufacturer’s instructions (Pacific Biosciences). The blunt adapter ligation reaction was conducted on purified end-repair DNA, and after purification DNA sequencing polymerases became bound to SMRTbell templates. Finally, the library was quantified using a Qubit fluorometer v3.0. After sequencing with the PacBio Sequel System at the Hong Kong University of Science and Technology (HKUST) and Novogene, approximately 72LGb of long reads were generated, with reads less than 4 kb in length discarded.

For ONT sequencing, a total of 3–5 μg of gDNA were used for the construction of each library following the ‘1D gDNA selecting for long reads (SQK-LSK109)’ protocol from ONT. Briefly, gDNA was repaired and end-prepped as per standard protocol, before it was cleaned up with a 0.4× volume of AMPure XP beads. Adapter ligation and clean-up of the cleaned-repaired DNA were performed as per the standard protocol and the purified-ligated DNA was eluted using elution buffer. The DNA library was mixed with sequencing buffer and loading beads before it was loaded onto the SpotON sample port. Finally, sequencing was performed following the manufacturer’s guidelines using the FLO-MIN106 R9.4 flow cell coupled to the MinION^TM^ platform (ONT, Oxford, UK). Raw reads were base-called according to the protocol in MinKNOW and written into fastq files, and 9.8 Gb of long reads were generated with reads less than 4Kb discarded. MinION sequencing of total DNA from one gill specimen was conducted using the same procedures, generating 3.5 Gb of reads for endosymbiont genome scaffolding (see details of PacBio and ONT library construction in Supplementary Note 2). Illumina reads from gDNA were used for the genome survey of the Wocan *Alviniconcha marisindica*, and PacBio and ONT reads were used for the genome assembly (Supplementary Note 3).

For eukaryotic transcriptome sequencing of different tissues, a 250–300 bp insert cDNA library of each tissue was constructed after removing the prokaryotic RNA and sequenced on the Illumina NovaSeq platform at Novogene to produce 150 bp paired-end reads. Since the RNA of gills includes the sequences from both the host and the symbiont, another 250–300 bp insert strand-specific library of each gill specimen was constructed using Ribo-Zero™ Magnetic Kit to sequence both eukaryotic and microbial RNA. Therefore, two sets of transcript sequencing data were produced for the gills, one for both the host and the symbiont, and the other for only the host. The meta-transcriptome sequencing of the intestine was conducted using the same methods. Approximately 5–10 Gb of reads were generated from each tissue.

### de novo *hybrid assembly of the host genome*

Trimmomatic v0.39 (Bolger et al., 2014) was used to trim the Illumina adapters and low-quality bases (base quality ≤ 20). The genome size of *A. marisindica* was estimated to be 809.1LMb using the 17-mer histogram generated (Supplementary Note 3) and the genome heterozygosity was evaluated as 0.88% using GenomeScope (Vurture et al., 2017). Several genome assembly pipelines were applied to assemble the genome with PacBio and ONT reads, including PacBio-only approaches (e.g. minimp2+miniasm (Li, 2016) and wtdbg2 (Ruan and Li, 2019)) and PacBio-ONT hybrid approaches (e.g. MaSuRCA version 3.2.8 (Zimin et al., 2013), FMLRC (Wang et al., 2018) + smartdenovo (Ruan, 2018) and FMLRC (Wang et al., 2018) +wtdbg2 (Ruan and Li, 2019)). The detailed settings of each assembly pipeline are shown in Supplementary Note 3.

A comparison of the assembly statistics of different pipelines (Supplementary Note 3) showed that the FMLRC+wtdbg2 assembly was the best and therefore this assembly was used in the downstream analyses. The assembly was carried out as follows: the ONT reads were concatenated with PacBio reads and error corrected with Illumina reads using FMLRC (Wang et al., 2018). This hybrid error correction method was selected based on previous benchmarking analysis on the available tools using Illumina reads for correction of PacBio/ONT long reads (Fu et al., 2019). The corrected long reads were then assembled using wtdbg2 using the setting “-x preset2” (Ruan and Li, 2019). Bacterial contamination was removed from the assembly using a genome binning method in MetaBAT 2 (Kang et al., 2015) and MaxBin 2.0 (Wu et al., 2016) (Supplementary Note 3). The Illumina reads were mapped to the clean assembly with Bowtie2 (Langmead and Salzberg, 2012), and only uniquely mapped reads were retained. The resultant sorted .bam file was used to correct errors in the assembly using Pilon v1.13 (Walker et al., 2014). Two rounds of error correction were performed. Redundant genomic assembled contigs from highly heterozygous regions were then removed using Redundans (Pryszcz and Gabaldón, 2016) with the settings of “--identity 0.8 --minLength 5000”.

### Quality check of the assembled host genome

We monitored the genome assembly completeness and redundancy using the metazoan Benchmarking Universal Single-Copy Orthologs (BUSCOs) v4.0.6 pipeline against the Metazoan dataset (Simão et al., 2015). A total of 921 out of the 954 searched BUSCO groups (96.5%) were complete in the assembled genome, and only 2.3% BUSCOs were missing, suggesting a high level of completeness of the *de novo* assembly (Supplementary Table S1). QUAST v5.0.2 (Gurevich et al., 2013) was used to check genome assembly quality with PacBio and ONT reads (Supplementary Table S2).

### Annotation of the host genome

The Wocan *Alviniconcha* host genome annotation pipeline generally followed a previously published procedure (Sun et al., 2017). Briefly, the repeat content and the transposable elements were predicted and classified using the RepeatMasker pipeline (Smit and Hubley, 2010) which searched against the known repeat library in Repbase and also the species-specific repeat library constructed by RepeatModeler (Supplementary Note 4).

Two versions of transcriptome assembly, i.e. the *de novo* assembled version and the genome-guided version, were independently assembled using Trinity v2.8.5 (Grabherr et al., 2011) and concatenated. Sequences with similarity over 0.97 were clustered with cd-hit-est (Li and Godzik, 2006). Maker v3.0 (Cantarel et al., 2008) was used to annotate the genome. In the first round of Maker annotation, only the transcriptomic evidence was used, and only a gene model with an annotation edit distance (AED) score less than 0.01 (Supplementary Figure S1) and predicted protein length over 200 amino acids was reported. The resultant genome annotation *.gff* file was used to train another *de novo* gene predictor, Augustus v3.3 (Stanke and Morgenstern, 2005). The gene model with only one exon with an incomplete open reading frame and inter-genic sequences less than 3 Kb was removed. The rest of the *bona-fide* gene models were used to train Augustus. In the second round of Maker, evidence from three different sources, i.e. the transcriptome, proteins from the Swiss-Prot database, and Augustus, were merged using EvidenceModer (Haas et al., 2008). The merged data was also integrated using Maker.

Gene functions were determined by using BLASTp to align the candidate sequences with NCBI non-redundant (NR) and Swiss-Prot protein databases with the settings of “-evalue 1e-5 - word_size 3 -num_alignments 20 -max_hsps 20”. Blast2GO^®^ (Götz et al., 2008) together with EggNOG mapper (Huerta-Cepas et al., 2017) was applied to assign Gene Ontology (GO) terms and clusters of orthologous groups (COGs) to the protein sequences via GO and EggNOG databases. The Kyoto Encyclopedia of Genes and Genomes (KEGG) Automatic Annotation Server (KAAS) (Kanehisa and Goto, 2000) was used to conduct the KEGG pathway annotation analysis via the bidirectional best hit method. The Pfam database was searched using profile hidden Markov models (profile HMMs) (with an e-value of 0.001) to classify the gene families (El-Gebali et al., 2019).

### Host gene family identification and phylogenomic analysis

A total of 26 lophotrochozoan genomes were analysed for clues to the gene family evolution (Supplementary Note 4). Orthologs among all species were deduced via the OrthoMCL pipeline (Li et al., 2003) with the BLASTp threshold set to 1e-5. Only single-copy genes in at least two-thirds of the taxon sampled (i.e. in at least 18 species) were used in the phylogenetic tree analysis, resulting in 492 orthologous groups. Protein sequences within each orthologue was aligned using MUSCLE with the default settings; spurious sequences and poorly aligned sequences were trimmed using TrimAL v1.4 (Capella-Gutiérrez et al., 2009). The final alignment of each orthologue was concatenated with partition information for the phylogenetic analysis using RaxML v8.2.11 (Stamatakis et al., 2005) with the GTR + Γ model. The MCMCTree v4.7 (Reis and Yang, 2011) was used for tree dating. The root calibration point was set to 590 Ma in MCMCTree, and the LG+Γ model of evolution was selected. Time frame constraints imposed to calibrate the topology tree generated from RAxML are shown in Supplementary Note 4. The MCMCTree was run for 1.0 × 10^7^ generations, sampling every 1.0 × 10^3^ and discarding 20% of the samples as burn-in. Gene family expansion/contraction analysis was performed using CAFÉ v3.1 (Han et al., 2013). Only a family level with *P*<0.01 and *P<*0.01 deduced by the Viterbi method was considered to be expanded or contracted.

### Microbial metagenome assembly, annotation, and functional analysis

For microbial metagenome assembly of the gill, Trimmomatic v0.39 (Bolger et al., 2014) and FastUniq (Xu et al., 2012) were used to trim the Illumina reads and remove duplicates. The bacterial abundance of gill metagenomic sequences was deduced using Kaiju (Menzel et al., 2016) based on the subset of the NCBI BLAST *nr* database containing all proteins belonging to Archaea, Bacteria, and Viruses. The clean reads were assembled using metaSPAdes v3.13.1 (Bankevich et al., 2012) with k-mer sizes of 21, 33, 55, 77, 99, and 127 bp, and the products were pooled. Contigs potentially belonging to the campylobacterotal endosymbiont genome were separated from its host genome using a genome binning method as described in previous studies (Albertsen et al., 2013; Yang et al., 2020) (Supplementary Note 5). A genome of presumably parasitic Mollicutes was removed (Supplementary Figure S3A). Contigs of the endosymbiont genome were further determined using MetaBAT 2 (Kang et al., 2015) and MaxBin 2.0 (Wu et al., 2016), assessed using CheckM v1.1.2 (Parks et al., 2015), and further scaffolded using SSPACE-LongRead v1.1 (Boetzer and Pirovano, 2014) and *npScarf* (Cao et al., 2017) by adding ONT long reads. The newly assembled scaffolds were binned again using the above pipeline. GapFiller v1.10 (Boetzer and Pirovano, 2012) and Gap2Seq v3.1 (Salmela et al., 2016) were used to fill the gaps in the binned endosymbiont genome. CheckM v1.1.2 (Parks et al., 2015) was used to estimate the completeness and potential contamination of the binned genome. Coding sequences (CDS) in the genome of the *Alviniconcha* endosymbiont were predicted and translated using Prodigal v2.6.3 (Hyatt et al., 2010). Gene function annotation of the predicted protein sequences followed the same pipeline as that described above for the host snail (Supplementary Note 5). The protein sequences were annotated based on GO, EggNOG, KEGG and Pfam databases.

For the gut metagenome assembly, reads were trimmed and duplicates removed as described above. The host’s interference in the analysis of intestinal content was minimised by removing reads that were aligned with the host genome using Bowtie2 (Langmead and Salzberg, 2012) before the assembly. The remaining reads were assembled using metaSPAdes v3.13.1 (Bankevich et al., 2012) with the same settings as above. The abundance and systematic classification of intestinal metagenomic microbial sequences were carried out using Kaiju (Menzel et al., 2016) (Supplementary Figure 3B and 6B). Network analysis of intestinal microbes was conducted based on their relative abundance. To reduce the complexity of the datasets, relative abundances higher than 0.01% were retained for the construction of the network. All pairwise Spearman’s rank correlations were calculated in the R package “picante”. Only robust (*r*>0.8 or *r*<-0.8) and statistically significant correlations (*P*<0.01) are shown in the network. Network visualisation and modular analysis were conducted in Gephi v0.9.2. Prodigal v2.6.3 (Hyatt et al., 2010) was used to predict and translate the coding sequences in the intestinal metagenome, and BLASTp was then used to align the candidate sequences with the NCBI NR protein database. The systematic assignment of each protein was imported to MEGAN v5.7.0 (Huson et al., 2011) using the lowest common ancestor (LCA) method with the parameters of Min Score 50, Max Expected 0.01, Top Percent 5, and LCA Percent 100. Based on the systematic results, the microbial protein sequences were selected for further gene functional analysis, following the gene annotation pipeline described above. Blast2GO® (Götz et al., 2008) and EggNOG mapper (Huerta-Cepas et al., 2017) were applied to assign GO and COG terms to the intestinal prokaryotic protein sequences. KAAS (Kanehisa and Goto, 2000) was used to annotate the KEGG meta-pathway of intestinal flora using the single-directional best hit (SBH) method. All the annotated information of intestinal flora was in Dataset S2; the potential function and interaction of gut microbiome were shown in Supplementary Note 6.

### Phylogenomic analysis and genomic comparison of the endosymbiont

A total of 120 single-copy orthologous genes found in all genomes of nine Deltaproteobacteria (outgroup) and 111 campylobacterotal representatives by Proteinortho6 (Lechner et al., 2011) (BLAST threshold E = 1 × 10^-10^) were retained for phylogenomic analysis. Sequences of each orthologue were aligned using MUSCLE and trimmed using TrimAL (Capella-Gutiérrez et al., 2009). The final alignment of each orthologue was concatenated with partition information for the phylogenetic analysis using RaxML v8.2.11 (Stamatakis et al., 2005) with the GTR + Γ model. The gill endosymbiont of the Wocan *Alviniconcha marisindica* was compared with the endosymbiont of *Lamellibrachia* tubeworm (Patra et al., 2016), the epibiont of the giant tubeworm *Riftia pachyptila* (Giovannelli et al., 2016), and two free-living Campylobacterota (Giovannelli et al., 2016; Nakagawa et al., 2007) from deep-sea hot vents (Table 1), which were clustered within the same clade (Figure 3). Whole-genome ANI of orthologous gene pairs shared between two microbial genomes was calculated using fastANI (Jain et al., 2018) with the default settings. A Venn web tool (http://bioinformatics.psb.ugent.be/webtools/Venn/) was used to illustrate the shared and unique orthologous genes among the five Campylobacterota representatives (Figure 4b). Orthologous genes only present in the Wocan *A. marisindica* endosymbiont were classified as its unique genes. Orthologous genes that were present in all the other four reference genomes but not in the endosymbiont of *Alviniconcha* were classified as reduced genes (Supplementary Note 7). In addition, an HMM-based approach delta-bitscore (Wheeler et al., 2016) was used to identify loss-of-function mutations in shared orthologous genes of the five Campylobacterota (Dataset S3, Supplementary Note 7).

In addition to the above genomic comparisons, a total of 23 metagenome sequences were obtained from 13 Wocan *A. marisindica* snails. A genome binning method was used following the pipeline described in previous sections to assemble and extract another 22 endosymbiont genomes and the ANI among these 23 endosymbiont genomes was calculated (Jain et al., 2018) (Table 2 and Supplementary Note 8). The core genome shared across all 23 endosymbiont genomes was obtained using Proteinortho6 (Lechner et al., 2011) (BLAST threshold E = 1 × 10^-10^). The pan genome including isolate-specific genes were also detected. In addition, we also captured genome-wide variation of endosymbionts by comparing variations present in two parts of the gills (anterior and posterior) in each snail individual, across multiple snails (Supplementary Note 8). Selecting from the above 23 genomes, SNPs among 20 endosymbiotic isolates of the anterior and posterior gills from 10 snail individuals were called by aligning clean high-quality Illumina reads from each gill sample with a complete reference genome using the novel high-accuracy pipeline BactSNP (Yoshimura et al., 2019), in a single step. Pseudo genomes of input isolates were obtained. For each isolate, all contigs in the pseudo genome were concatenated into one sequence and submitted to phylogeny analysis using RaxML v8.2.11 (Stamatakis et al., 2005) under the GTR + CAT model.

### Quantification of gene expression level

For host transcriptome sequencing data, the raw reads of each tissue were trimmed with Trimmomatic v0.39 (Bolger et al., 2014), the gene expression level in each tissue was expressed in transcripts per million (TPM) with Salmon (Patro et al., 2017), and the number of read counts for genes was also included in the quantification results. For meta-transcriptome sequencing data of the symbionts, the same pipeline was followed, with a Salmon index built for the transcripts of symbionts obtained and translated from their genome data. The trimmed reads were then quantified directly against this index and expressed in TPM using Salmon (Patro et al., 2017). In addition, using this quantification method, the gene expression levels of the gills were produced from two sets of RNA sequencing data of the gills (one is a meta-transcriptome dataset including both the host and symbionts, and the other is a eukaryotic transcriptome including only host transcripts). The consistency of gene expression levels for the gills from these two sets of sequencing data also confirmed the accuracy of our transcript-level quantification in the host and its symbionts.

Differentially expressed genes were determined by DESeq2 using the normalisation method of Loess, a minimum read count of 10, and a paired test (*n* = 5). A gene was considered specifically expressed in a particular tissue based on its expression levels compared across all other tissue types (paired-test method). Genes overexpressed with over twofold changes and false discovery rate (FDR) < 0.05 when compared with other tissue types were considered to be highly expressed (Dataset S4). WEGO (http://wego.genomics.org.cn/cgi-bin/wego/index.pl) was used to plot GO annotations of highly expressed genes in the different selected tissues. Statistically overrepresented GO terms in the different tissues were identified through topGO package in R session (Alexa and Rahnenführer, 2009). The GO enrichment network is visualised using the Cytoscape application (Shannon et al., 2003). Differentially expressed genes of different tissues were shown in Supplementary Note 9.

### Data availability

All raw sequencing data generated in the present study are available from NCBI via the accession numbers SRR11781614–SRR11781681, and BioSample accessions SAMN14907812– SAMN14907827. The data generated in the present study have been deposited in the NCBI database as BioProject PRJNA632343. All software commands used in the host genome assembly are given in the Supplementary Information. The assembled transcriptome, predicted transcripts, and proteins are openly available from Dryad (DOI: XXX).

## Acknowledgements

This work was supported by grants from the China Ocean Mineral Resources Research and Development Association (DY135-E2-1-03), the Hong Kong Branch of Southern Marine Science and Engineering Guangdong Laboratory (Guangzhou) (SMSEGL20SC01), the Southern Marine Science and Engineering Guangdong Laboratory (Guangzhou) (GML2019ZD0409), and the Major Basic and Applied Basic Research Projects of Guangdong Province (2019B030302004-04) awarded to P.Y.Q., and the National Natural Science Foundation of China (NSFC) (grant no. 91951201). Thank Ms Pui Shuen (Joyce) Wong at BioCRF, HKUST for her assistance in PacBio sequencing. Alice Cheung edited the final version of the paper.

## Author Contributions

PYQ conceived the project. YY and JS designed the experiments. YZ and CW collected the *Alviniconcha* snails. CC dissected specimens. YY performed DNA extraction, RNA extraction, Nanopore and PacBio sequencing, gene expression and metabolic pathway analyses of the symbionts and the host *Alviniconcha* snail. YY performed genome assemblies, phylogenetic and genomic comparative analyses of the symbionts. JS performed genome assembly, phylogenetic and gene family analyses of the host *Alviniconcha* snail. JS and YY performed genome annotation of the host *Alviniconcha* snail. YY checked bacteria contamination, performed genomic comparative and the remaining analyses of the host *Alviniconcha* snail. YY prepared the figures and tables and drafted the manuscript. CC, JS, LY, CVD, JWQ and PYQ contributed to manuscript editing.

