## Supplementary Information for "Tripartite holobiont system in a vent snail broadens the concept of chemosymbiosis"

**This PDF file includes:**

- Figures S1 to S13
- Tables S1 to S3
- Supplementary Note 1 to Note 9
- Legends for Datasets S1 to S4
- SI References

**Other supplementary materials for this manuscript include the following:**

Datasets S1 to S4

Table S5. The nucleic acid preparation and Illumina paired-end sequencing information of different tissues/organs from 16 *Alviniconcha marisindica* individuals in this study. 28

**Supplementary Figures**

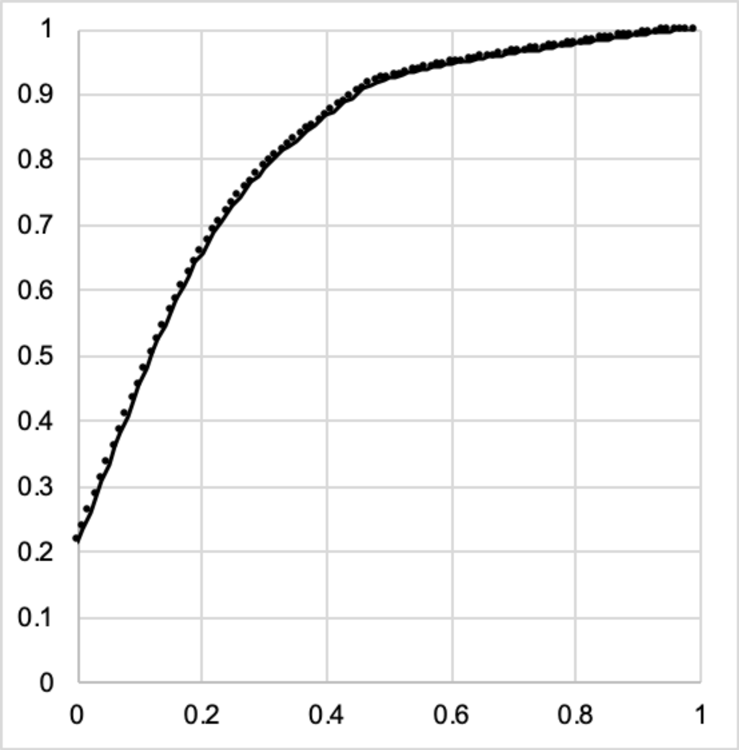

**Figure S1.** AED score distribution deduced from all of the gene models.

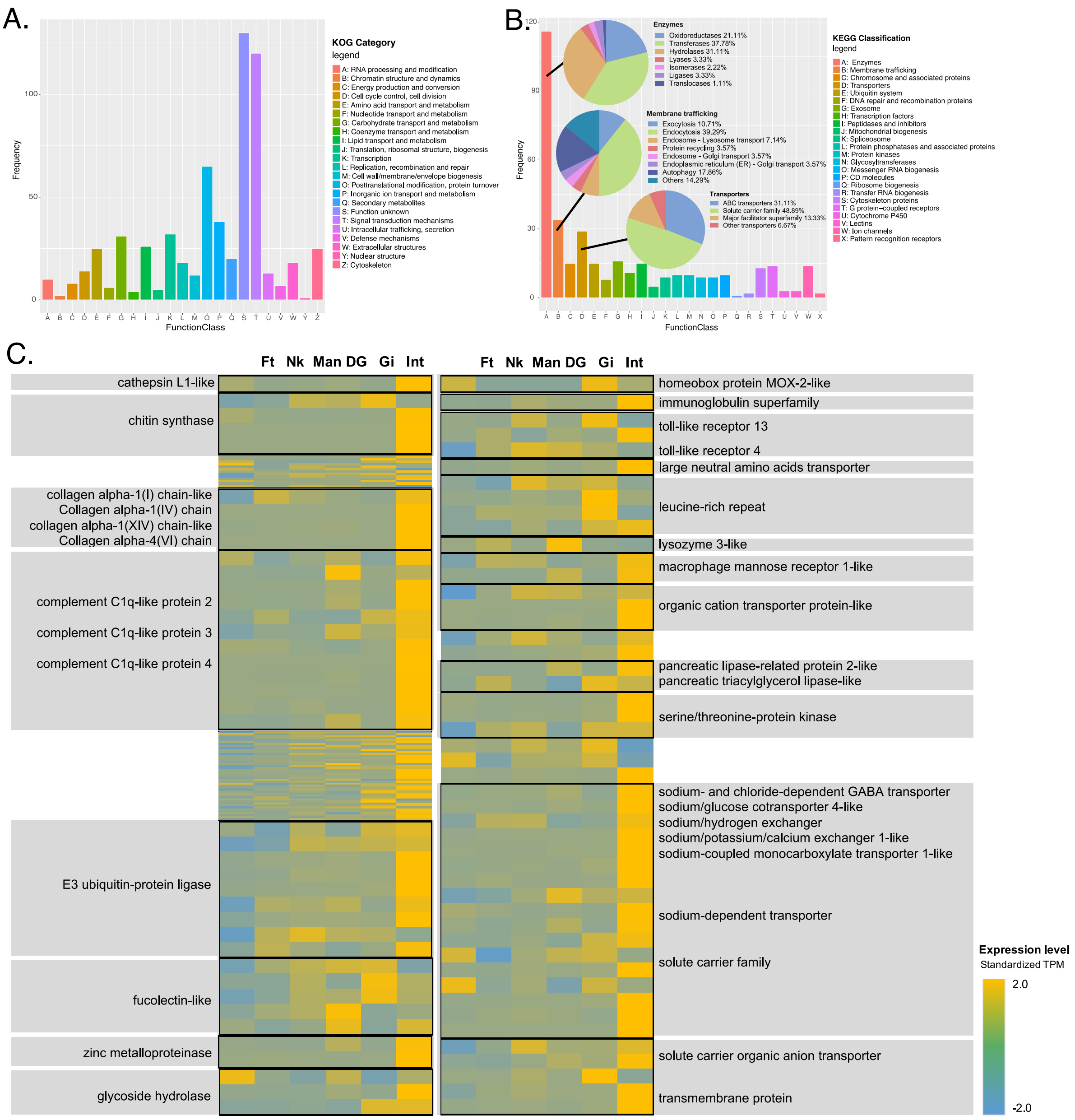

**Figure S2. Unique orthologous genes of the Wocan *Alviniconcha marisindica* based on genomic comparisons with the other three Lophotrochozoa.** Distribution of *Alviniconcha*-unique orthologous genes in different functional categories based on (**A**) KOG and (**B**) KEGG annotations. (**C**) Expression level of *Alviniconcha*-unique genes in the foot (Ft), neck (Nk), mantle (Man), digestive gland (DG), gill (Gi), and intestine (Int) tissues are shown in the heat map.

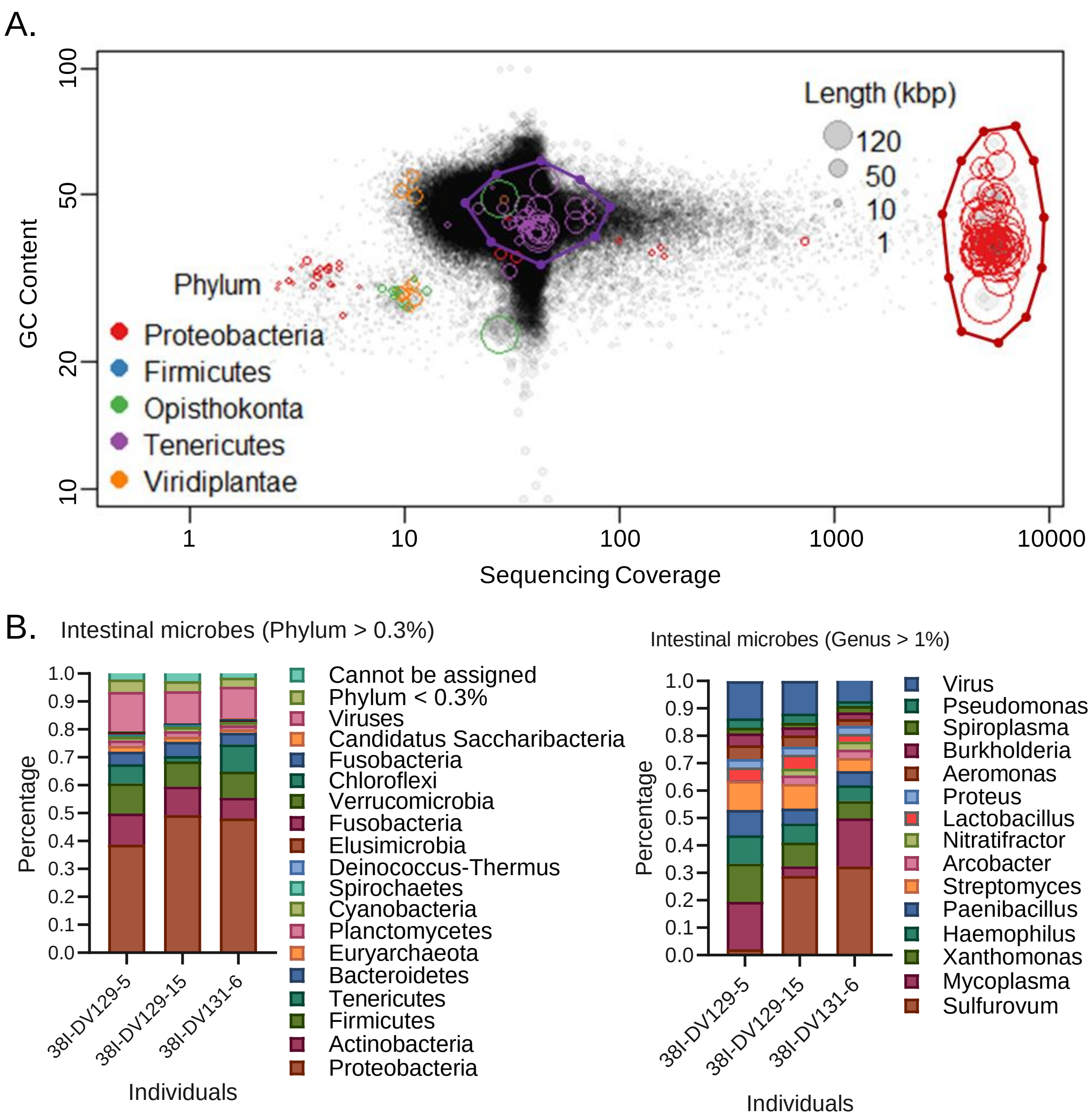

**Figure S3. Gill endosymbionts and intestinal ectosymbionts of *Alviniconcha marisindica* from the Wocan vent field.** (**A**) Differential coverage binning of *Alviniconcha* and its gill endosymbiont (one dominant phylotype, Campylobacterota). Each circle represents a contig and its area indicates the contig length. The colour represents the systematic affinity of the contig. The contigs of *A. marisindica* and its endosymbiont are grouped based on their different levels of GC content and sequencing coverage. The area of red circles represents one single dominant Campylobacterota and the purple represents one less abundant Mollicutes. (**B**) Abundance and community structure of the ectosymbionts in the intestine. Microbial taxonomic structures are deduced from the intestinal metagenomes. The community compositions are displayed at the phylum and genus level based on Kaiju classification. Phylum totaling >0.3% and genus totaling >1% of the samples are shown respectively.

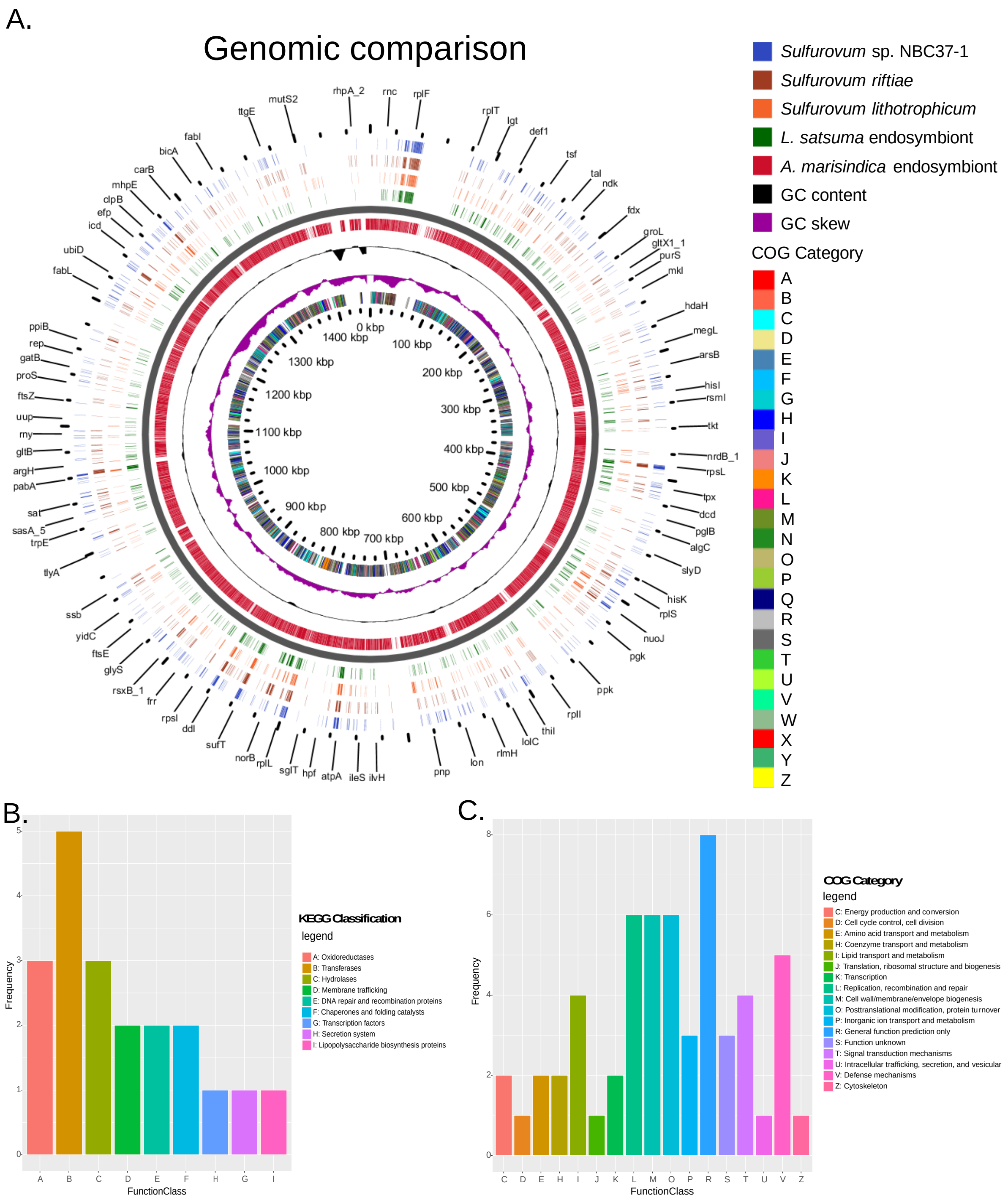

**Figure S4. Genomic comparisons and gene family analyses of the Wocan *Alviniconcha marisindica* endosymbiont and the other four campylobacterotal close relatives.** (A) Genome-similarity comparisons of the endosymbiont of *A. marisindica* and the other four Campylobacterota constructed using GView. From outside to the centre: genes of four Campylobacterota on forward strand, the innermost circular part represents the endosymbiont of *A. marisindica*, including genes on forward strand, GC content (%), GC skew. Colours indicate categories of clusters of orthologous groups (COGs) and the scale is in kbp. The identifiers of outside-to-inside rings are listed on the right. The unique orthologous genes of the *A. marisindica* endosymbiont are classified into different functional categories based on (**B**) KEGG and (**C**) COG annotations.

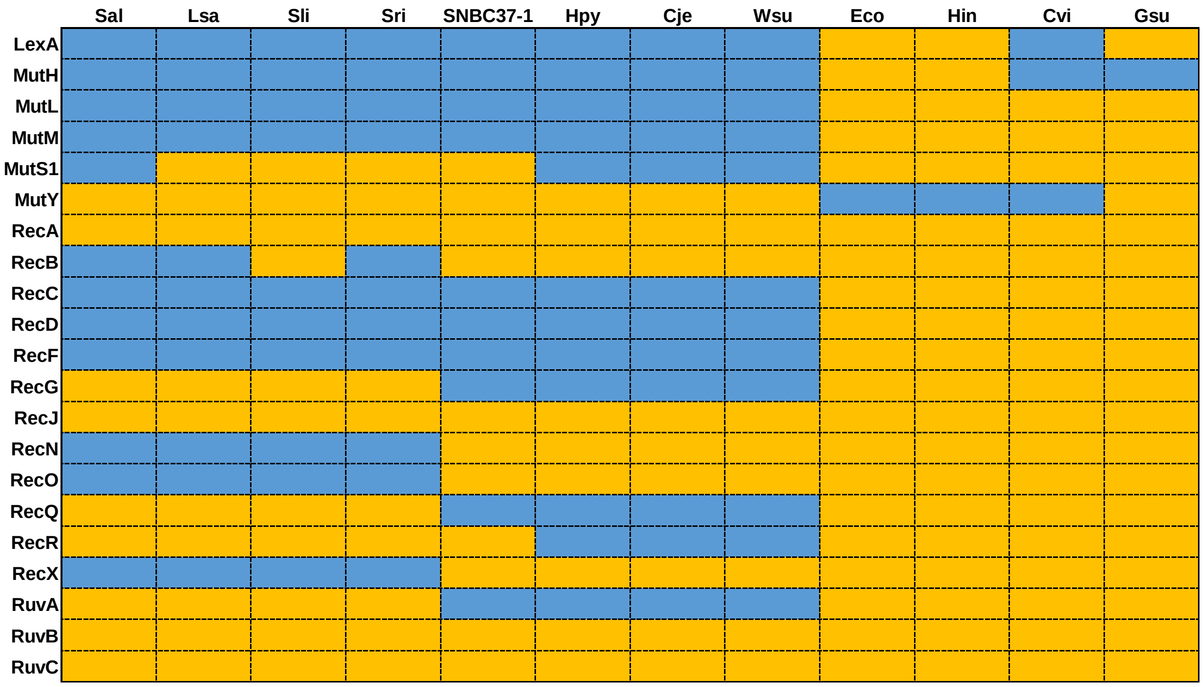

**Figure S5. DNA-repair genes in representative Proteobacteria.** Blue indicates absence, and yellow indicates presence. Sal, *Sulfurovum alviniconcha* CR; Lsa, the endosymbiont of *Lamellibrachia satsuma*; Sli, *Sulfurovum lithotrophicum*; Sri, *Sulfurovum riftiae*; SNBC37-1, *Sulfurovum* sp. NBC37-1; Hpy, *Helicobacter pylori* 26695; Cje, *Campylobacter jejuni* NCTC11168; Wsu, *Wolinella succinogenes* DSM1740; Eco, *Escherichia coli* K12 (γ-proteobacterium); Hin, *Haemophilus influenzae* Rd KW20 (γ-proteobacterium); Cvi, *Chromobacterium violaceum* ATCC12472 (β-proteobacterium); Gsu, *Geobacter sulfurreducens* PCA (δ-proteobacterium).

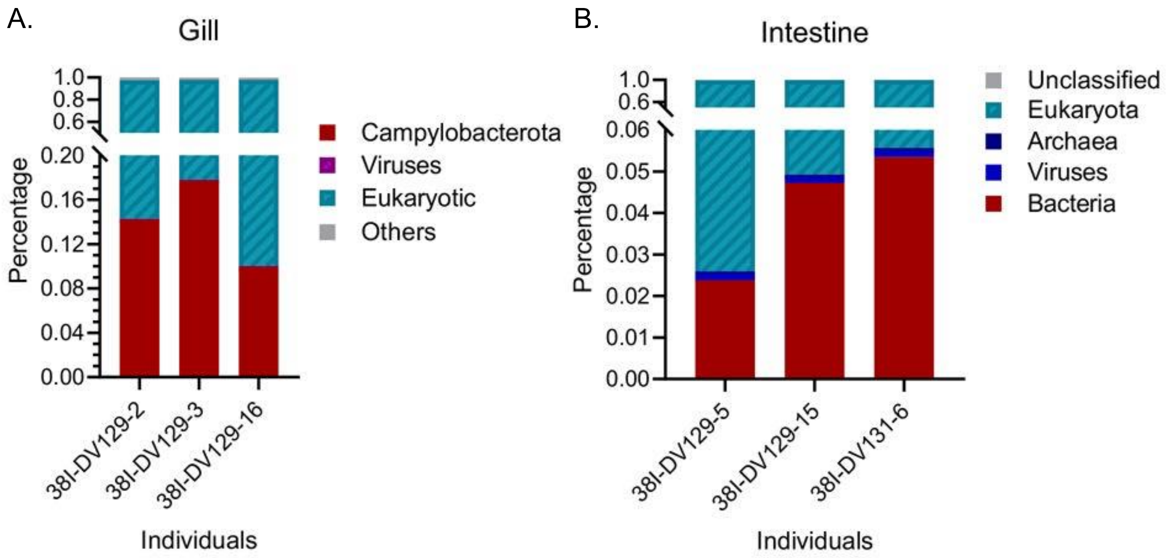

**Figure S6. The abundance of the symbionts in gill and intestine of snail *Alviniconcha marisindica*.** (**A**) The abundance of campylobacterotal endosymbionts in the gill metagenomes. (**B**) The abundance of ectosymbionts in the intestinal metagenomes.

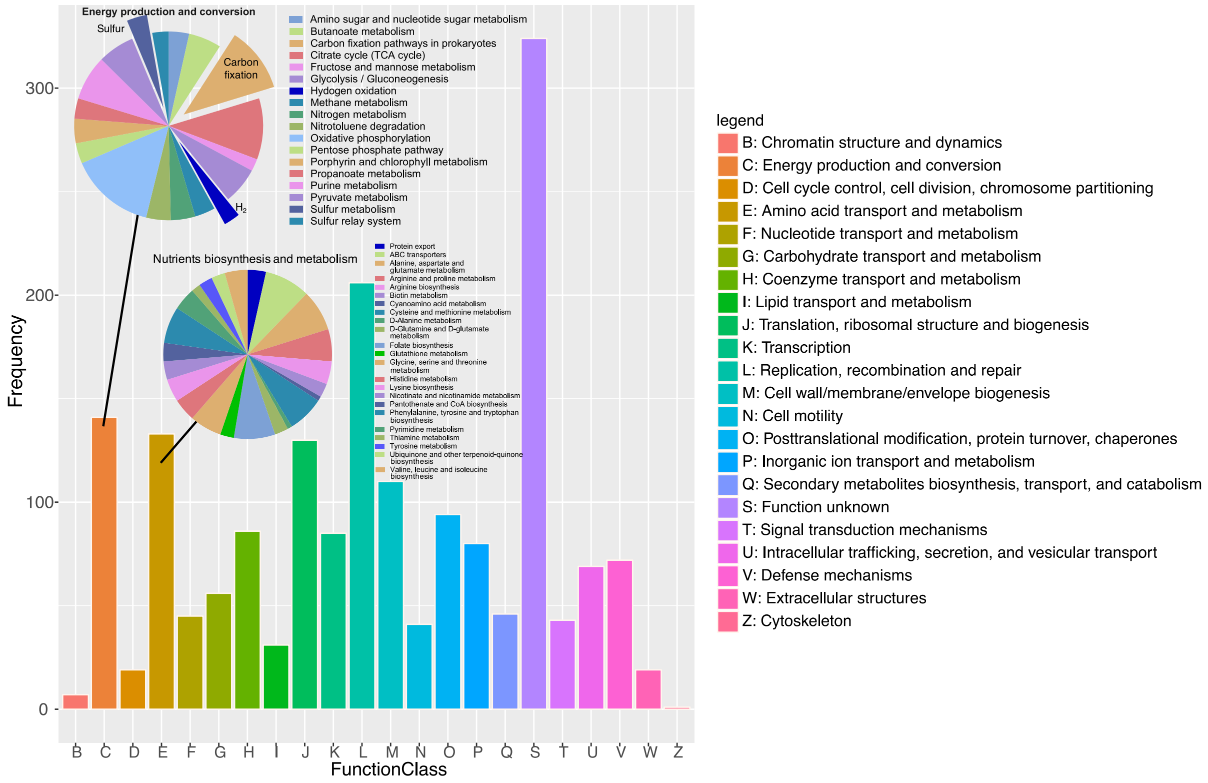

**Figure S7. Pan-genome of 23 *Alviniconcha marisindica* endosymbionts from the Wocan vent field.** Pan-genes are classified into different functional categories based on COG annotations. The number of genes in each category is shown in the histogram. In particular, genes involved in energy production and conversion, nutrient biosynthesis, and metabolism are classified into different metabolic pathways based on KEGG annotations. The percentage of genes involved in each type of metabolic pathway is shown in the pie chart.

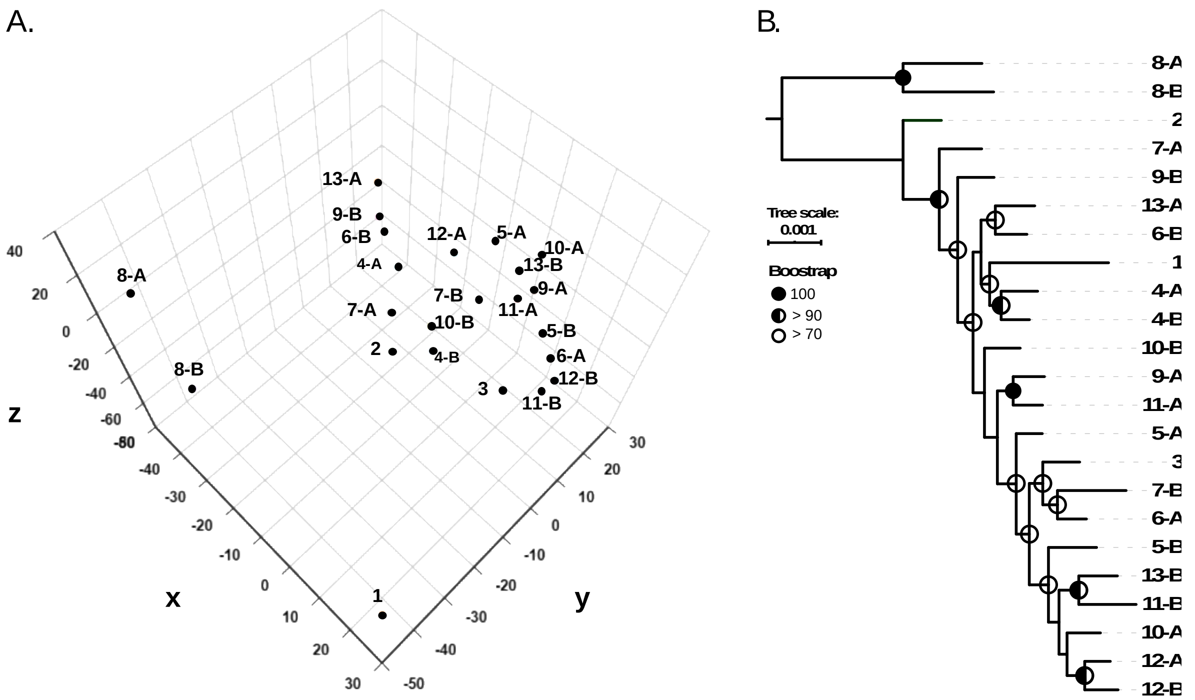

**Figure S8. Genome-based comparisons of 23 endosymbiotic isolates of *Alviniconcha marisindica* from the Wocan vent field.** (**A**) Principal component analysis under the BLOSUM62 model and (**B**) phylogenomic analysis on the orthologous proteins in 941 single-copy shared orthologues of 23 endosymbionts from 13 *A. marisindica* individuals.

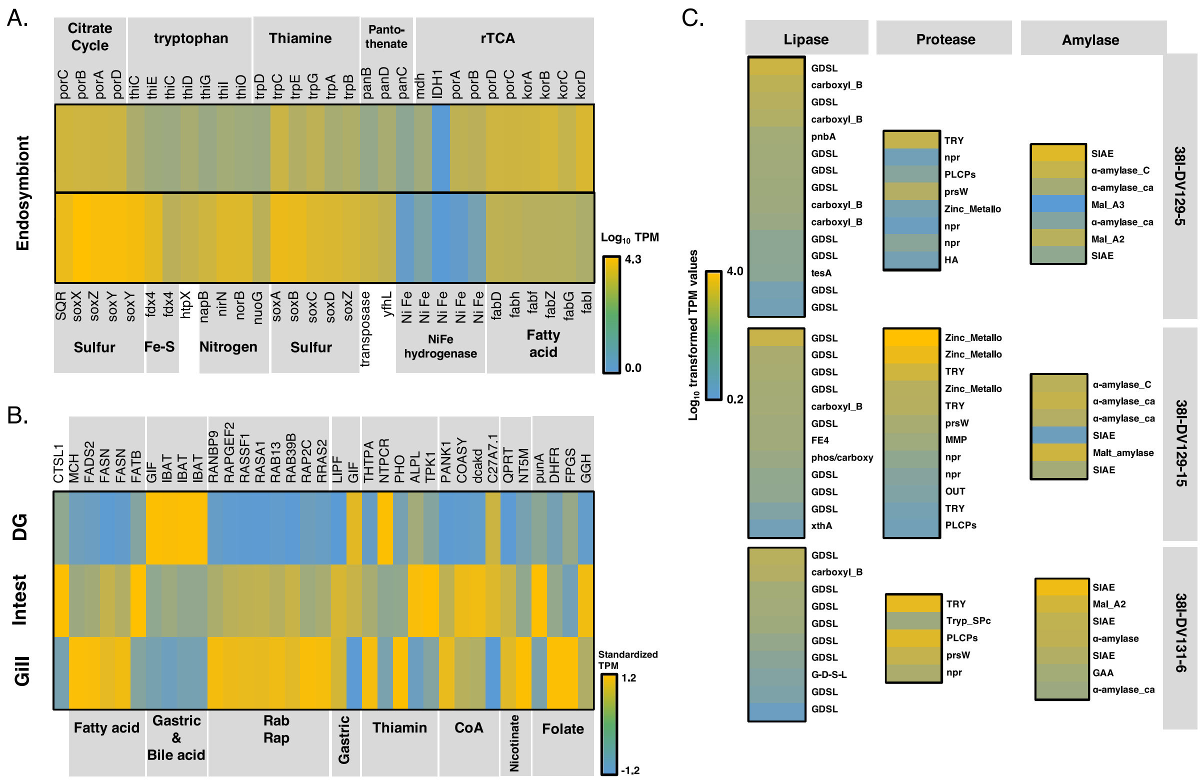

**Figure S9. Transcriptional activity of genes participating in tripartite syntrophic interactions in the Wocan *Alviniconcha marisindica* holobiont.** Expression level of genes that participate in (**A**) chemoautotroph, nutrient biosynthesis and metabolism, and nitrogen and carbon metabolism in the endosymbiont of *A. marisindica*, (**B**) nutrient biosynthesis and food digestion in the *A. marisindica* host, and (**C**) gut bacterial exo-hydrolase biosynthesis for intestinal food digestion in the *A. marisindica* host for three *A. marisindica* individuals.

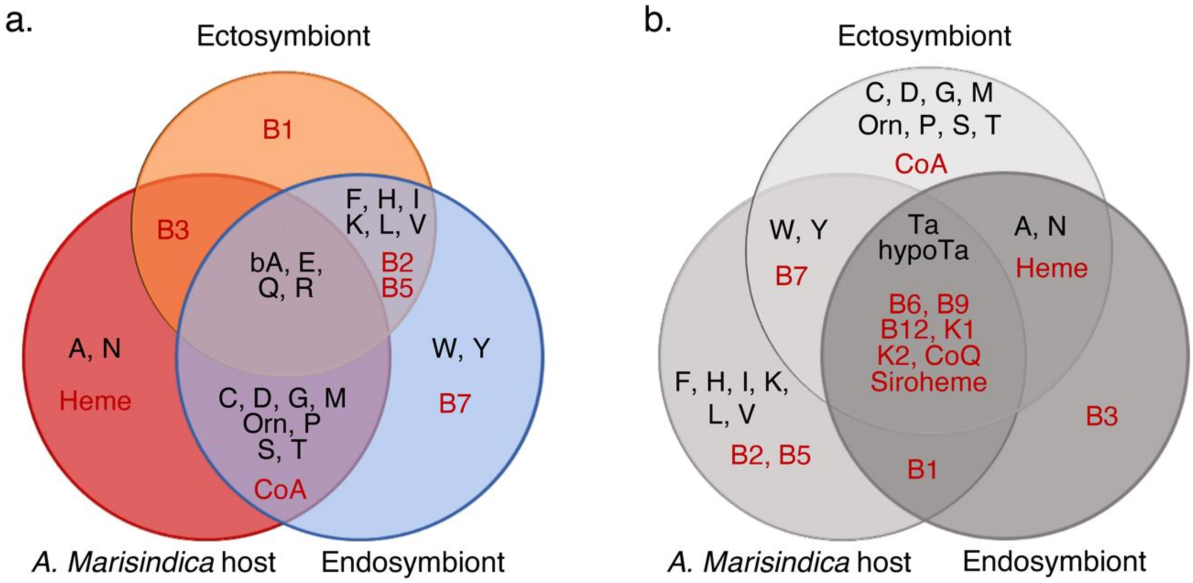

**Figure S10. Nutrient biosynthesis capability of the Wocan *Alviniconcha marisindica* holobiont.** Venn diagram showing the nutrients **(A)** with or **(B)** without complete biosynthesis pathways in the genomes of *A. marisindica* and its symbionts. Amino acids (black colour): A – Alanine, bA – β-Alanine, C – Cysteine, D – Aspartate (aspartic acid), E – Glutamic acid, F – Phenylalanine, G – Glycine, H – Histidine, hypoTa – Hypotaurine, I – Isoleucine, K – Lysine, L – Leucine, M – Methionine, N – Asparagine, Orn – Ornithine, P – Proline, Q – Glutamine, R – Arginine, S – Serine, T – Threonine, Ta – Taurine, V – Valine, W – Tryptophan, Y – Tyrosine; Vitamins/cofactors (red colour): B1 – Thiamin, B2 – Riboflavin, B3 – Nicotinate and nicotinamide, B5 – Pantothenate, B6 – Pyridoxine, B7 – Biotin, B9 – Folate, B12 – Cobalamin, K1 – Phylloquinone, K2 – Menaquinone, CoA – Coenzyme A, CoQ – Coenzyme Q

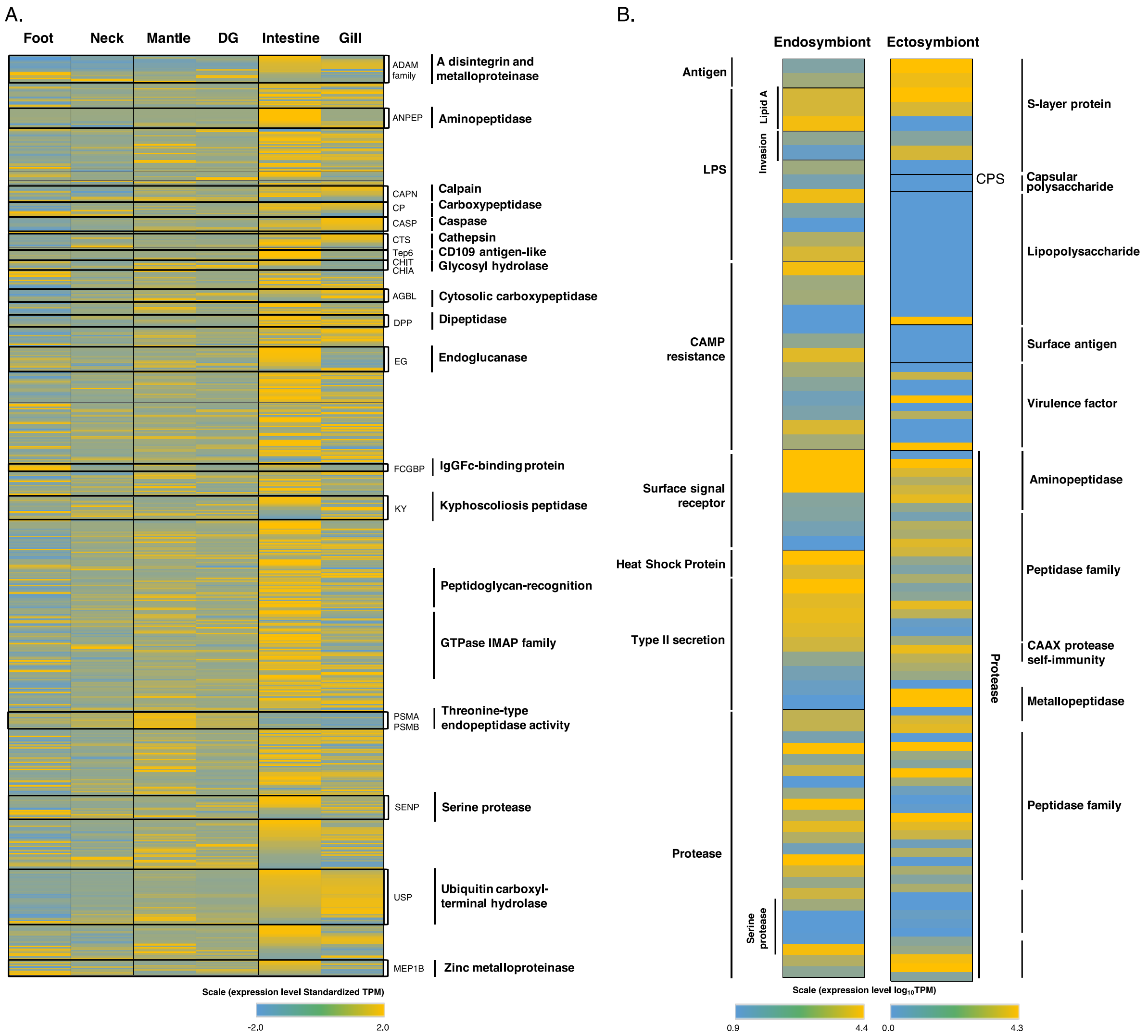

**Figure S11. Transcriptional activity of genes participating in the nutrient digestion and absorption of the Wocan *Alviniconcha marisindica* host and bacterial immune evasion and defence.** Heat map of transcriptional activity of genes that (**A**) encode various hydrolases, including proteases, glycoside hydrolase, and peptidoglycan recognition proteins (PGRPs) in the foot, neck, mantle, digestive gland (DG), intestine, and gill tissues, (**B**) participate in bacterial surface-associated virulence factors, surface modification, protease synthesis, and secretion in the gill endosymbiont and the intestinal ectosymbionts. Each grid in the heat map represents a gene identified in the respective sample. The colour represents the gene expression level (based on normalised TPM values). The annotated gene names and their functional classifications are listed on the sides.

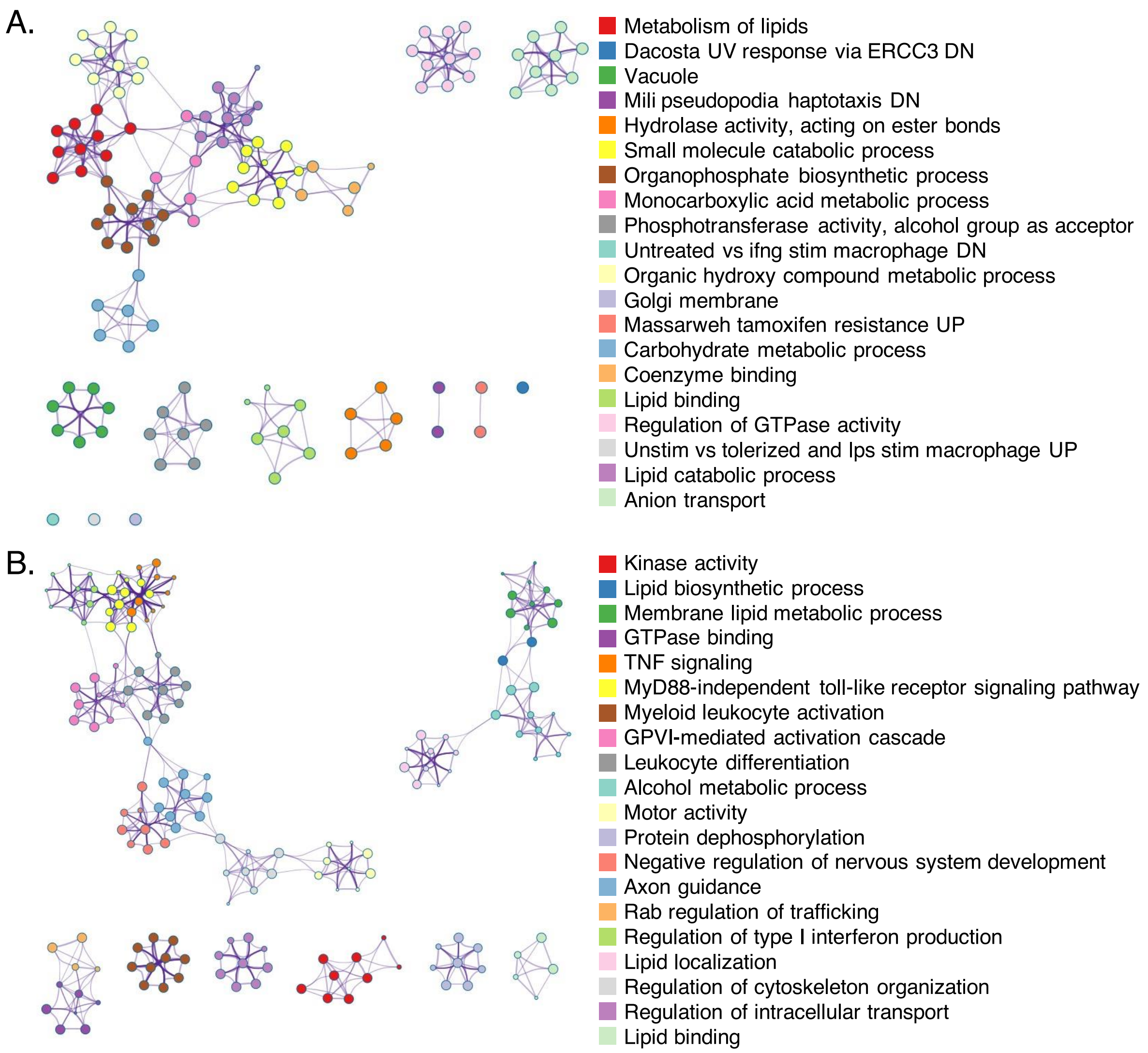

**Figure S12. Gene Ontology (GO) enrichment network of differentially expressed genes (DEGs).** The significantly (p-value < 0.01) enriched GO terms of selected highly expressed genes in the (**A**) intestine and (**B**) gills of *Alviniconcha marisindica* are clustered according to their functional category. The connecting pairs of nodes showing the intra-cluster and inter-cluster similarities of enriched terms. The colour code represents different cluster annotations. Each node represents an enriched term.

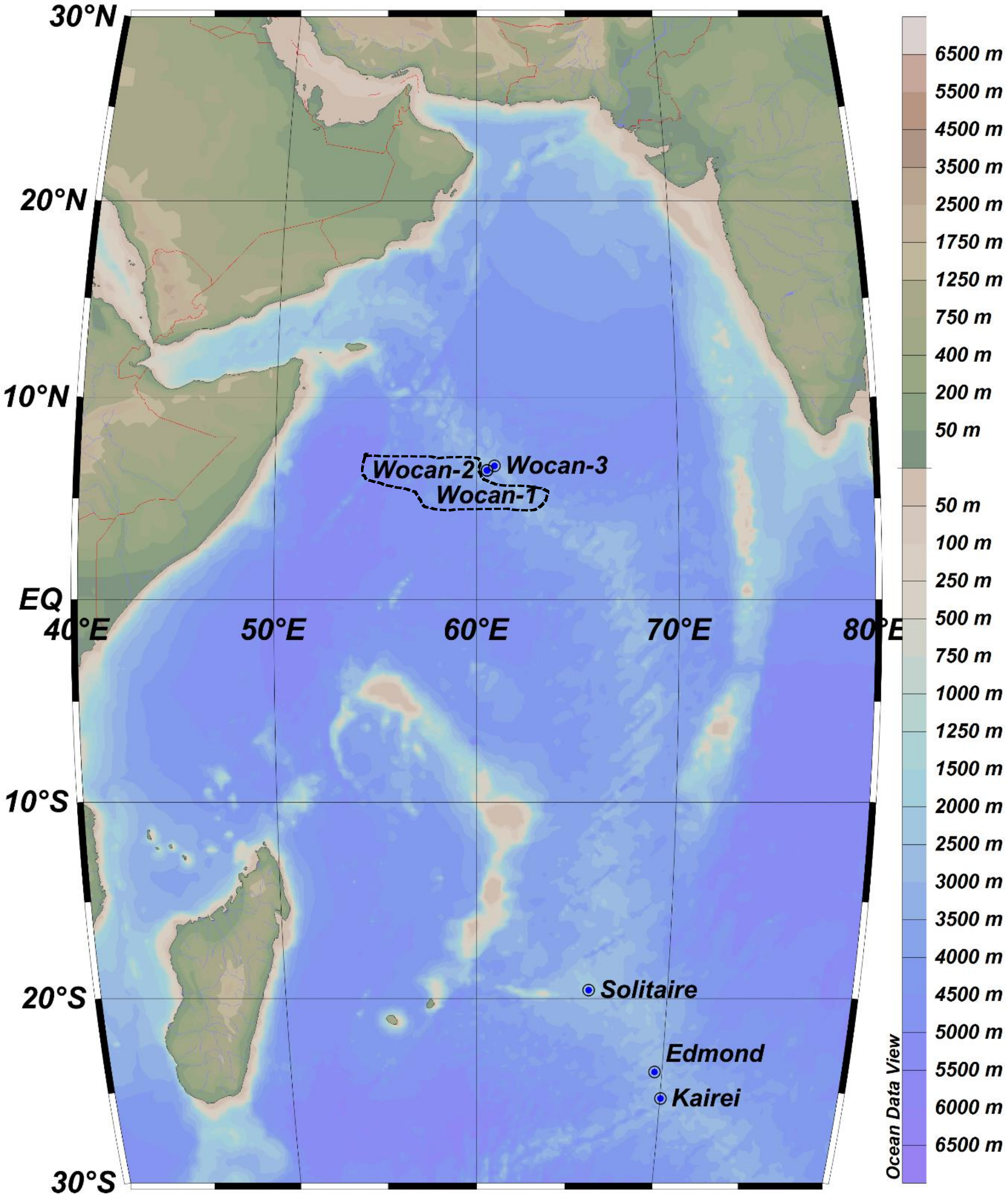

**Figure S13. Bathymetric map of the hydrothermal vents over Central Indian Ridge and Carlsberg Ridge.** The location of the Wocan Hydrothermal Field (WHF) in this study is on Carlsberg Ridge and marked as one point 'Wocan-3'. 'Wocan-1' and 'Wocan-2' are found by DY 24th cruise in 2012, and marked as one point. The three vent sites of 'Solitaire', 'Edmond', and 'Kairei' are marked as three points which located on Central Indian Ridge.

**Supplementary Tables**

**Table S1.** The BUSCO score for different version of the assembly.

| Version | BUSCO score |
|---|---|
| **After 1 round of Pilon correction** | C:90.1% [S:88.7%,D:1.4%], F:1.2%, M:8.7%, n:954 |
| **After 2 rounds of Pilon correction** | C:95.6% [S:94.2%,D:1.4%], F:0.8%, M:3.6%, n:954 |
| **After Redundans** | **C:96.5%** [S:95.9%,D:0.6%], F:1.2%, **M:2.3%**, n:954 |

**Table S2.** Quast genome assembly assessment report of *Alviniconcha marisindica*.

| Assembly parameters | Assessment results |
|---|---|
| **No. of contigs (>= 5000 bp)** | 3,926 |
| **No. of contigs (>= 10000 bp)** | 2,814 |
| **No. of contigs (>= 25000 bp)** | 2,097 |
| **No. of contigs (>= 50000 bp)** | 1,762 |
| **Total length (>= 5000 bp)** | 829,612,046 |
| **Total length (>= 10000 bp)** | 821,908,196 |
| **Total length (>= 25000 bp)** | 810,769,599 |
| **Total length (>= 50000 bp)** | 798,818,352 |
| **No. of contigs** | 3,926 |
| **Largest contig (bp)** | 4,734,527 |
| **Total length (bp)** | 829,612,046 |
| **GC (%)** | 45.47 |
| **N50** | 727,552 |
| **N75** | 366972 |
| **L50** | 336 |
| **L75** | 743 |
| **Total reads** | 17,708,417 |
| **Mapped (%)** | 98.72 |
| **Avg. coverage depth** | 81 |
| **Coverage >= 1X (%)** | 99.40 |
| **Coverage >= 5X (%)** | 99.01 |
| **Coverage >= 10X (%)** | 99.18 |
| **N's per 100 kbp** | 0.02 |

**Table S3.** The annotation and expression levels of genes involved in encoding representative exo-hydrolases in intestinal flora of three *Alviniconcha marisindica* individuals.

**Individual 38I-DV129-5**

| **Gene_ID** | **Annotation** | **Gene** | **COG** | **Best Tax-Level** | **TPM** |
|---|---|---|---|---|---|
| **k141_240474_2** | sialate O-acetylesterase-like | | G | Cytophagia | 283.14 |
| **k141_338559_2** | sialate O-acetylesterase | | G | Leeuwenhoekiella | 122.07 |
| **k141_402846_1** | sialate O-acetylesterase-like | | G | Sphingobacteriia | 61.69 |
| **k141_216833_3** | Alpha amylase, catalytic domain | treY | G | Microbacteriaceae | 490.45 |
| **k141_34583_1** | Alpha-amylase domain | | G | Sphingomonadales | 304.57 |
| **k141_698499_2** | maltase A2-like | | G | Lactobacillaceae | 328.87 |
| **k141_698499_2** | maltase A2-like | | G | Lactobacillaceae | 702.71 |
| **k141_240474_2** | sialate O-acetylesterase-like | | G | Cytophagia | 262.29 |
| **k141_338559_2** | sialate O-acetylesterase | | G | Leeuwenhoekiella | 112.56 |
| **k141_402846_1** | sialate O-acetylesterase-like | | G | Sphingobacteriia | 4050.85 |
| **k141_240474_2** | sialate O-acetylesterase-like | | G | Cytophagia | 262.29 |
| **k141_216833_3** | Alpha amylase, catalytic domain | treY | G | Microbacteriaceae | 30.38 |
| **k141_295070_4** | alpha-glucosidase | aglA | G | Myxococcales | 46.35 |
| **k141_34583_1** | Alpha-amylase domain | | G | Sphingomonadales | 232.23 |
| **k141_188836_2** | GDSL-like Lipase/Acylhydrolase family | | E | Cytophagia | 18.91 |
| **k141_283355_1** | GDSL-like Lipase/Acylhydrolase family | | E | Cytophagia | 1.65 |
| **k141_513053_1** | GDSL-like Lipase/Acylhydrolase family | tesA | E | Leeuwenhoekiella | 36.49 |
| **k141_60153_1** | GDSL-like Lipase/Acylhydrolase | | E | Micromonosporales | 45.59 |
| **k141_479102_1** | PFAM GDSL-like Lipase Acylhydrolase | | E | Nostocales | 1.55 |
| **k141_299134_1** | GDSL-like Lipase/Acylhydrolase family | | E | Proteobacteria | 10.65 |
| **k141_747374_1** | G-D-S-L family lipolytic protein | | E | Proteobacteria | 6.50 |
| **k141_435642_1** | GDSL-like Lipase/Acylhydrolase family | | E | Sphingomonadales | 156.20 |
| **k141_572192_2** | GDSL-like Lipase/Acylhydrolase family | | E | Sphingomonadales | 43.11 |
| **k141_72843_1** | GDSL-like Lipase/Acylhydrolase family | | E | Streptosporangiales | 4.95 |
| **k141_499802_1** | carboxylesterase 3B | lipT | I | Gordoniaceae | 383.44 |
| **k141_344000_3** | para-nitrobenzyl esterase | pnbA | I | Paenibacillaceae | 1.55 |
| **k141_219741_3** | esterase E4-like | | I | Sphingomonadales | 114.20 |
| **k141_175913_1** | Exodeoxyribonuclease III | | L | Bacteria | 98.89 |
| **k141_113311_7** | Exodeoxyribonuclease III | | L | Bacteria | 47.97 |
| **k141_8172_1** | Exodeoxyribonuclease III | | L | Bacteria | 4.49 |
| **k141_40481_1** | Exodeoxyribonuclease III | | S | Proteobacteria | 7.73 |
| **k141_262990_1** | Exodeoxyribonuclease III | holA | L | Tenericutes | 608.82 |
| **k141_175913_1** | Exodeoxyribonuclease III | | L | Bacteria | 98.89 |
| **k141_39157_1** | Trypsin | | E | Corynebacteriaceae | 2420.31 |
| **k141_791929_1** | Trypsin-like serine protease | | M | Vibrionales | 33.11 |

| | | | | | |
|---|---|---|---|---|---|
| **k141_39157_1** | Trypsin | | E | Corynebacteriaceae | 2420.31 |
| **k141_562146_2** | neutral protease-like | lasB | Q | Colwelliaceae | 2.96 |
| **k141_541590_1** | neutral protease-like | lasB | E | Vibrionales | 75.77 |
| **k141_562146_2** | neutral protease-like | lasB | Q | Colwelliaceae | 2.96 |
| **k141_46506_1** | Papain-like cysteine protease AvrRpt2 | | H | Bacillus | 1557.63 |
| **k141_46506_1** | Papain-like cysteine protease AvrRpt2 | | H | Bacillus | 1557.63 |
| **k141_824392_1** | hemagglutinin | lasB | E | Shewanellaceae | 1.56 |
| **k141_776239_3** | Protease prsW family | | S | Dermatophilaceae | 305.60 |
| **Individual 38I-DV129-15** | | | | | |
| **k141_688074_2** | sialate O-acetylesterase-like | | G | Cytophagia | 1.91 |
| **k141_543608_2** | sialate O-acetylesterase | | G | Leeuwenhoekiella | 47.14 |
| **k141_42238_1** | SMART alpha amylase catalytic sub domain | | G | Chloroflexi | 324.57 |
| **k141_352388_1** | SMART alpha amylase catalytic sub domain | | G | Chloroflexi | 115.94 |
| **k141_819448_3** | Maltogenic Amylase, C-terminal domain | treS | G | Rhizobiaceae | 534.59 |
| **k141_236157_3** | 1,4-alpha-glucan-branching enzyme | GLC3 | G | Chaetomiaceae | 178.97 |
| **k141_908504_2** | GDSL-like Lipase/Acylhydrolase | | E | Alteromonadaceae | 16.04 |
| **k141_660256_1** | GDSL-like Lipase/Acylhydrolase family | ypmR | E | Carnobacteriaceae | 51.37 |
| **k141_318019_1** | GDSL-like Lipase/Acylhydrolase family | | E | Cytophagia | 0.76 |
| **k141_672619_1** | GDSL-like Lipase/Acylhydrolase family | | E | Flavobacteriia | 6.04 |
| **k141_366024_1** | GDSL-like Lipase/Acylhydrolase family | tesA | E | Leeuwenhoekiella | 0.97 |
| **k141_637991_1** | GDSL-like Lipase/Acylhydrolase | | E | Micromonosporales | 372.32 |
| **k141_519721_1** | GDSL-like Lipase/Acylhydrolase | | E | Micromonosporales | 66.99 |
| **k141_781927_1** | GDSL-like Lipase/Acylhydrolase | | E | Micromonosporales | 59.82 |
| **k141_39007_1** | PFAM GDSL-like Lipase Acylhydrolase | | E | Nostocales | 8.19 |
| **k141_640620_1** | PFAM GDSL-like Lipase Acylhydrolase | | E | Nostocales | 2.14 |
| **k141_353962_1** | GDSL-like Lipase/Acylhydrolase family | | S | Pleosporales | 15.63 |
| **k141_302376_2** | GDSL-like Lipase/Acylhydrolase | | E | Porphyromonadaceae | 3.02 |
| **k141_319940_1** | GDSL-like Lipase/Acylhydrolase family | | E | Sphingomonadales | 39.71 |
| **k141_368234_1** | GDSL-like Lipase/Acylhydrolase family | | E | Streptosporangiales | 1.30 |
| **k141_908504_2** | GDSL-like Lipase/Acylhydrolase | | E | Alteromonadaceae | 16.04 |
| **k141_463823_1** | Esterase FE4 | | G | Ascomycota | 39.42 |
| **k141_542556_1** | Exodeoxyribonuclease III | | G | Ascomycota | 2.70 |
| **k141_664301_1** | Belongs to the type-B carboxylesterase lipase family | | I | Eurotiales | 48.53 |

| | | | | | |
|---|---|---|---|---|---|
| **k141_238923_1** | carboxylesterase 3B | lipT | I | Gordoniaceae | 83.59 |
| **k141_712297_1** | Exodeoxyribonuclease III | | | Agaricomycetes incertae sedis | 0.75 |
| **k141_542556_1** | Exodeoxyribonuclease III | | G | Ascomycota | 2.70 |
| **k141_621930_1** | Exodeoxyribonuclease III | | L | Bacteria | 110.90 |
| **k141_611192_1** | Exodeoxyribonuclease III | | L | Bacteria | 13.32 |
| **k141_49768_1** | Exodeoxyribonuclease III | | L | Bacteria | 6.64 |
| **k141_489964_2** | Exodeoxyribonuclease III | | L | Bacteria | 4.03 |
| **k141_322147_1** | Exodeoxyribonuclease III | ypmS | S | Listeriaceae | 19.94 |
| **k141_448732_3** | Exodeoxyribonuclease III | exoA | L | Neisseriales | 3.38 |
| **k141_560323_2** | Exodeoxyribonuclease III | | S | Proteobacteria | 405.28 |
| **k141_711647_1** | Exodeoxyribonuclease III | NAR1 | Y | Taphrinomycotina | 22.07 |
| **k141_396224_1** | fibrinolytic enzyme, isozyme C-like | | O | Actinobacteria | 139.91 |
| **k141_246661_1** | chymotrypsin-like serine proteinase | | O | Actinobacteria | 3.24 |
| **k141_2041_1** | Trypsin | | E | Corynebacteriaceae | 613.73 |
| **k141_396224_1** | fibrinolytic enzyme, isozyme C-like | | O | Actinobacteria | 139.91 |
| **k141_246661_1** | chymotrypsin-like serine proteinase | | O | Actinobacteria | 3.24 |
| **k141_686364_2** | neutral protease-like | lasB | Q | Colwelliaceae | 0.40 |
| **k141_462614_1** | neutral protease-like | lasB | E | Gammaproteobacte ria | 8.32 |
| **k141_148422_1** | neutral protease-like | lasB | E | Vibrionales | 15.26 |
| **k141_462614_1** | neutral protease-like | lasB | E | Gammaproteobacte ria | 8.32 |
| **k141_825267_1** | Papain-like cysteine protease AvrRpt2 | | S | Clostridia | 4.42 |
| **k141_605305_1** | Papain-like cysteine protease AvrRpt2 | | S | Clostridia | 2.49 |
| **k141_534720_1** | Papain-like cysteine protease AvrRpt2 | | S | Clostridia | 1.73 |
| **k141_470447_1** | Papain-like cysteine protease AvrRpt2 | | S | Clostridia | 1.10 |
| **k141_726180_1** | Papain-like cysteine protease AvrRpt2 | | S | Clostridia | 1.06 |
| **k141_686583_1** | Papain-like cysteine protease AvrRpt2 | | S | Clostridia | 0.87 |
| **k141_417641_4** | Protease prsW family | | S | Dermatophilaceae | 64.06 |
| **k141_890620_1** | matrilysin family metalloendoprotease | | O | Proteobacteria | 35.72 |
| **k141_847292_1** | OTU-like cysteine protease | | OT | Eurotiales | 5.79 |
| **Individual 38I-DV131-6** | | | | | |
| **k141_623736_1** | sialate O-acetylesterase | | G | Leeuwenhoekiella | 27.19 |
| **k141_469403_1** | sialate O-acetylesterase-like | | G | Sphingobacteriia | 2004.59 |
| **k141_366488_2** | SMART alpha amylase catalytic sub domain | | G | Chloroflexi | 85.31 |
| **k141_758050_2** | maltase A2-like | | G | Lactobacillaceae | 226.31 |
| **k141_513093_1** | maltase A3-like | | G | Leuconostocaceae | 1.54 |
| **k141_469403_1** | sialate O-acetylesterase-like | | G | Sphingobacteriia | 2004.59 |
| **k141_461237_1** | alpha-glucosidase | malZ | G | Oceanospirillales | 14.50 |

| | | | | | |
|---|---|---|---|---|---|
| **k141_474653_5** | 1,4-alpha-glucan-branching enzyme | GLC3 | G | Chaetomiaceae | 466.84 |
| **k141_366488_2** | SMART alpha amylase catalytic sub domain | | G | Chloroflexi | 85.31 |
| **k141_511492_1** | GDSL-like Lipase/Acylhydrolase | | E | Alteromonadaceae | 12.12 |
| **k141_290837_1** | GDSL-like Lipase/Acylhydrolase | | E | Clostridia | 2.77 |
| **k141_454472_2** | Lysophospholipase L1 | | E | Cytophagia | 11.80 |
| **k141_652332_1** | hydrolase GDSL | | E | Cytophagia | 3.40 |
| **k141_566963_1** | GDSL-like Lipase/Acylhydrolase family | | E | Flavobacteriia | 41.46 |
| **k141_23211_1** | GDSL-like Lipase/Acylhydrolase family | | E | Hyphomonadaceae | 484.00 |
| **k141_483011_1** | GDSL-like Lipase/Acylhydrolase | | E | Micromonosporales | 152.28 |
| **k141_445749_1** | GDSL-like Lipase/Acylhydrolase | | E | Micromonosporales | 40.19 |
| **k141_599789_1** | GDSL-like Lipase/Acylhydrolase | estA | E | Sphingobacteriia | 12.80 |
| **k141_553159_2** | GDSL-like Lipase/Acylhydrolase family | | E | Sphingomonadales | 37.03 |
| **k141_546352_1** | G-D-S-L family lipolytic protein | | E | Sphingomonadales | 4.12 |
| **k141_511492_1** | GDSL-like Lipase/Acylhydrolase | | E | Alteromonadaceae | 12.12 |
| **k141_378516_2** | Belongs to the type-B carboxylesterase lipase family | | I | Actinobacteria | 32.96 |
| **k141_776825_1** | para-nitrobenzyl esterase | pnbA | I | Bacillus | 45.32 |
| **k141_564754_1** | Belongs to the type-B carboxylesterase lipase family | | I | Eurotiales | 107.07 |
| **k141_568994_1** | carboxylesterase 3B | lipT | I | Gordoniaceae | 360.87 |
| **k141_785481_1** | Belongs to the type-B carboxylesterase lipase family | lipT | I | Mycobacteriaceae | 209.83 |
| **k141_166509_1** | Belongs to the type-B carboxylesterase lipase family | | T | Nectriaceae | 26.71 |
| **k141_378516_2** | Belongs to the type-B carboxylesterase lipase family | | I | Actinobacteria | 32.96 |
| **k141_62324_7** | Exodeoxyribonuclease III | exoA | L | Alteromonadaceae | 12.69 |
| **k141_603633_1** | Exodeoxyribonuclease III | | L | Bacteria | 9.79 |
| **k141_354178_2** | Exodeoxyribonuclease III | | L | Bacteria | 6.34 |
| **k141_199648_2** | Exodeoxyribonuclease III | | L | Bacteria | 5.53 |
| **k141_25149_1** | Exodeoxyribonuclease III | ypmS | S | Listeriaceae | 80.29 |
| **k141_46566_1** | Exodeoxyribonuclease III | | S | Proteobacteria | 3.20 |
| **k141_283492_1** | Exodeoxyribonuclease III | NAR1 | Y | Taphrinomycotina | 32.33 |
| **k141_281926_1** | Exodeoxyribonuclease III | holA | L | Tenericutes | 108.10 |
| **k141_62324_7** | Exodeoxyribonuclease III | exoA | L | Alteromonadaceae | 12.69 |
| **k141_31145_1** | Trypsin | | E | Corynebacteriaceae | 303.71 |
| **k141_111587_4** | neutral protease-like | lasB | Q | Colwelliaceae | 2.54 |
| **k141_385692_1** | neutral protease-like | lasB | E | Gammaproteobacteria | 1.76 |
| **k141_588201_1** | neutral protease-like | lasB | E | Vibrionales | 10.02 |
| **k141_111587_4** | neutral protease-like | lasB | Q | Colwelliaceae | 2.54 |

| | | | | |
|---|---|---|---|---|
| **k141_46789_4** | Papain-like cysteine protease AvrRpt2 | H | Bacillus | 8.81 |
| **k141_445402_1** | peptidase domain protein | S | Clostridia | 1.71 |
| **k141_660310_1** | Papain-like cysteine protease AvrRpt2 | S | Clostridia | 0.74 |
| **k141_534044_1** | Papain-like cysteine protease AvrRpt2 | S | Clostridia | 0.68 |
| **k141_488656_1** | Papain-like cysteine protease AvrRpt2 | S | Clostridia | 0.65 |
| **k141_774423_1** | Papain-like cysteine protease AvrRpt2 | S | Clostridia | 0.44 |
| **k141_46789_4** | Papain-like cysteine protease AvrRpt2 | H | Bacillus | 8.81 |
| **k141_759841_8** | Protease prsW family | S | Dermatophilaceae | 113.86 |

### Supplementary Notes

#### Supplementary Note 1

**Sample Dissection.** The frozen snails were thawed in RNAlater® (Invitrogen, USA) on ice, dissected with different tissues fixed separately in RNAlater®, and then prepared for nucleic acid extraction. A single specimen of *Alviniconcha marisindica* from Wocan was used for the holobiont genome assembly. A total of 16 individuals were dissected, and tissues/organs were collected for nucleic acid preparation and sequencing. DNA extracted from muscle of foot and neck from one male individual was used for host genome sequencing and assembly. DNA extractied from the endosymbiont-harbouring gills were used for endosymbiont genome sequencing and assembly. Three snail individuals, including the one used for identifying the host genome, were dissected into 7–10 tissue types each with RNA extraction performed on the different tissues, namely cephalic tentacle (Ct), digestive gland (DG), gonad (testis or ovary, Go), foot (Ft), endosymbiont-containing ctenidium (Gi), ectosymbiont-containing intestine (Int), mantle (Man), mantle edge (ME), neck furrow (NF), nephridium (Ne), and ventricle heart (VH), following the previously published anatomy of *Alviniconcha marisindica* from the Kairei deep-sea hydrothermal field (see Fig. 1 in Suzuki et al., 2005). Moreover, endosymbiont-containing ctenidium (gill, Gi) of several individuals were divided into different parts, including gill base (GB), gill distal (GD), gill posterior (GP) and gill anterior (GA). The gills of 10 individuals were divided into anterior and posterior parts, and the DNA of these 20 parts were extracted separately for metagenome sequencing. The intestines of the three individuals were dissected for total DNA and RNA extraction for the gut microbiome, and the gills of these individuals were also dissected for total RNA extraction.

**Nucleic Acid Preparation.** Genomic DNA (gDNA) was extracted using the E.Z.N.A.® Mollusc DNA Kit (Omega Bio-tek, Georgia, USA) and then purified using Genomic DNA Clean & ConcentratorTM-10 Kit (Zymo Research, CA, USA) according to the manufacturer’s protocol. Total DNA of the gills and that of the intestines were extracted using the same protocol. Total RNA was extracted using Trizol (Invitrogen, USA) from different tissues following the manufacturer’s protocol and prepared for RNA-Seq. Nucleic acid quality was evaluated using agarose gel electrophoresis and a BioDrop µLITE (BioDrop, Holliston, MA, US), and nucleic acid concentrations were quantified using a Qubit fluorometer v3.0 (Thermo Fisher Scientific, Singapore).

**Morphological Observation.** The external morphology of a complete *Alviniconcha* individual preserved in absolute ethanol was shown in Figure S14, the internal morphology was observed under a Leica MZ9.5 stereozoom microscope, and the snails were dissected to isolate their radula. Radular morphologies of 2 specimens were imaged using a scanning electron microscopy (SEM). Radular sacs dissected from the body cavities and stored in pure ethanol, then treated with half-strength commercial bleach, leaving the clean radular teeth. Subsequently, the radula for SEM was rinsed in MilliQ water and dehydrated by increasing concentration of ethanol solution (20, 40, 60, 75, 100%). The dehydrated radula was dried completely using hexamethyldisilazane and then brought to the next step of SEM observation uncoated at 15 kV using a Hitachi TM-3000SEM. Radular characteristics of the *Alviniconcha* snail showed this snail has a broad radula with well-developed anterior supporting on the central tooth (Figure S15), indicating its ability of grazing on flat surface. The following linear measurements of 19 specimens were taken with digital vernier calipers: shell height (H), shell width (W), shell depth (D) and shell aperture height (AH), aperture width (AW). These measurements followed the methodology proposed by Chen *et al*., 2015. The following ratios were calculated from the values of each linear measurement: shell height/shell width (H/W); shell depth/shell width (D/W); shell aperture height/shell width (AH/W); shell aperture width/shell width (AW/W). The shell parameters of 19 specimens were shown in Table S4, and a scatter plot of shell width against shell height was shown in Figure S16. The measured five parameters of 19 *Alviniconcha* individuals indicated these snails might be at different growth stages, and they were linear across all life stages. Morphologically, all *Alviniconcha* populations are known to be extremely similar regardless of the species (Johnson et al., 2015), and this population was no exception, here molecular taxonomy can be used to provide reliable identification for these cryptic species.

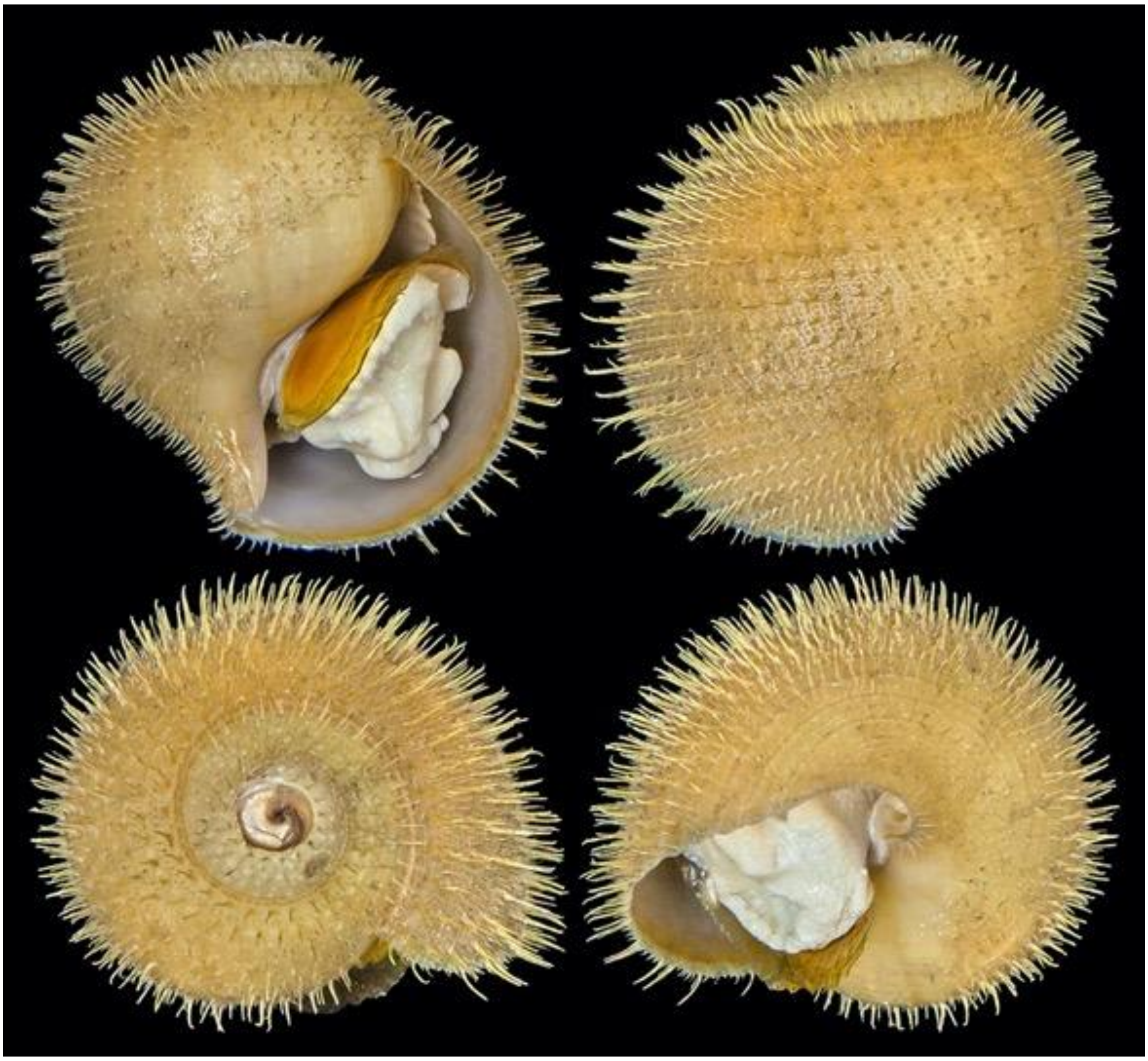

**Figure S14.** A photograph of snail *Alviniconcha marisindica* collected from the WHF which stored in absolute ethanol. Scale bar = 10 cm.

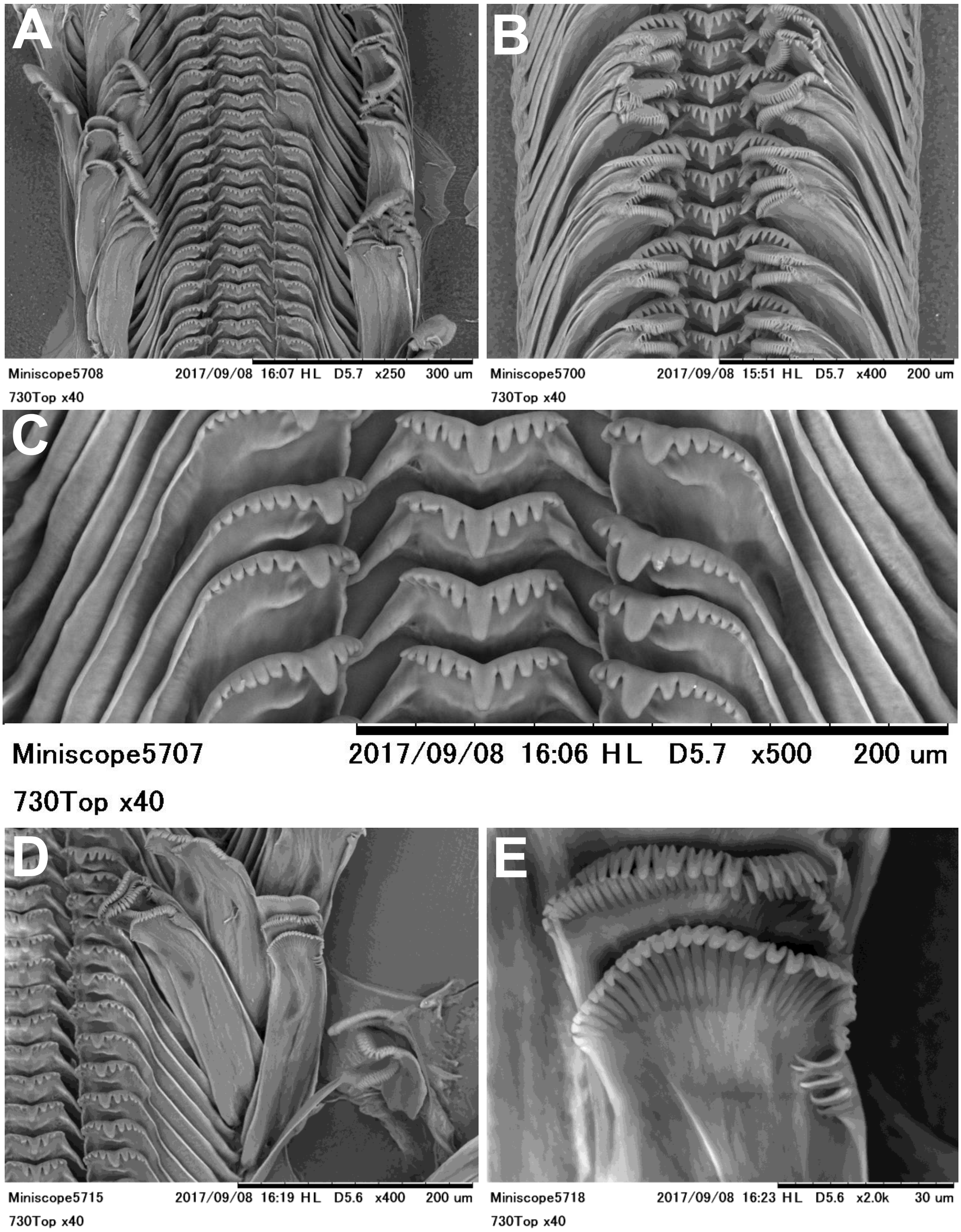

**Figure S15. SEM images of radula.** Overview: (**A**) *Alviniconcha marisindica* (individual 01); scale bar = 300 µm (**B**) *A. marisindica* (individual 01); scale bars = 200 µm. Central and lateral teeth close-up: (**C**) *A. marisindica* (individual 01); scale bars = 200 µm (**D**) *A. marisindica* (individual 02); scale bars = 200 µm. Marginal teeth close-up: (**E**) *A. marisindica* (individual 02); scale bars = 30 µm.

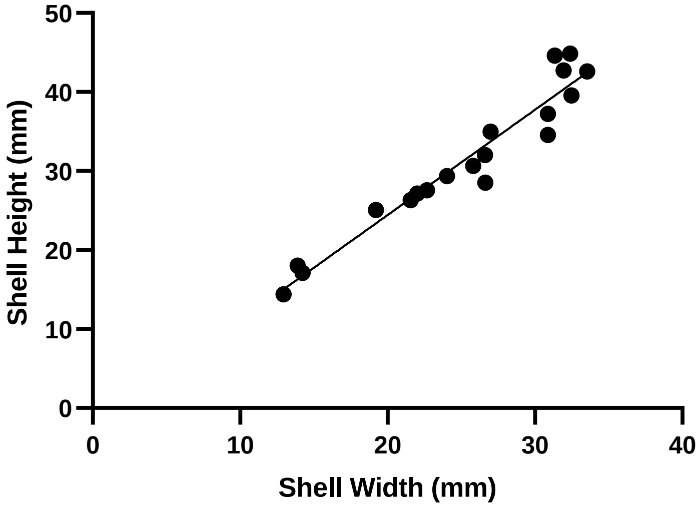

**Figure S16.** Scatter plot of shell width (diameter) vs. shell height across a size range of 19 specimens from *Alviniconcha* snails. (line of best fit formula: y = 1.3327x - 2.2346, $R^2$ = 0.93).

**Table S4.** Shell parameters of *Alviniconcha marisindica*. Range and proportion to shell width (diameter) are calculated from 19 specimens across a size range in this snail.

| Parameters (mm) | Shell | | | Aperture | |
|---|---|---|---|---|---|
| | **Width** | **Height** | **Depth** | **Width** | **Height** |
| **38I-DV129-1-$N_2$** | 14.23 | 17.12 | 13.27 | 9.48 | 14.66 |
| **38I-DV129-14** | 22.00 | 27.16 | 19.77 | 15.34 | 19.90 |
| **38I-DV129-15** | 21.57 | 26.33 | 20.69 | 15.57 | 22.65 |
| **38I-DV129-19-1** | 26.99 | 34.97 | 23.08 | 22.62 | 19.58 |
| **38I-DV129-19-2** | 31.35 | 44.62 | 31.40 | 25.69 | 31.33 |
| **38I-DV129-2-$N_2$** | 12.94 | 14.41 | 10.13 | 7.69 | 12.24 |
| **38I-DV129-20** | 25.80 | 30.65 | 21.57 | 16.85 | 25.31 |
| **38I-DV129-3-$N_2$** | 26.63 | 28.50 | 24.22 | 18.48 | 22.45 |
| **38I-DV129-4-$N_2$** | 31.94 | 42.73 | 31.42 | 21.75 | 27.39 |
| **38I-DV129-5-$N_2$** | 30.87 | 37.23 | 30.50 | 21.92 | 34.07 |
| **38I-DV129-5** | 32.48 | 39.57 | 30.45 | 23.49 | 31.41 |
| **38I-DV129-8** | 26.60 | 32.01 | 26.31 | 18.08 | 27.18 |
| **38I-DV129-9** | 24.04 | 29.34 | 21.71 | 21.99 | 18.75 |
| **38I-DV131-1** | 19.22 | 25.08 | 18.66 | 16.38 | 20.99 |
| **38I-DV131-3** | 22.67 | 27.55 | 21.19 | 15.73 | 22.77 |
| **38I-DV131-5** | 33.54 | 42.59 | 31.10 | 25.52 | 34.11 |
| **38I-DV131-6** | 30.87 | 34.55 | 26.53 | 22.58 | 29.10 |
| **38I-DV131-7** | 32.38 | 44.85 | 32.39 | 24.55 | 35.39 |
| **38I-DV131-9** | 13.90 | 18.01 | 12.99 | 10.43 | 15.00 |
| Range | 12.94–33.54 | 14.41–44.85 | 10.13–32.39 | 7.69–25.69 | 12.24–35.39 |
| Proportion of shell diameter (width) | 1 | 1.238 | 0.927 | 0.734 | 0.970 |
| SD of proportion | – | 0.089 | 0.060 | 0.078 | 0.107 |

**Molecular Taxonomy.** Fragments of the mitochondrial cytochrome c oxidase subunit I (COI) gene were amplified by polymerase chain reaction from 20 specimens, using the universal primers c (Folmer et al., 1994). Reaction volumes consisted of 1 µL template DNA, 17 µL deionized sterilized water, 20 µL 2× PCR MasterMix (Tiangen Biotech Co., Beijing, China), 1 µL of each primer (GenScript Biotech Co., Nanjing, China), for a total of 40 µL. PCR products were purified using TIANGEN Universal DNA Purification Kit (Tiangen Biotech Co., Beijing, China), and sequenced using Sanger sequencing platforms at the Beijing Genomics Institute (BGI)-Shenzhen. Bidirectional sequences were trimmed and assembled using DNASTAR EditSeq version 7.1.0. A phylogenetic tree was constructed based on *COI* gene sequences of *Alviniconcha* snails. Genetic distance estimated based on the *COI* sequence was about 3.4% between this new population and the CIR populations of *A. marisindica*, and a phylogenetic tree based on COI gene sequences of *Alviniconcha* snails shows our collected snail samples are closest to *A. marisindica* from the CIR (Figure S17A). Trimmomatic v0.39 (Bolger et al., 2014) and FastUniq (Xu et al., 2012) were used to trim the Illumina reads and remove duplicates. The clean reads were assembled using SPAdes v3.13.1 (Bankevich et al., 2012) with with k-mer sizes of 21, 33, 55, 77, 99, and 127 bp, and the products were pooled. Then, sequences of *12S*, *16S*, *18S*, *28S-D1*, *28S-D6*, *COI* and *H3* genes were picked out from the assembled contigs. A phylogenic tree constructed based on concatenated sequences of these seven genes indicated that this population was the closest to *A. marisindica* from the CIR (Figure S17B). The above results showed the *Alviniconcha* population from the CR was *Alviniconcha marisindica* which is the same species from the CIR.

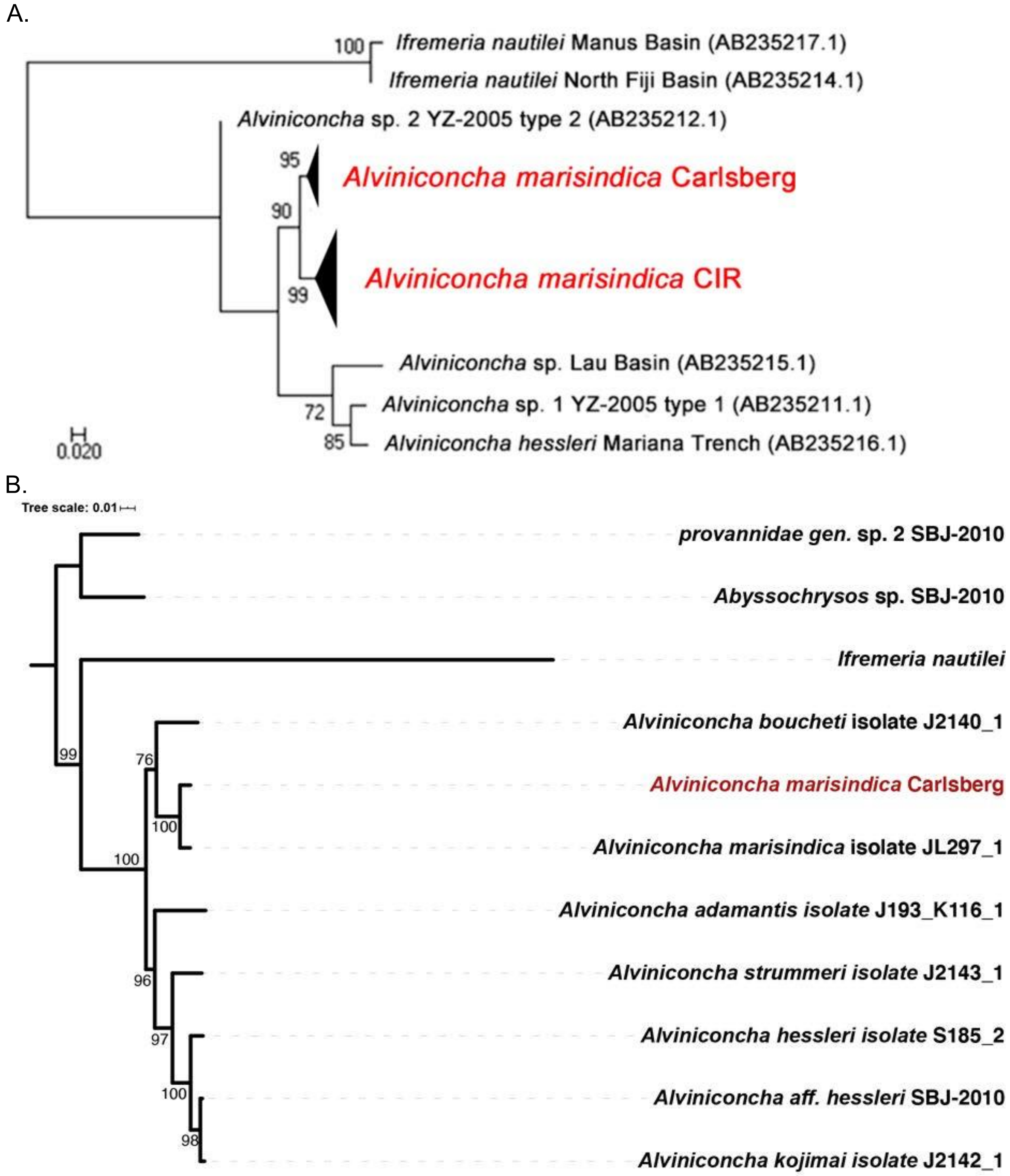

**Figure S17. Molecular taxonomy of the population of *Alviniconcha* snails from Wocan vent site on Carlsberg Ridge.** A phylogenetic tree of *Alviniconcha* snails based on (**A**) *COI* genes and (**B**) concatenated *12S*, *16S*, *18S*, *28S-D1*, *28S-D6*, *COI* and *H3* genes, the *Alviniconcha marisindica* population from Carlsberg Ridge (CR) and Central Indian Ridge (CIR) on Indian Ocean are in red colour. Numbers on the nodes indicate bootstrap support.

**Supplementary Note 2**

**Library Construction and Sequencing.** Genomic DNA was aliquoted and submitted to three sequencing platforms: Illumina, PacBio Sequel, and Oxford Nanopore Technologies (ONT). A library with a 350 bp insert size was constructed from gDNA following the standard protocol provided by Illumina (San Diego, CA, USA). After paired-end sequencing of the library at Novogene (Beijing, China), approximately 50 Gb of Illumina NovaSeq reads with a read length of 150 bp were generated. Illumina sequencing of total DNA from the gills and that of total DNA from the intestines were conducted similarly, with approximately 50 Gb of reads generated from each gill sample for endosymbiont genome assembly, approximately 6–8 Gb of reads generated from each of 20 gill filaments for symbiont genetic diversity analysis, and approximately 12 Gb of reads generated from each of three intestine specimens for metagenome analysis (Table S5).

For preparation of the single-molecule real-time (SMRT) DNA template for PacBio sequencing, the gDNA was sheared into large fragments (10 kb on average) using a Covaris® g-TUBE® device and then concentrated using AMPure® PB beads. DNA repair and purification were carried out according to the manufacturer's instructions (Pacific Biosciences). The blunt adapter ligation reaction was conducted on purified end-repair DNA, and after purification DNA sequencing polymerases became bound to SMRTbell templates. Finally, the library was quantified using a Qubit fluorometer v3.0. After sequencing with the PacBio Sequel System at the Hong Kong University of Science and Technology (HKUST) and Novogene, approximately 72 Gb of long reads were generated, with reads less than 4 kb in length discarded.

For ONT sequencing, a total of 3–5 μg of gDNA were used for the construction of each library following the '1D gDNA selecting for long reads (SQK-LSK109)' protocol from ONT. Briefly, gDNA was repaired and end-prepped as per standard protocol, before it was cleaned up with a 0.4× volume of AMPure XP beads. Adapter ligation and clean-up of the cleaned-repaired DNA were performed as per the standard protocol and the purified-ligated DNA was eluted using elution buffer. The DNA library was mixed with sequencing buffer and loading beads before it was loaded onto the SpotON sample port. Finally, sequencing was performed following the manufacturer's guidelines using the FLO-MIN106 R9.4 flow cell coupled to the MinIONTM platform (ONT, Oxford, UK). Raw reads were base-called according to the protocol in MinKNOW and written into fastq files, and 9.8 Gb of long reads were generated with reads less than 4Kb discarded. MinION sequencing of total DNA from one gill specimen was conducted using the same procedures, generating 3.5 Gb of reads for endosymbiont genome scaffolding. Illumina reads from gDNA were used for the genome survey of the Wocan *Alviniconcha marisindica*, and PacBio and ONT reads were used for the genome assembly (Table S6).

For eukaryotic transcriptome sequencing of different tissues, a 250–300 bp insert cDNA library of each tissue was constructed after removing the prokaryotic RNA and sequenced on the Illumina NovaSeq platform at Novogene to produce 150 bp paired-end reads. Since the RNA of gills includes the sequences from both the host and the symbiont, another 250–300 bp insert strand-specific library of each gill specimen was constructed using Ribo-Zero™ Magnetic Kit to sequence both eukaryotic and microbial RNA. Therefore, two sets of transcript sequencing data were produced for the gills, one for both the host and the symbiont, and the other for only the host. The meta-transcriptome sequencing of the intestine was conducted using the same methods. Approximately 5–10 Gb of reads were generated from each tissue (Table S5).

**Table S5.** The nucleic acid preparation and Illumina paired-end sequencing information of different tissues/organs from 16 *Alviniconcha marisindica* individuals in this study.

| *A. marisindica* Individuals | | Tissues/organs (Illumina sequencing reads) | | | | | | | | | | | | | |
|---|---|---|---|---|---|---|---|---|---|---|---|---|---|---|---|
| | | | | | | Gill | | | | | | | | | |
| Sample No. | Nucleic acid | Ct | DG | Go | Ft | GB | GD | GP | GA | Int | Man | ME | NF | Ne | VH |
| 38I-DV129-2-$N_2$ | RNA | 19,395,743 | 16,881,965 | 20,376,253 | 20,891,223 | 18,799,982<br>31,379,093 (meta) | 20,550,443 | – | – | – | 20,247,288 | 23,291,531 | 20,620,721 | 19,504,431 | – |
| | DNA | – | – | – | 260,164,976 | 132,877,148 (meta) | | | | – | – | – | – | – | – |
| 38I-DV129-3-$N_2$ | RNA | 26,279,803 | 22,597,795 | 26,184,458 | 31,126,320 | 24,055,311<br>28,067,797 (meta) | | | | – | 27,119,141 | – | 23,612,722 | – | – |
| | DNA | – | – | – | 193,071,397 | 208,807,531 (meta) | | | | – | – | – | – | – | – |
| 38I-DV129-16 | RNA | – | 19,369,344 | 20,374,319 | 20,238,317 | 19,805,564<br>27,801,163 (meta) | 21,251,951 | – | – | – | 22,464,985 | 20,884,683 | 27,397,183 | 20,404,036 | 21,587,150 |
| | DNA | – | – | – | – | 245,337,981 (meta) | | | | – | – | – | – | – | – |
| 38I-DV129-5-$N_2$ | RNA | – | – | – | – | 29,936,901 (meta) | | | | 38,477,522 (meta) | – | – | – | – | – |
| | DNA | – | – | – | – | – | | | | 45,356,320 (meta) | – | – | – | – | – |
| 38I-DV129-15 | RNA | – | – | – | – | 29,480,256 (meta) | | | | 30,937,560 (meta) | – | – | – | – | – |
| | DNA | – | – | – | – | – | | | | 43,407,952 (meta) | – | – | – | – | – |

| *A. marisindica* Individuals | | Tissues/organs (Illumina sequencing reads) | | | | | | | | | | | | | |
|---|---|---|---|---|---|---|---|---|---|---|---|---|---|---|---|
| | | Ct | DG | Go | Ft | Gill | | | | Int | Man | ME | NF | Ne | VH |
| **Sample No.** | **Nucleic acid** | | | | | **GB** | **GD** | **GP** | **GA** | | | | | | |
| 38I-DV131-6 | RNA | – | – | – | – | 27,747,516 (meta) | | | | 39,406,904 (meta) | – | – | – | – | – |
| | DNA | – | – | – | – | – | | | | 39,646,956 (meta) | – | – | – | – | – |
| 38I-DV129-14-1 | DNA | – | – | – | – | – | – | 22,194,485 (meta) | 26,009,046 (meta) | – | – | – | – | – | – |
| 38I-DV129-14-2 | DNA | – | – | – | – | – | – | 21,577,279 (meta) | 20,902,985 (meta) | – | – | – | – | – | – |
| 38I-DV129-19-1 | DNA | – | – | – | – | – | – | 24,755,940 (meta) | 30,199,263 (meta) | – | – | – | – | – | – |
| 38I-DV129-19-2 | DNA | – | – | – | – | – | – | 22,972,456 (meta) | 20,232,613 (meta) | – | – | – | – | – | – |
| 38I-DV129-20 | DNA | – | – | – | – | – | – | 24,269,919 (meta) | 26,623,681 (meta) | – | – | – | – | – | – |
| 38I-DV131-1 | DNA | – | – | – | – | – | – | 26,890,577 (meta) | 27,436,916 (meta) | – | – | – | – | – | – |
| 38I-DV131-3 | DNA | – | – | – | – | – | – | 26,916,916 (meta) | 25,686,865 (meta) | – | – | – | – | – | – |
| 38I-DV131-8 | DNA | – | – | – | – | – | – | 23,250,612 (meta) | 19,848,394 (meta) | – | – | – | – | – | – |

| *A. marisindica* Individuals | | Tissues/organs (Illumina sequencing reads) | | | | | | | | | | | | | |
|---|---|---|---|---|---|---|---|---|---|---|---|---|---|---|---|
| | | Ct | DG | Go | Ft | Gill | | | | Int | Man | ME | NF | Ne | VH |
| **Sample No.** | **Nucleic acid** | | | | | **GB** | **GD** | **GP** | **GA** | | | | | | |
| 38I-DV131-9-1 | DNA | – | – | – | – | – | – | 25,868,238 (meta) | 20,261,863 (meta) | – | – | – | – | – | – |
| 38I-DV131-9-2 | DNA | – | – | – | – | – | – | 27,342,826 (meta) | 19,909,893 (meta) | – | – | – | – | – | – |

**Table S6.** The PacBio and ONT sequencing information of DNA from one *Alviniconcha marisindica* individual for holobiont genome assembly.

| 38I-DV129-2-$N_2$ | Tissues/organs (Long-read sequencing reads) | |
|---|---|---|
| | **Foot/Neck** | **Gill** |
| **PacBio** | 11,232,982 | – |
| **ONT** | 2,758,946 | 833,727 |

**Supplementary Note 3**

***de novo* Hybrid Assembly of the Host Genome.** Trimmomatic v0.39 (Bolger et al., 2014) was used to trim the Illumina adapters and low-quality bases (base quality ≤ 20). The genome size was estimated by a 17-mer histogram which was shown in Figure S18. The kmer histogram was generated by Platanus v1.2.4 (Kajitani et al., 2014) with settings of -k 17 -s 10 -u 0.2 -t 20 -m 320. The genome heterozygosity was evaluated as 0.88% using GenomeScope (Vurture et al., 2017). Several genome assembly pipelines were applied to assemble the genome with PacBio and ONT reads, including PacBio-only approaches (e.g. minimp2+miniasm (Li, 2016) and wtdbg2 (Ruan and Li, 2019) and PacBio-ONT hybrid approaches (e.g. MaSuRCA version 3.2.8 (Zimin et al., 2013), FMLRC (Wang et al., 2018) + smartdenovo (Ruan, 2018) and FMLRC (Wang et al., 2018) + wtbg2 (Ruan and Li, 2019)). The detailed settings of each assembly pipeline are shown below.

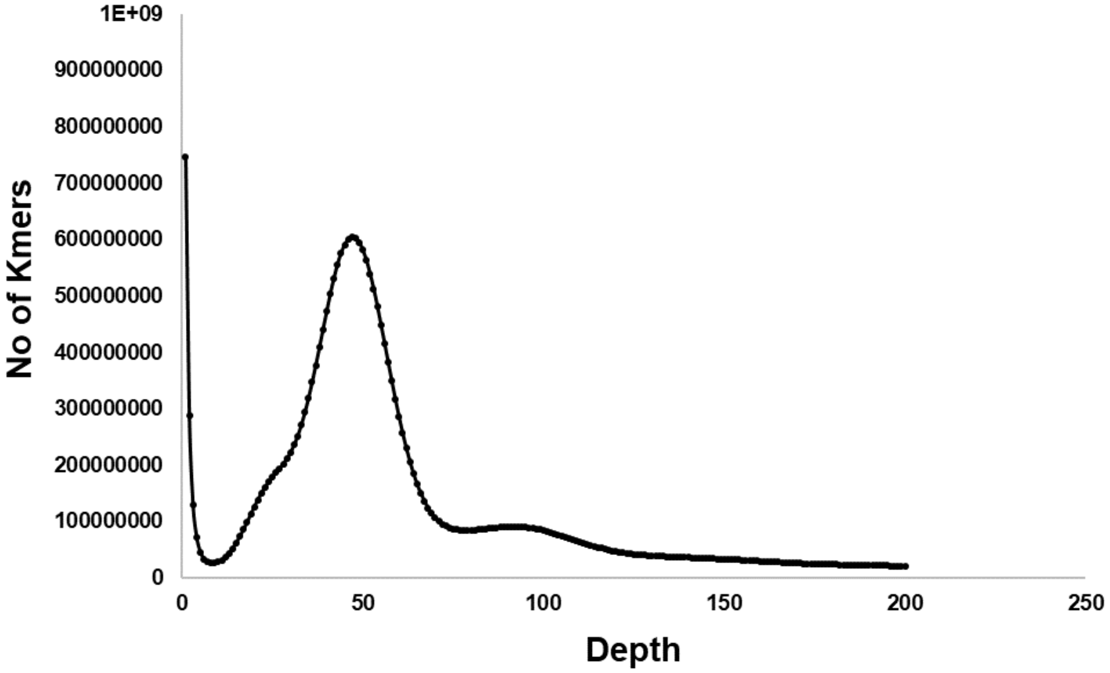

**Figure S18.** A 17-mer histogram for the *Alviniconcha marisindica* genome.

**Genome Assembly.**

1) minimap2 + miniasm on all of the Pacbio reads (Li, 2016).

```
minimap2 -X -t 64 -x ava-pb AlviPacBio_4K.fasta AlviPacBio_4K.fasta > reads.paf
miniasm -f AlviPacBio_4K.fasta reads.paf > Amar_default.gfa
awk '/^S/{print ">"$2"\n"$3}' Amar_default.gfa | fold > Amar_default.fa
```

2) wtdbg2 pipeline on all of the Pacbio reads (Ruan and Li, 2019).

```
wtdbg2 -t 12 -i AlviPacBio_4K.fasta -fo Amar_Pb -x sq -L 5000 -g 0.8g
wtpoa-cns -t 12 -i Amar_Pb.ctg.lay.gz -fo Amar_Pb.ctg.lay.fa
```

3) MaSuRCA v3.2.8 pipeline (Zimin et al., 2013).
The ONT reads were concatenated with the PacBio reads serve as the input of 'NANOPORE=' in the configure file, as suggested by the developer. The rest is as default.

4) FMLRC (Wang et al., 2018) hybrid correction and further assembled by third party assembler. The command was listed as follows:

```
gunzip -c All_trim.fq.gz | awk 'NR % 4 == 2' | sort | tr NT TN | ropebwt2 -LR | tr NT TN | fmlrc-convert Amar_comp_msbwt.npy
awk 'NR % 4 == 2' Illumina_trim.fq | sort | tr NT TN | ropebwt2 -LR | tr NT TN | fmlrc-convert Amar_comp_msbwt.npy
fmlrc -p 12 Amar_comp_msbwt.npy Amar_Pb_ONT.fa Amar_Pb_ONT_fmlrc_EC.fasta
```

The corrected long reads were either assembled by smartdenovo (https://github.com/ruanjue/smartdenovo) or wtdbg2 (Ruan and Li, 2019).

A comparison of the assembly statistics of different pipelines (Table S7) showed that the FMLRC+wtdbg2 assembly was the best and therefore this assembly was used in the downstream analyses. The assembly was carried out as follows: the ONT reads were concatenated with PacBio reads and error corrected with Illumina reads using FMLRC (Wang et al., 2018). This hybrid error correction method was selected based on previous benchmarking analysis on the available tools using Illumina reads for correction of PacBio/ONT long reads (Fu et al., 2019). The corrected long reads were then assembled using wtdbg2 using the setting "-x preset2" (Ruan and Li, 2019).

**Table S7.** The statistics of the genome assembly. The number in bold indicates the best among all of the assemblers.

| | **minimap2+ miniasm*** | **wtdbg2*** | **MaSuRCA** | **FMLRC+ smartdenovo** | **FMLRC+ wtdbg2** |
|---|---|---|---|---|---|
| **Total size** | 934.7Mb | **824.2Mb** | 917.9Mb | 846.3Mb | 853.2Mb |
| **Number of contig** | 15,233 | 6526 | 30,250 | **5220** | 6704 |
| **N50** | 106.8Kb | 346.2Kb | 71.5Kb | 320.6Kb | **708.1Kb** |
| **NG50** | 122.9Kb | 343.7Kb | 104.1Kb | 325.9Kb | **728.0Kb** |
| **Longest contig** | 0.89Mb | 2.38Mb | 0.83Mb | 2.68Mb | **4.75Mb** |
| **Mean length** | 61.4Kb | 126.3Kb | 37.8Kb | **162.1Kb** | 127.3Kb |
| **No. of contig over 1Mb** | 0 | 56 | 0 | 59 | **204** |

* using PacBio sequences only.

**Bacterial Contamination Removal.** MetaBAT 2 (Kang et al., 2015) and MaxBin 2.0 (Wu et al., 2016) were used to perform genome binning of the assembled contigs for checking microbial contamination, the resulting output microbial genomes were checked by CheckM v1.1.2 (Parks et al., 2015), based on the presence of particular marker genes. Open reading frames (ORFs) of the binned microbial genome were predicted using Prodigal v2.6.3 (Hyatt et al., 2010), then BLASTp was used to align all predicted protein sequences from the binned genome to the NCBI NR protein database using an e-value 1e-5 with 20 best hits, and the taxonomic assignment of each protein was imported to MEGAN v5.7.0 (Huson et al., 2011) using the lowest common ancestor (LCA) method. These sequences were confirmed as contamination and removed from the downstream analysis (Table S8).

**Table S8.** Information of the bacterial sequences removed from the assembled host contigs.

| Marker lineage | Total contig clusters (Mb) | Total ORFs | GC content (%) | Completeness (%) | Contamination (%) |
|---|---|---|---|---|---|
| k__Bacteria (UID3060) | 9.44 | 9,452 | 41.00 | 83.65 | 0 |
| c__Mollicutes (UID2395) | | | | 31.56 | 3.26 |

**Supplementary Note 4**

**Annotation of the Host Genome.** The proportion of repeat content in the *Alviniconcha* genome was evaluated as 20.25%, the classified repeat content in the genome was shown in Table S9.

**Table S9.** The classified repeat content in the genome.

| Class | Count | bpMasked | %masked |
|---|---|---|---|
| DNA transposons | 130115 | 6754060 | 0.82% |
| Academ | 759 | 40214 | 0.00% |
| Academ2 | 3 | 454 | 0.00% |
| CMC-Chapaev | 541 | 29410 | 0.00% |
| CMC-Chapaev-3 | 106 | 4290 | 0.00% |
| CMC-EnSpm | 60985 | 4146368 | 0.50% |
| CMC-Transib | 923 | 48612 | 0.01% |
| Crypton | 546 | 32220 | 0.00% |
| Crypton-H | 39 | 23800 | 0.00% |
| Crypton-V | 754 | 33472 | 0.00% |
| Dada | 3988 | 218202 | 0.03% |
| Ginger | 30422 | 2183414 | 0.27% |
| Harbinger | 4 | 253 | 0.00% |
| IS3EU | 3370 | 170165 | 0.02% |
| Kolobok | 68 | 3191 | 0.00% |
| Kolobok-Hydra | 18710 | 1454083 | 0.18% |
| Kolobok-T2 | 2497 | 131087 | 0.02% |
| MULE-F | 1 | 105 | 0.00% |
| MULE-MuDR | 3833 | 244134 | 0.03% |
| MULE-NOF | 22 | 6202 | 0.00% |
| Maverick | 21062 | 1112659 | 0.14% |
| Merlin | 1196 | 52982 | 0.01% |
| Novosib | 41391 | 5173020 | 0.63% |
| P | 751 | 34659 | 0.00% |
| PIF-Harbinger | 8296 | 532246 | 0.06% |
| PIF-ISL2EU | 2246 | 105459 | 0.01% |
| PiggyBac | 816 | 41911 | 0.01% |
| Sola | 60298 | 6225659 | 0.76% |

| | | | | |
|---|---|---|---|---|
| | TcMar | 951 | 44699 | 0.01% |
| | TcMar-Ant1 | 4 | 198 | 0.00% |
| | TcMar-Fot1 | 980 | 51365 | 0.01% |
| | TcMar-ISRm11 | 54 | 2772 | 0.00% |
| | TcMar-Mariner | 36 | 4471 | 0.00% |
| | TcMar-Pogo | 2 | 113 | 0.00% |
| | TcMar-Sagan | 1 | 94 | 0.00% |
| | TcMar-Stowaway | 60 | 3541 | 0.00% |
| | TcMar-Tc1 | 1807 | 90403 | 0.01% |
| | TcMar-Tc2 | 93 | 11042 | 0.00% |
| | TcMar-Tc4 | 7 | 456 | 0.00% |
| | TcMar-Tigger | 62 | 28309 | 0.00% |
| | TcMar-m44 | 2 | 144 | 0.00% |
| | Zator | 4 | 286 | 0.00% |
| | Zisupton | 856 | 55699 | 0.01% |
| | hAT | 37068 | 2087935 | 0.25% |
| | hAT-Ac | 26183 | 1603319 | 0.20% |
| | hAT-Blackjack | 404 | 18786 | 0.00% |
| | hAT-Charlie | 8142 | 661251 | 0.08% |
| | hAT-Pegasus | 2596 | 134949 | 0.02% |
| | hAT-Tag1 | 2889 | 151116 | 0.02% |
| | hAT-Tip100 | 4325 | 219024 | 0.03% |
| | hAT-Tol2 | 10 | 324 | 0.00% |
| | hAT-hAT5 | 75 | 15348 | 0.00% |
| | hAT-hATm | 162 | 8305 | 0.00% |
| | hAT-hATw | 81 | 18227 | 0.00% |
| | hAT-hATx | 1 | 125 | 0.00% |
| | hAT-hobo | 3 | 210 | 0.00% |
| LINE | | 2857 | 233083 | 0.03% |
| | Ambal | 165 | 11493 | 0.00% |
| | CR1 | 854 | 114328 | 0.01% |
| | CR1-Zenon | 144 | 36221 | 0.00% |
| | CRE | 5 | 498 | 0.00% |
| | DRE | 68 | 4503 | 0.00% |
| | Dong-R4 | 8 | 350 | 0.00% |
| | I | 478 | 80933 | 0.01% |
| | Jockey | 2741 | 346028 | 0.04% |
| | L1 | 7507 | 541190 | 0.07% |
| | L1-Tx1 | 6572 | 412478 | 0.05% |
| | L2 | 8569 | 861778 | 0.10% |
| | LOA | 6 | 1229 | 0.00% |
| | Penelope | 6485 | 384678 | 0.05% |
| | Proto1 | 23 | 1187 | 0.00% |
| | R1 | 3362 | 372621 | 0.05% |
| | R2 | 244 | 25636 | 0.00% |
| | RTE | 5 | 428 | 0.00% |
| | RTE-BovB | 395 | 91147 | 0.01% |
| | RTE-RTE | 1 | 48 | 0.00% |

| | | | | |
|---|---|---|---|---|
| | RTE-X | 675 | 104048 | 0.01% |
| | Rex-Babar | 2063 | 164985 | 0.02% |
| | Tad1 | 33 | 4512 | 0.00% |
| LTR | | 4210 | 235229 | 0.03% |
| | Caulimovirus | 1 | 39 | 0.00% |
| | Copia | 6077 | 331270 | 0.04% |
| | DIRS | 552 | 40187 | 0.00% |
| | ERV | 8929 | 598448 | 0.07% |
| | ERV-Foamy | 1 | 164 | 0.00% |
| | ERV1 | 26664 | 1251321 | 0.15% |
| | ERV4 | 45 | 2232 | 0.00% |
| | ERVK | 14030 | 620405 | 0.08% |
| | ERVL | 138 | 8245 | 0.00% |
| | ERVL-MaLR | 1 | 64 | 0.00% |
| | Gypsy | 31901 | 3007753 | 0.37% |
| | Ngaro | 3173 | 245231 | 0.03% |
| | Pao | 819 | 88223 | 0.01% |
| | Viper | 17 | 1082 | 0.00% |
| Other | | 9 | 3237 | 0.00% |
| | DNA_virus | 11 | 541 | 0.00% |
| RC | | -- | -- | -- |
| | Helitron | 19982 | 1423511 | 0.17% |
| **Retroposon** | | 1 | 111 | 0.00% |
| | SVA | 5 | 1767 | 0.00% |
| SINE | | 54 | 6752 | 0.00% |
| | 5S | 3 | 233 | 0.00% |
| | 5S-Deu-L2 | 57 | 4163 | 0.00% |
| | 5S-Sauria-RTE | 2 | 153 | 0.00% |
| | 7SL | 3 | 688 | 0.00% |
| | Alu | 1 | 46 | 0.00% |
| | B4 | 117 | 5651 | 0.00% |
| | ID | 16 | 778 | 0.00% |
| | MIR | 128 | 7151 | 0.00% |
| | RTE-BovB | 3 | 524 | 0.00% |
| | U | 8 | 466 | 0.00% |
| | tRNA | 5209 | 239468 | 0.03% |
| | tRNA-7SL | 4 | 207 | 0.00% |
| | tRNA-C | 31 | 1306 | 0.00% |
| | tRNA-CR1 | 11 | 489 | 0.00% |
| | tRNA-Core | 8 | 354 | 0.00% |
| | tRNA-Deu | 2 | 187 | 0.00% |
| | tRNA-Deu-L2 | 379 | 21699 | 0.00% |
| | tRNA-L2 | 833 | 45173 | 0.01% |
| | tRNA-Mermaid | 385 | 19608 | 0.00% |
| | tRNA-RTE | 5 | 173 | 0.00% |
| | tRNA-Sauria-RTE | 1 | 0 | 0.00% |
| | tRNA-V | 5 | 188 | 0.00% |
| Segmental | | 1 | 54 | 0.00% |

| | | | | |
|---|---|---|---|---|
| Unknown | | 25302 | 1390838 | 0.17% |
| | centromeric | 3 | 364 | 0.00% |
| **total** | interspersed | 672997 | 47414017 | 5.77% |
| **Low_complexity** | | 120231 | 7742961 | 0.94% |
| **RNA** | | 147 | 13262 | 0.00% |
| **Satellite** | | 46972 | 5890412 | 0.72% |
| | 5S | 112 | 13331 | 0.00% |
| | W-chromosome | 1 | 155 | 0.00% |
| | centr | 7 | 1762 | 0.00% |
| | macro | 12 | 821 | 0.00% |
| | telo | 3 | 1409 | 0.00% |
| Simple_repeat | | 1277876 | 105294431 | 12.81% |
| rRNA | | 65 | 25179 | 0.00% |
| scRNA | | 2 | 74 | 0.00% |
| snRNA | | 59 | 7422 | 0.00% |
| tRNA | | 503 | 29244 | 0.00% |
| Total | | **2118987** | **166434480** | **20.25%** |

**Host Gene Family Identification and Phylogenomic Analysis.** A total of 26 lophotrochozoan genomes were analysed for clues to the gene family evolution (Table S10). Time frame constraints imposed to calibrate the topology tree generated from RAxML (Figure S19) are shown in Table S11.

**Table S10.** Lophotrochozoan genomes that were used for the phylogeny and gene family analyses.

| **Phylum** | **Species** | **Reference** |
|---|---|---|
| **Annelida** | *Capitella teleta* | Simakov, Marletaz *et al.*, 2013 (88) |
| **Nemertea** | *Notospermus geniculatus* | Luo, Kanda *et al.*, 2018 (89) |
| **Phoronida** | *Phoronis australis* | Luo, Kanda *et al.*, 2018 (89) |
| **Brachiopoda** | *Lingula anatina* | Luo, Kanda *et al.*, 2018 (89) |
| **Mollusca** | *Euprymna scolopes* | Belcaid, Casaburi *et al.*, 2019 (90) |
| | *Octopus bimaculoides* | Albertin, Simakov *et al.*, 2015 (91) |
| | *Sinonovacula constricta* | Dong, Zeng *et al.*, 2019 (92) |
| | *Ruditapes philippinarum* | Yan, Nie *et al.*, 2019 (93) |
| | *Mizuhopecten yessoensis* | Wang, Zhang *et al.*, 2017 (94) |
| | *Bathymodiolus platifrons* | Sun, Zhang *et al.*, 2017 (95) |
| | *Modiolus philippinarum* | Sun, Zhang *et al.*, 2017 (95) |
| | *Pinctada fucata* | Takeuchi, Koyanagi *et al.*, 2016 (96) |
| | *Crassostrea gigas* | Zhang, Fang *et al.*, 2012 (97) |
| | *Haliotis rufescens* | Masonbrink, Purcell *et al.*, 2019 (98) |
| | *Haliotis rubra* | Gan, Tan *et al.*, 2019 (99) |
| | *Chrysomallon squamiferum* | Sun *et al.*, 2020 (100) |
| | *Lottia gigantea* | Simakov, Marletaz *et al.*, 2013 (88) |
| | *Alviniconcha marisindica* | This study |
| | *Lanistes nyassanus* | Sun, Mu *et al.*, 2019 (101) |
| | *Marisa cornuarietis* | Sun, Mu *et al.*, 2019 (101) |
| | *Pomacea canaliculata* | Sun, Mu *et al.*, 2019 (101) |
| | *Aplysia californica* | NCBI AplCal3.0 |
| | *Achatina fulica* | Guo, Zhang *et al.*, 2019 (102) |

| | | |
|---|---|---|
| | *Biomphalaria glabrata* | Adema, Hillier *et al.*, 2017 (103) |
| | *Radix auricularia* | Schell, Feldmeyer *et al.*, 2017 (104) |
| | *Elysia chlorotica* | Cai, Li *et al.*, 2019 (105) |

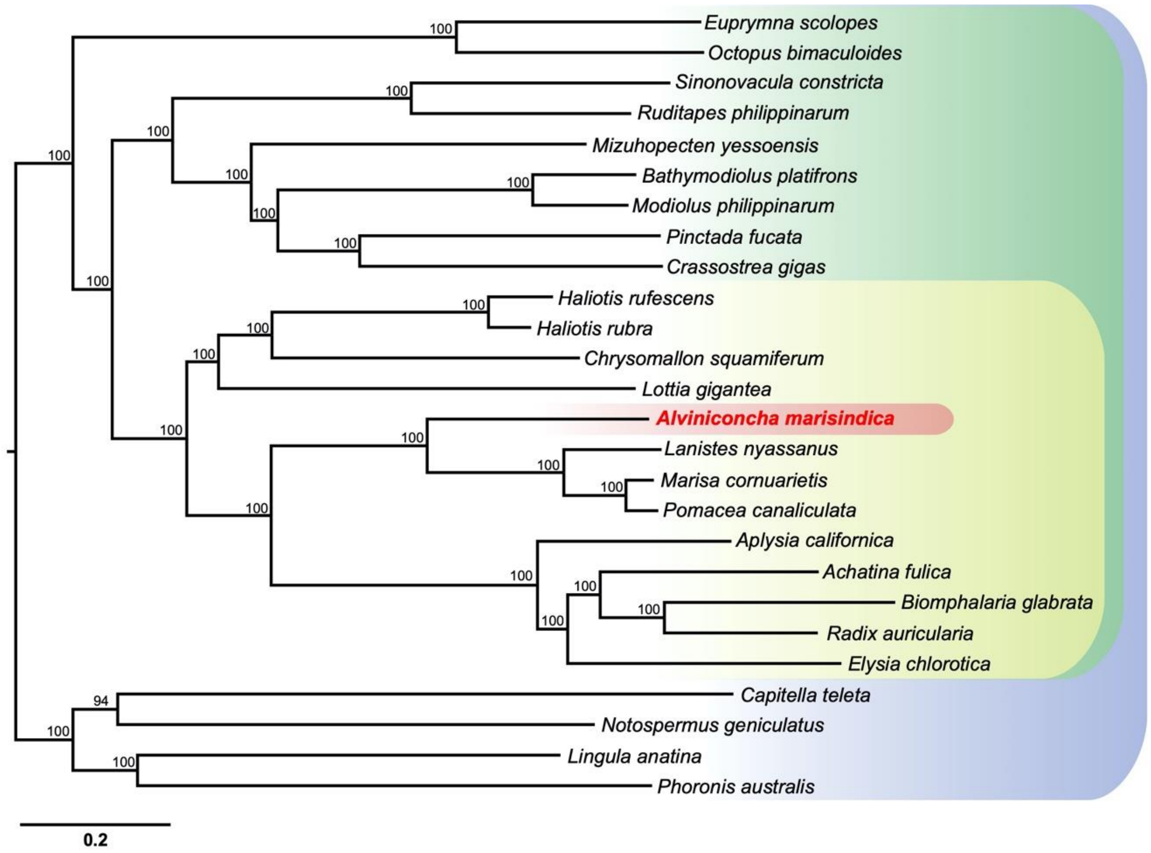

**Figure S19.** Genome-based phylogeny of selected taxa showing the position of the *Alviniconcha marisindica* among lophotrochozoans.

**Table S11.** The time constrain that was applied to calibrate the species divergent time in the MCMCTree analysis.

| Calibration node | Date Range | Reference |
|---|---|---|
| **The first appearance of Abyssochrysidae** | hard minimum bound = 168 Ma | Kaim and Conti, 2010 (106) |
| ***L. nyassanus and P. canaliculata*** | hard max time-point of 150 Ma | Hayes, Cowie *et al.*, 2009; Sun, Mu *et al.*, 2019 (101, 107) |
| ***A. californica* and *B. glabrata*** | minimum = 168.6 Ma and soft maximum = 473.4 Ma | Benton, Donoghue *et al.*, 2009 (108) |
| **The first appearance of both the Stylommatophora and Hygrophila** | hard minimum bound = 130 Ma | Tillier, 1996; Wade, Mordan *et al.*, 2001 (109, 110) |
| **Caenogastropoda and Heterobranchia** | hard minimum bound = 390 Ma | Jörger, Stöger *et al.*, 2010 (111) |
| ***A. californica* (or *B. glabrata*) and *L. gigantea*** | minimum = 470.2 Ma and soft maximum = 531.5 Ma | Benton, Donoghue *et al.*, 2015 (112) |
| **The first appearance of molluscs** | minimum = 532 Ma and soft maximum = 549 Ma | Benton, Donoghue *et al.*, 2015 (112) |
| **The first appearance of Pteriomorpha** | hard minimum = 465.0 Ma | Stoger, Sigwart *et al.*, 2013 (113) |

| | | |
|---|---|---|
| **The first appearance of Lophotrochozoan** | minimum = 550.25 Ma and soft maximum = 636.1Ma | Benton, Donoghue *et al.*, 2015 (112) |

**Supplementary Note 5**

**Microbial Community Composition of Gill Symbionts.** To determine the microbial ribotype and community composition in the gill of our specimen, approximately 1,500-bp 16S rRNA gene fragment was amplified by PCR from the gill using universal bacterial primers 8f and 1492r (Paster et al., 1998). A clone library of 16S rRNA gene was constructed, and 50 clones were randomly picked from the library and sequenced. The resultant sequences were analyzed using RDP Naive Bayesian rRNA Classifier v2.11 (Wang et al., 2007) with an 80% confidence threshold to reveal bacterial species composition in the gill, the result indicated a single bacterial ribotype in the gill of *A. marisindica*. To understand the phylogenetic position of the *Alviniconcha* endosymbiont, a phylogenetic tree was constructed based on 16S rRNA gene sequences, 16S rRNA gene sequence of our specimen was obtained from the above clone library result and 16S rRNA gene sequences of other marine bacteria including endosymbionts from other *Alviniconcha* snail species were obtained from GenBank. The sequences were aligned using MUSCLE and trimmed using TrimAL v1.4 (Capella-Gutiérrez et al., 2009). The Maximum Likelihood (ML) method was used to construct the phylogenetic tree using RaxML v8.2.11 (Stamatakis et al., 2005) under the GTR + CAT model with 1000 bootstrap replicates. The endosymbiont of *A. marisindica* from the CR in this study was clustered with the endosymbiont of *A. marisindica* from the CIR and free-living Campylobacterota from deep-sea vents (Figure S20).

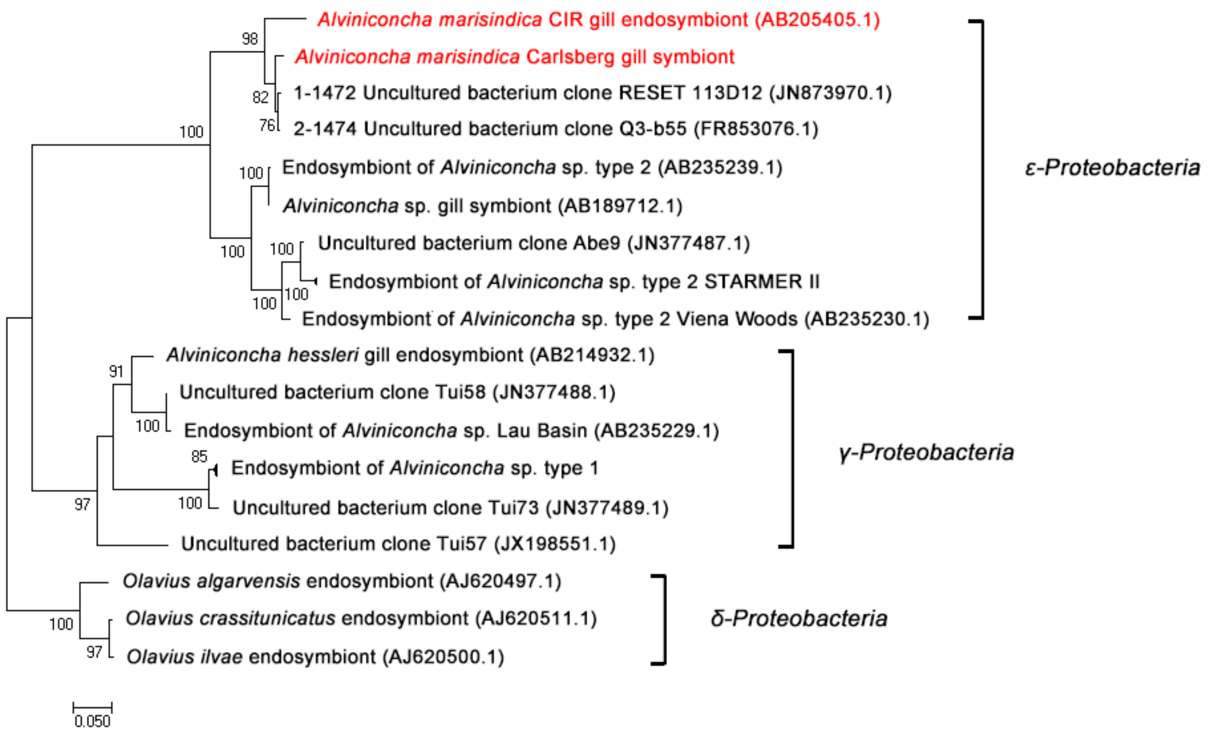

**Figure S20. The maximum likelihood (ML) phylogenetic tree of endosymbionts of *Alviniconcha* snails and other marine bacteria based on 16S rRNA gene.** Numbers above branches represent ML bootstrap values based on 1,000 iterations, with 100 as the highest value. *δ-Proteobacteria* symbionts of genus *Olavius* are taken as outgroup, the endosymbiont of *Alviniconcha marisindica* from the CR (this study) and the CIR are in red color.

**Genome Binning, Annotation, and Functional Analysis.** Contigs potentially belonging to the campylobacterotal endosymbiont genome were separated from its host genome using three binning methods. The first method was modified from Albertsen *et al*., 2013 and the modified binning process was followed as our previous study (Yang et al., 2020). First, the clean Illumina reads were mapped to the assembled contigs using Bowtie2 v2.3.4.3 (Langmead and Salzberg, 2012), and the coverage of each contig was calculated using SAMTOOLS v1.9 (Li et al., 2009). Prodigal v2.6.3 (Hyatt et al., 2010) was used to predict open reading frames (ORFs) and protein functional domains were predicted using HMMER 3.1b2 (Eddy et al., 2009) under the 100 + HMM model. Taxonomic affiliation of all HMM positive ORFs were determined using BLASTp (Altschul et a., 1990) against NCBI nonredundant (NR) protein database, and the taxonomic assignment of each protein was imported to MEGAN v5.7.0 (Huson et al., 2011) using the lowest common ancestor (LCA) method with the parameters of Min Score 50, Max Expected 0.01, Top Percent 5 and LCA Percent 100. The results were subsequently analyzed in RStudio (https://www.rstudio.com/) with the libraries of vegan, plyr, RColorBrewer and alphahull. Sequences representing the draft symbiont genome were then extracted from the assembled contigs of both the host and the symbiont, based on the combination of sequencing coverage, GC content and taxonomic classification (Figure S3A).

Coding sequences (CDS) in the genomes of *Alviniconcha* symbionts were predicted and translated using Prodigal v2.6.3 (Hyatt et al., 2010). Gene functions were determined by using BLASTp to align the candidate sequences to the NCBI non-redundant (NR) and SwissProt protein databases with the settings of "-evalue 1e-5 -word_size 3 -num_alignments 20 -max_hsps 20". Blast2GO® (Götz et al., 2008) employed with EggNOG mapper (Huerta-Cepas et al., 2017) was applied to assign Gene Ontology (GO) terms and Cluster of Orthologous Groups (COG) to the protein sequences via GO and EggNOG databases. Kyoto Encyclopedia of Genes and Genomes (KEGG) Automatic Annotation Server (KAAS) (Kanehisa and Goto, 2000) was used to conduct the KEGG pathway annotation analysis with the bi-directional best hit (BBH) method. The RAST Server (Overbeek et al., 2014) was used to annotate the genome based on SEED database and build metabolic model from KEGG annotation.

In addition to Campylobacterotal symbiont which has been confirmed as the dominant endosymbiont in the gill from the above results of clone library, another Mollicutes symbiont was found coexist in the metagenome binning (Figure S3A) which showed a much lower abundance than the Campylobacterotal symbiont. The general genomic features of these two symbionts were shown in Table S12. The genomic overview of Mollicutes was shown in Figure S21. Large coding sequences in the genome of Mollicutes were annotated with hypothetical protein, only 448 protein sequences were assigned functions based on the GO, COG, KEGG and SEED databases. Metabolic pathways that sustain life activities such as glycolysis/gluconeogenesis, DNA replication and repair were found in the Mollicutes, however, many genes involved in the pathways of carbon fixation, amino acids biosynthesis and chemoautotrophs (e.g. sulfur oxidation, methane oxidation, etc.) were missing, indicating the Mollicutes in the gill of *A. marisindica* cannot offer the host benefits for surviving in deep-sea vents. This is contrast with mutualism, in which both the Campylobacterotal endosymbiont and *A. marisindica* snail benefit from each other, in this case, we concluded the Mollicutes have commensal relationship with the *Alviniconcha marisindica*.

**Table S12.** The general genomic features of *A. marisindica* symbionts.

| Symbiont | Campylobacterota | Mollicutes |
|---|---|---|
| **Genome size (Mb)** | 1.47 | 0.79 |
| **No. of scaffolds** | 2 | 15 |
| **$N_{50}$ (Kb)** | 1458.35 | 175.95 |
| **GC%** | 37.09 | 24.78 |
| **Completeness%** | 98.16 | 82.33 |
| **Contamination%** | 0.82 | 1.88 |
| **No. of Genes** | 1,429 | 1,358 |
| **No. of CDS** | 1,386 | 1,332 |
| **No. of functions assigned** | 1,324 | 448 |

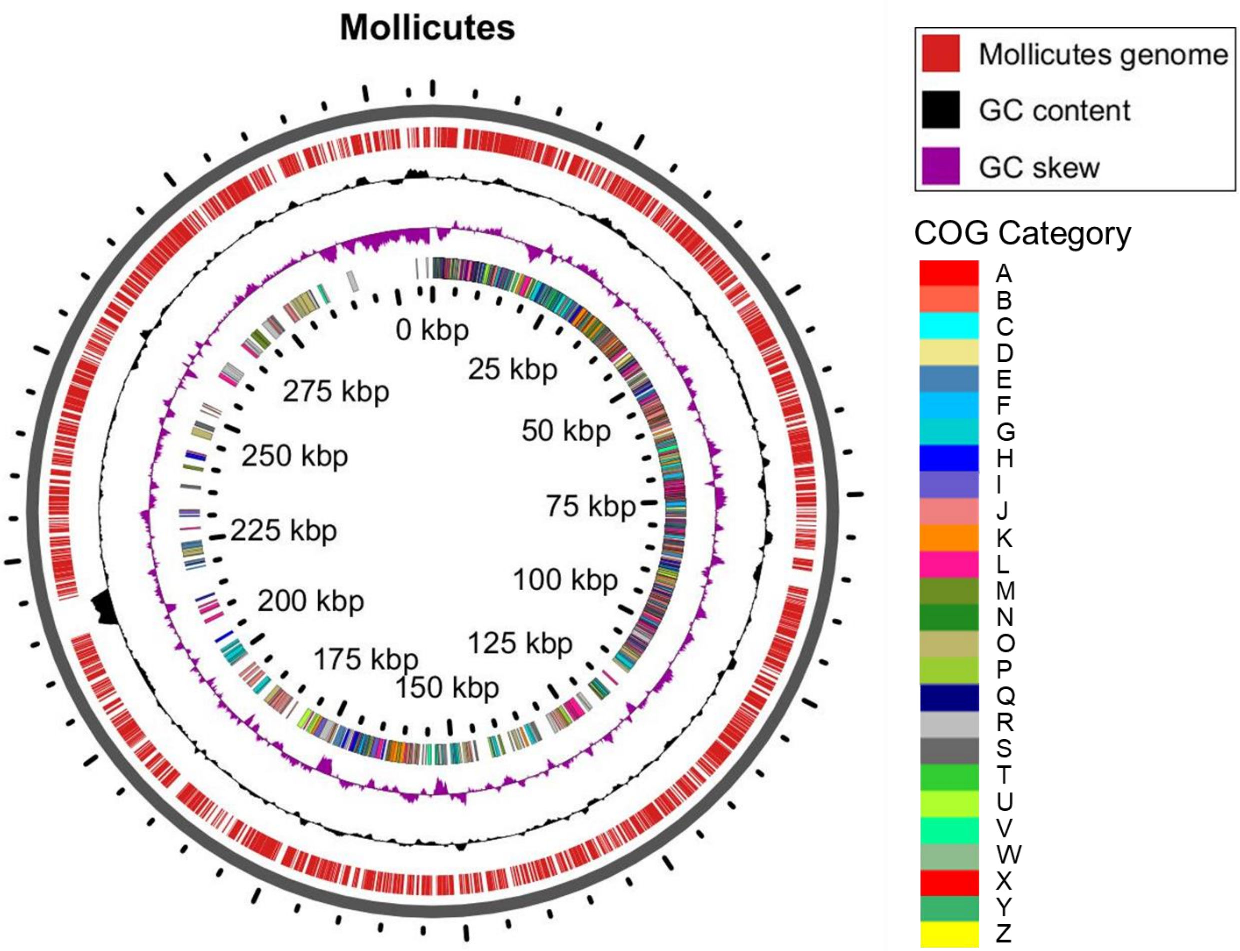

**Figure S21. The overview of genome of Mollicutes symbiont in the gill *of A. marisindica* constructed using GView Server.** The innermost ring represents the COG categories of the symbiont. The second and third rings show GC skew and GC content. The outermost ring represents the symbiont genome sequence. The identifiers of outside-to-inside rings are listed on the right.

**Supplementary Note 6**

**Gut microbial community and relative abundance.** The bacterial abundance of metagenomic sequences was figured out using Kaiju (Menzel et al., 2016) based on subset of NCBI BLAST *nr* database containing all proteins belonging to Archaea, Bacteria and Viruses. In the metagenome of gill, the bacterial sequences account for 12.08–21.44% (Figure S6A) which is much higher than the intestinal bacteria (2.43–5.41%, Figure S6B). After the metagenome assembly of the intestinal contents, bacterial taxonomic classification of the assembled contigs was figured out using Kaiju (Menzel et al., 2016), the microbial composition includes 38.74–49.30% of Proteobacteria, 7.44–11.15% of Actinobacteria, 8.98–10.79% of Firmicutes, 1.89–9.75% of Tenericutes and 3.20–4.42% of viruses, etc (Figure 3B).

**Gene Prediction and Annotation of Gut Microbiome.** The gene set of intestinal flora is estimated to be about 3,881–5,389 annotated genes that harbored an extensive metabolic repertoire, which is distinct from, but complements the activity of host enzymes in the digestive gland and intestine, including functions essential for food digestion and nutrient absorption. In addition, ammonium (the major nitrogenous waste of the host) assimilation in the meta-pathway of intestinal flora indicate their ability of recycling host's metabolic waste, and other metabolites generated by intestinal microbiota, such as bacteriocins, short-chain fatty acids, and quorum-sensing autoinducers, were essential for intestinal homeostasis (Gao et al., 2009). Here, we discuss the main intestinal flora, particularly bacteria, and genes involved in microbial pathways associated with the metabolism of host-ingested substances, including carbohydrates, proteins, vitamins and minerals, which could be used to expand the nutrient acquisition capacities of the *Alviniconcha* intestine. Genes involved in intestinal microbial hydrolases were extracted from the metagenome, their annotations and expression levels were summarized in Dataset S1, all the annotated information of intestinal flora was in Dataset S2.

**Potential Cross-feeding of Gut Microbiome.** Nondigestible carbohydrates can be converted into critical metabolites such as vitamins and short chain fatty acids (SCFAs) by saccharolytic bacteria. SCFAs produced by the intestinal microbiota affect lipid, glucose, and cholesterol metabolism in animals and humans (Pranal et al., 1996). Moreover, this fermentation results in SCFAs together with the gases $CO_2$ and $H_2$, which can be utilised by acetogens or archaeal microbiota in the gut of *A. marisindica*. For example, acetogens convert $CO_2$ into acetate, methanogenic archaea can generate $CH_4$ using $H_2$ and $CO_2$ from polysaccharide processing, and sulfate-reducing bacteria use $H_2$ for sulfate reduction to produce $H_2S$. Nitrogen is essential for bacterial survival in the intestine and dissimilatory nitrate reduction and ammonium assimilation are found in the intestinal microbial meta-pathway (Figure 5). Genes encoding for ureases are found in the metagenome of intestinal microbes, and urea produced by the host can be hydrolysed to ammonia by these urease-producing bacteria. The ammonia in turn can be used for protein metabolism. The occurrence of urea-nitrogen recycling suggests that the gut microbiome plays an important role in the nitrogen balance of *A. marisindica* (Figure 5). Co-abundance and meta-pathway analyses indicate that the gut microbiome likely play critical ecological roles via bacterial interdependencies and mutual cooperation to maintain intestinal homeostasis and provide the host with symbiont metabolites that serve as host nutrients.

**Gut Microbial Hydrolases.** There are three main types of hydrolases secreted by intestinal flora, including glycoside hydrolases (GHs) which can deconstruct complex carbohydrates (e.g. cellulose, chitin), protease (including aminopeptidases) and esterase (mainly are lipases). In *Alviniconcha marisindica*, digestive exoenzymes produced by the intestinal microbes help the enzymatic digestion of food in intestine (Table S3), α-Amylase was found abundant in the digestive exoenzymes of intestinal flora which breaks down long-chain saccharides, trypsin and Zinc-dependent metalloprotease were major exoproteases which are necessary for protein absorption, GDSL esterases/lipases were the major lipolytic enzymes for carbon source provision. Especially, glycoside hydrolases (GHs) were found abundant in the intestine of *Alviniconcha marisindica* and has important roles aiding the digestion of dietary carbohydrates. Cellulases, xylanase/chitin deacetylase, glucanases, phosphorylase and some other glycoside hydrolases (GHs) were found extensively in Proteobacteria (Dickeya, Xanthomonadales, Cellvibrio), Actinobacteria, Firmicutes, Tenericutes, Chloroflexi, Bacteroidetes (Cytophagia), and some fungi (Eurotiales, Debaryomycetaceae, Chaetomiaceae) to form multi-enzyme complexes that help the host digestion of ingested carbohydrates. The *Alviniconcha marisindica* lack the enzymes to degrade the bulk of dietary fibers, these nondigestible carbohydrates are converted into utilizable and important metabolites by saccharolytic bacteria, such as vitamins and short chain fatty acids (SCFAs). SCFAs were produced as the key products for nutrition provision via intestinal microbial digestion of the dietary carbohydrates (den Besten et al., 2013), like acetate, propionate, and butyrate, which have a healthy and beneficial effect. Previous studies of rats/mice and humans showed SCFAs produced by the intestinal microbiota affect lipid, glucose, and cholesterol metabolism in various tissues (Gao et al., 2009, Todesco et al., 1991), for example, the fatty acid metabolism is regulated by SCFAs in the body including the balance between fatty acid synthesis, fatty acid oxidation, and lipolysis (den Besten et al., 2013); glucose metabolism is beneficially affected by SCFAs via normalizing plasma glucose levels and increasing glucose handling (den Besten et al., 2013). A large part of the SCFAs is used as a source of energy in various animal species, for example, SCFAs contribute ~10% of the daily caloric requirements of humans, ~20–30% for several other omnivorous or herbivorous animals (Bergman, 1990).

Gene encoded the sialate O-acetylesterase was highly expressed in the meta-transcriptome of intestinal flora which indicate sialic acid degradation is active in the intestine of *Alviniconcha marisindica*. Sialic acids (Sia) are prominent outermost carbohydrates of the intestine which are important components of mucus layer (Li et al., 2015). A pathway for the transport and catabolism of Sia was encoded in the metagenome of intestinal flora which indicate sialic acid can be utilized as a carbon source.

**Lactic Acid Bacteria in Gut.** Various lactic acid bacteria were found co-exist in the intestine, i.e. families of 1.29–1.55% of Lactobacillaceae, 0.20–0.43% of Streptococcaceae, 0.25–0.34% of Enterococcaceae, 0.14% of Leuconostocaceae (Leuconostoc), 0.14% of Carnobacteriaceae and 0.12% of Aerococcaceae. Mixed lactic acid bacteria have the potential to prevent pathogens causing intestinal infection and play an important role in the stability of intestinal microbes for maintaining host health (Ren et al., 2018, Pessione, 2012), which exhibit a mutualistic relationship with the host. For example, *Lactobacillus* species found in the gastrointestinal tracts are commonly used as probiotics that confer a health benefit on the host (Walter, 2008), at the family level, Lactobacillaceae and Streptococcaceae became prevalent after probiotic treatment, and microbial community composition was more stable during the period of probiotics treatment (Hemarajata and Versalovic, 2013).

**Summary of Gut Microbial Functional Analysis.** The above mentioned functional analysis of intestinal flora indicate the microbiota has great impact on intestinal nutrient degradation and absorption, host immune system stimulation and help defense against pathogens. The degradative processes in the intestine following the action of bile acid, pancreatic and intestinal enzymes were essential for nutritional homeostasis in the *Alviniconcha holobiont*. Hence, even if the intestinal flora at a lower abundance than the endosymbionts in the gill, they are significant to the host health and survival. In this study, intestinal microbial ecology and its interplay with the host metabolome are critical to nutritional and metabolic demands of *Alviniconcha* holobiont and help to shape unique microenvironments of intestine.

**Supplementary Note 7**

**Genomic Comparison of the Endosymbiont.** The gill endosymbiont of the Wocan *Alviniconcha marisindica* was compared with the endosymbiont of *Lamellibrachia* tubeworm (Patra et al., 2016), the epibiont of the giant tubeworm *Riftia pachyptila* (Giovanneli et al., 2016), and two free-living Campylobacterota (Inagaki et al., 2004; Nakagawa et al., 2007) from deep-sea hot vents. Whole-genome ANI of orthologous gene pairs shared between two microbial genomes was calculated using fastANI (Jain and Rodriguez-R, 2018). The orthologous groups (OGs) from the above five Campylobacterotal genomes were detected using Proteinortho v5.16b (Lechner et al., 2011) (BLAST threshold E = $1 \times 10^{-10}$), a total of 1,053 shared single copy orthologs were identified. PCA analysis on the shared orthologous proteins was performed using Jalview (Waterhouse et al., 2009), BLOSUM62 model was used to calculate the similarity scores between each pair of sequences and form a matrix, then the components were generated and visualized using BioVinci (Bioturing, San Diego, CA, US) (Figure S22).

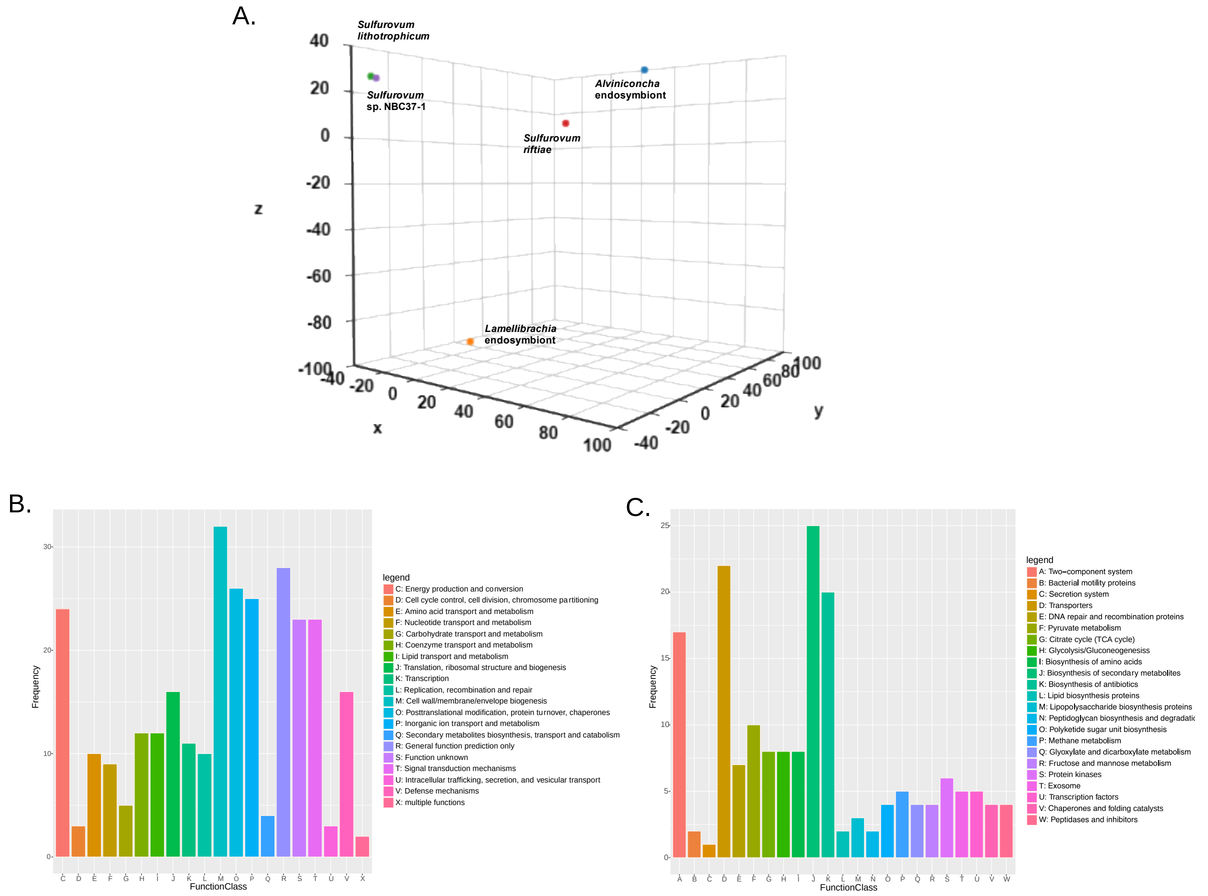

**Figure S22. Genomic comparison analysis across representatives of Campylobacterota.** (**A**) PCA analysis on the orthologous proteins in 1,053 OGs of five Campylobacterota under the BLOSUM62 model. The reduced orthologous genes of the endosymbiont of *Alviniconcha*

*marisindica* are classified into different functional categories based on (**B**) COG and (**C**) KEGG annotations.

**Reduced Genes of *A. marisindica* Endosymbiont.** Based on PCA analysis, the five Campylobacterota were separated according to habitat type (Figure S22A). The free-living Campylobacterota were clustered, the endosymbiont of shallow-water seep vestimentiferan, *Lamellibrachia satsuma*, the endosymbiont of deep-sea vent snail, *Alviniconcha marisindica*, and the epibiont of deep-sea vent vestimentiferan, *Riftia pachyptila* were well separated from each other. The horizontal transmitted endosymbionts and epibionts of seep- or vent-animals likely come from the environmental free-living population. The animal hosts have the potential to adopt endosymbionts that optimally adapt to both ambient seawater and host intracellular environment. Therefore, different habitats may have driven the molecular adaptation in the above symbionts. Comparison of complete genome sequences within clades have revealed that the endosymbiont of *A. marisindica* exhibit the small genome with fewer coding sequences. As a result, 352 orthologue genes were found lost in this endosymbiont. The functional composition of the loss genes included genes involved in [M] cell wall/membrane/envelope biogenesis (e.g. capsular polysaccharides), [O] posttranslational modification, protein turnover, chaperons, [P] inorganic ion transport and metabolism, [C] energy production and conversion, [T] signal transduction metabolism and [V] defensive mechanisms account for the main part, in which 174 reduced genes annotated via KEGG and mainly distributed in two-component system (e.g. chemotaxis proteins and response regulators), DNA repair and recombination (e.g. DNA polymerase polX), carbohydrate metabolism (e.g. part of pyruvate metabolism), biosynthesis of amino acids, ABC transporters and other element transporters (Figure S22 B and C). The endosymbiont doesn't have many non-essential metabolic genes and part of central carbohydrate metabolism, for example, partial Citrate cycle which is one of the optional from pyruvate to aceyl-coA (genes *ace* and *DLAT*) was missing.

**Loss-of-Function Mutations.** The effects of variation in shared orthologous proteins of the endosymbiont of *Alviniconcha marisindica* and four Campylobacterotal references were revealed by the DBS method. Functional genetic changes in conserved domains within shared orthologous protein sequences showed the number of genes, classified by functions, with loss-of-function mutations in the endosymbiont of *Alviniconcha marisindica* were much less than the other four Campylobacterotal references, only 10–13 loss-of-function mutated orthologous proteins were found (Figure 4A). Comparing with the two free-living and one epibiotic Campylobacterota, several bacterial surface proteins were found lost their functions in the endosymbiont of *Alviniconcha marisindica*, including outer membrane protein/protective bacterial surface antigen, OMA87 (PF03279.13), bacterial lipid A biosynthesis acyltransferase (PF03279.13), putative membrane protein insertion efficiency factor (PF01809.18) and Sel1 repeats (SLRs) (PF08238.12) which regarded as bacterial virulence determinants and may also be recognized by host cells (Newton et al., 2007; Emiola et al., Tabatabai, 2008). Furthermore, NnrS protein (PF05940.12) was found lost its function in the endosymbiont of *Alviniconcha marisindica* compared to the endosymbiont of *Lamellibrachia satsuma*. NnrS is particularly important for protecting bacterial resistance to nitrosative stress under anaerobic conditions (Stern et al., 2013), though protein sequences of this enzyme was found mutated and may lost its function in the endosymbiont of *Alviniconcha marisindica*, genes encoded nitric oxide synthase-interacting protein and NADPH--cytochrome P450 reductase (*POR*) which participated in nitric oxide catabolic process were found expressed in *Alviniconcha* host cells to alleviate this stress. The loss-of-function orthologous genes from the endosymbiont of *Alviniconcha marisindica* and other four Campylobacterotal species are listed in Dataset S3.

**Supplementary Note 8**

**Heterogeneous Genomes of the Endosymbiont Populations.** A total of 20 endosymbiont isolates were obtained from the posterior and anterior parts of the gill of 10 *A. marisindica* individuals. A pipeline BactSNP (Yoshimura et al., 2019) was used to identify single-nucleotide polymorphisms (SNPs) between isolates, this pipeline was capable of highly accurate and sensitive SNP calling in a single step even when target isolates are closely related. In this study, we used the genome of isolate '7-B' as reference genome, SNPs were called among isolates. Pseudo genomes of isolates were obtained with all contigs in concatenated into one sequence for each isolate, and then input to phylogeny analysis. In addition to the genomic ANI results, a phylogenetic tree using SNPs was constructed to represent reliable evolutionary relationships among 20 endosymbiont isolates. The distance between 2 isolates is the sum of the length of all branches connecting them, and we defined the isolates clustered in the same clade at different levels on the phylogenetic tree with distance less than 0.00198 as the same strains. In this case, five strains were identified from 20 endosymbiont isolates. The endosymbionts under stressful conditions can bestow a selective advantage via increasing genetic variation. In addition, genetic variation of the endosymbiont allows it to establish themselves in their chosen host and also allows them to resist host's subsequent innate immunity like many pathogenic bacteria (Robertson and Meyer, 1992). Interestingly, we found that the endosymbiont isolates were not clustered according to their host and the strains randomly distributed in different host individuals. Based on the semi-endosymbiotic model of *A. marisindica*, we supposed the *A. marisindica* had the ability to exchange or re-acquire gill endosymbionts from the environment, and had potential to obtain more flexible and diverse endosymbiotic bacteria than many other holobionts with true endosymbiosis.

**Supplementary Note 9**

**DEGs of Different Tissues/Organs.** The *Alviniconcha* genome encoded complete metabolic pathways within lysosomes, endosomes and phagosomes which serving as main digestive compartment of host cells. Especially, genes encoded large various proteases, bile salt-activated lipase and glycosyl hydrolases were highly expressed in the intestine, genes encoded gastric intrinsic factor and bile acid transporter were highly in the digestive gland, which strongly support the active digestion of macromolecular nutrients in the intestine. The annotation of highly expressed genes in the gill, intestine, digestive gland, and mantle of *A. marisindica* were shown respectively in Dataset S4 and Figure 23. Large amount of highly expressed genes in the intestine showed it has functional specializations that responsible for the effective and regulated nutrients transport. The number of differentially expressed genes in the gill and intestine, with their biological coefficient of variation (BCV) were shown respectively in the Figure S24. Based on the DEGs (selective highly expressed genes), functional enrichments of GO terms/KEGG pathways were also identified in the gill and intestine of *A. marisindica*. In the intestine, genes involved in vacuole, hydrolases activity, organic hydroxy compound metabolic process, and transmembrane transport were enriched, indicating its active nutrients digestion and absorption (Figure S12A). In the gill, leukocyte differentiation, myeloid leukocyte activation, MyD88-independent toll-like receptor signaling pathway, and TNF signaling which contribute to host immune system were enriched, indicating the ability of resisting invasions and controlling symbionts of the gill. Enrichment of lipid biosynthetic process, regulation of protein synthesis and intracellular transport in the gill showed its active nutrients metabolism (Figure S12B).

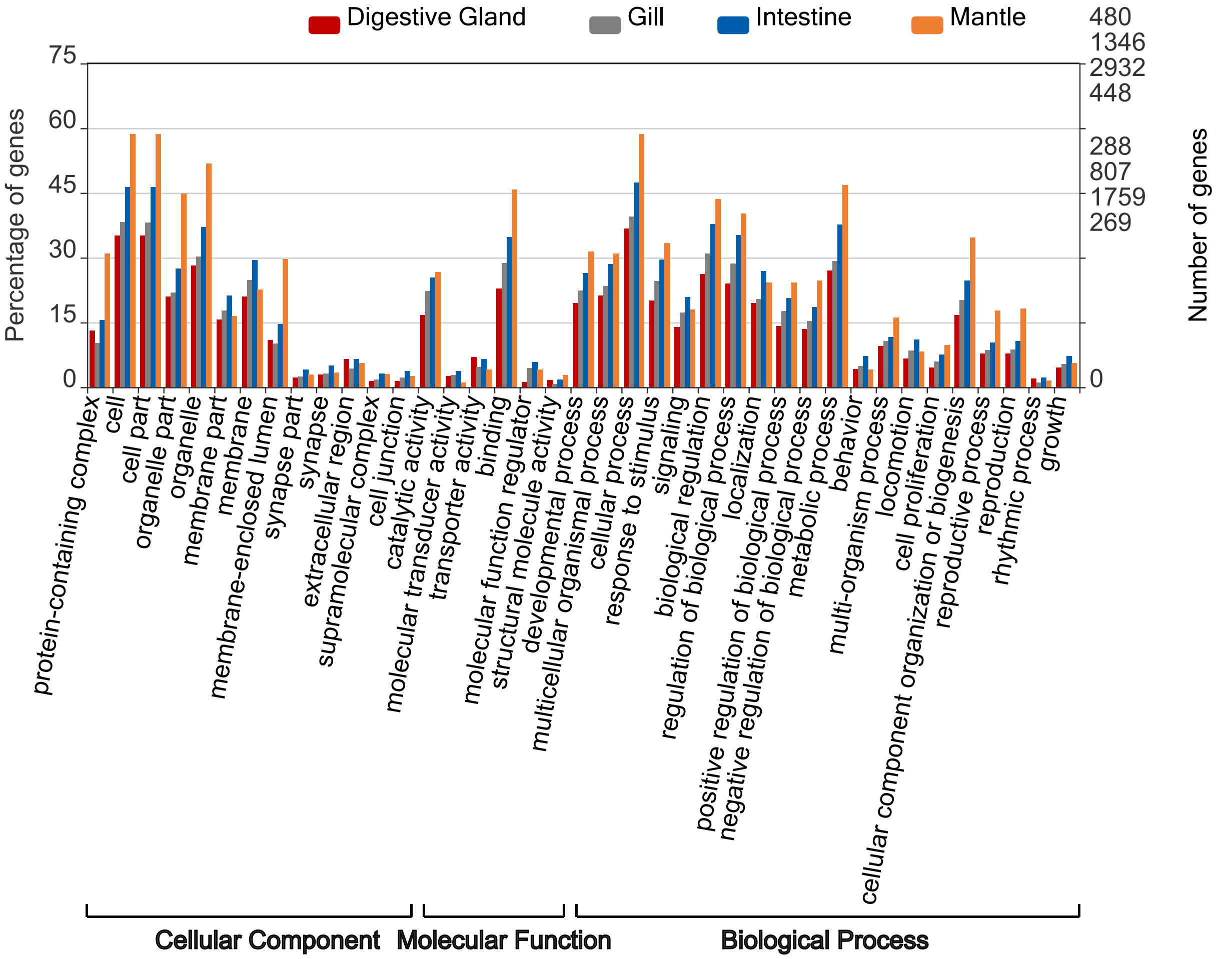

**Figure S23. Gene Ontology (GO) functional annotations of highly expressed genes in the digestive gland, foot, gill, and intestine.** X-axis represents the default GO terms selected automatically by WEGO online tool, the colour of bars represents different tissues of *A. marisindica* (red – digestive gland, grey – gill, blue – intestine, orange - mantle), the length of bars represents percentage and number of transcripts in different GO functional classes. GO annotation provides three main functional categories (cellular component, molecular function, and biological process).

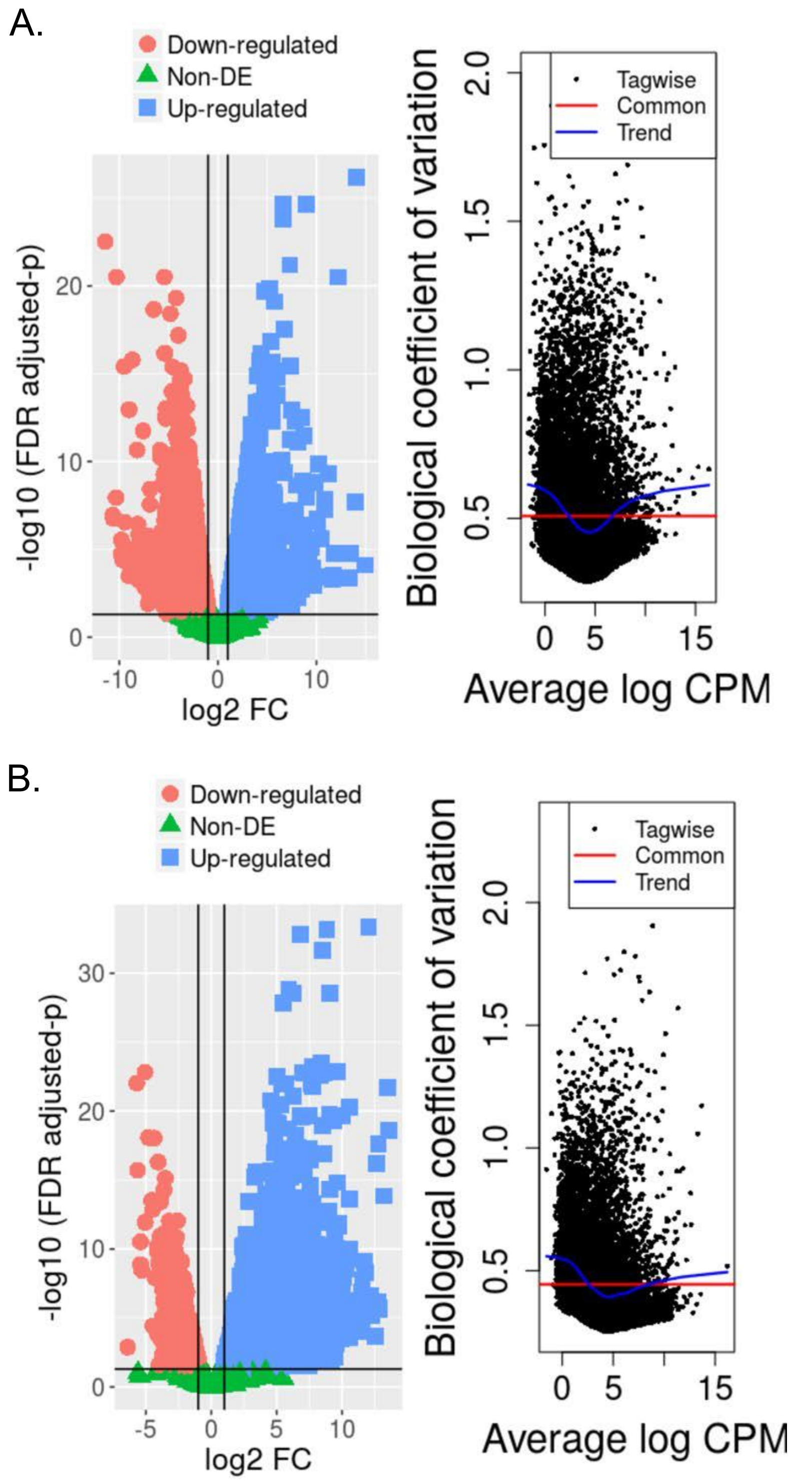

**Figure S24. Differentially expressed genes (DEGs) in the intestine and gill of *Alviniconcha marisindica*.** Volcano plot and biological coefficient of variation (BCV) plot of differentially expressed genes in the (**A**) gill and (**B**) intestine are identified by DESeq2 analysis. The log10 (FDR corrected p-values) are plotted against the log2 (FC) in gene expression. Upregulated genes by twofold or more and with a FDR corrected p-value < 0.05 are marked as blue dots, while down-regulated genes (FRD <= 2 with P < 0.05) are in red colour.

**Substrates Transport.** Previous study showed the substrates for chemosynthesis including sulfide and oxygen are obtained from ambient seawater through hemoglobin and hemocyanin in the gill of *Alviniconcha hessleri* (Wittenberg and Stein, 1995). However, the genome of *A. marisindica* does not have genes encoding hemoglobin subunits where this genome contains the genes encoding hemocyanin, globin-like (*glob1*, *NGB*), myoglobin-like (I*DO2*) and globin C, coelomic-like that are responsible for substrates transportation. Importantly, the hemocyanin genes showed an extremely high expression (top 10 in the intestine) in the intestine (Figure S25). Accordingly, a large volume of blue hemolymph was observed. In this study, we supposed that hemocyanin was not only oxygen transporter as previous studies suggested (Wittenberg and Stein, 1995). It also capable to bind and transport sulfur compounds to the symbionts.

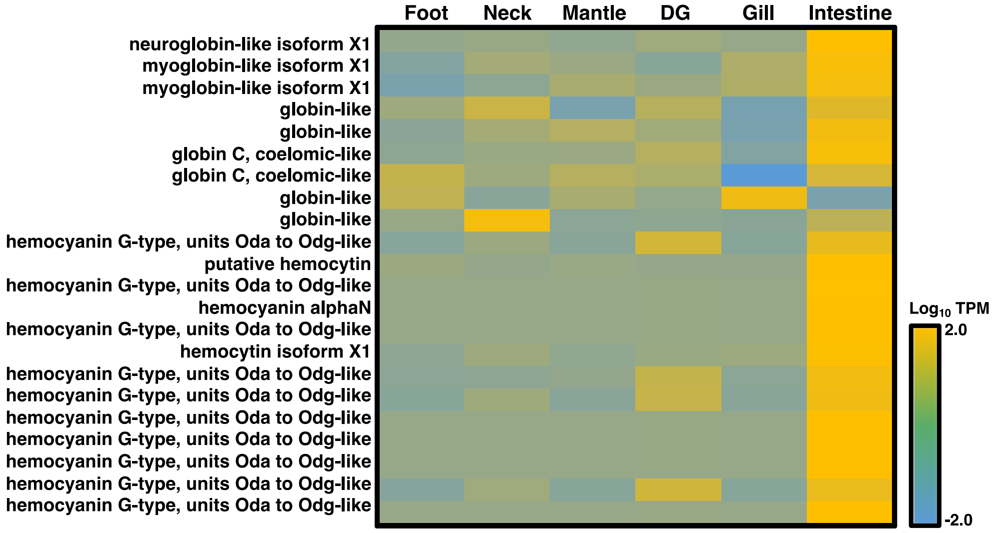

**Figure S25. Transcriptional activity of genes participating in the globin of Wocan *Alviniconcha marisindica* for substrates transportation.** Heat map of transcriptional activity of genes that encode neuroglobin, myoglobin, globin-like, and hemocyanin in different tissues, including foot, neck, mantle, digestive gland (DG), gill and intestine. Each grid in the heat map represents an identified gene in the respective sample. The colour represents the gene expression level (based on normalized TPM values). The annotated gene names are listed on the sides.

**Digestion of Symbionts and Nutrients Transportation.** Functional enzyme distribution of *A. marisindica* genome shows that the host contains numerous genes responsible for key hydrolases, and the expression of these genes is higher in the gill and intestine than in other tissues/organs (Figure S11A), especially intestine showing a highly active digestive system. The presence of none specific transporters of essential nutrients in gill endosymbiont suggests nutrients are released via leakage or digestion by host (Yang et al., 2020, Newton et al., 2007). For example, lysosomal cathepsins, non-lysosomal calpain and caspases are highly expressed in the gill. However, except of membrane-lytic enzymes such as aminopeptidases showing high expression, proteinases and glycosyl hydrolases are not more active in the gill than in other tissues. The active T2SS of the endosymbiont also has the ability to translocate intracellular proteins into outside. In this case, the endosymbiont releases the nutrients not only through being digested but also using T2SS to export. Genes that are involved in apoptosis regulation are expressed highly in the gill that harbors highly expressed caspases genes (Figure S11A) responsible for programmed cell death. Since the cell initiates intracellular apoptotic signaling in response to a stress, such as chemical cues in the environment, nutrient deprivation, viral infection or other immune stimuli (Kemp, 2017), it is reasonable to conclude that when endosymbionts proliferate quickly, the cytosolic concentration in host cells will induce the regulation of cell apoptosis. Hence, the density of endosymbionts shall be controlled by host bactericidal activity and apoptosis, which are triggered by limited space and nutrient deprivation in host cell. The cell apoptosis in the gill has the potential to help host cells obtain nutrients.

**Dataset S1 (separate file).** The annotation and expression levels of genes involved in encoding hydrolases in intestinal flora of three *A. marisindica* individuals.

**Dataset S2 (separate file).** The annotation of genes predicted from the metagenome of intestinal flora from three snail individuals.

**Dataset S3 (separate file).** The loss-of-function orthologous genes from the endosymbionts of *Alviniconcha marisindica* and four campylobacterotal references through their pair-wise genomic comparisons based on a profile-based method.

**Dataset S4 (separate file).** The annotation of highly expressed genes (Fold Change>2) in different tissues (gill, intestine, digestive gland, and mantle) of *A. marisindica*.
